# N-Aryl Pyrido Cyanine derivatives: nuclear and organelle DNA markers for two-photon and super-resolution imaging

**DOI:** 10.1101/2020.05.26.116335

**Authors:** Kakishi Uno, Nagisa Sugimoto, Yoshikatsu Sato

## Abstract

Live cell imaging using DNA-binding fluorescent probes is an essential molecular tool in various biological and biomedical fields. The major challenges in currently used DNA probes are to avoid UV light photo-excitation with high DNA selectivity and cell-permeability and are the availability of the cutting-edge imaging techniques such as a super-resolution microscopy. Herein we report new orange to red fluorogenic DNA probes having N-aryl pyrido cyanine (PC) moiety as a basic skeleton. Their DNA selectivity and cell-permeabilities are so high that organelle DNA as well as nuclear DNA can be clearly stained in various cell types and plant tissues with wash-free manner. PC dyes are also compatible with a stimulated emission depletion fluorescent lifetime imaging microscopy (STED-FLIM) for super-resolution imaging as well as two-photon microscopy for deep tissue imaging, should release the utilization limitation of synthetic DNA probes.

## Introduction

Synthetic fluorescent dyes for DNA stains are essential tool in current biological and biomedical science. In addition to gel electrophoresis^1,2^, polymerase chain reaction (PCR) in molecular biology^3–5^, and flow cytometry^6–7^, they are also greatly used in cell biological study such as for visualizing nuclear and organelle DNA^8,9^, cell proliferation analysis^10,11^, and diagnosis of virus infection^12^. The ideal properties for synthetic nucleus marker are i) high selectivity of DNA over RNA, ii) the applicability to long wavelength photo-excitation (>532nm)^13^, and iii) the ability to stain nucleus of diverse living cells and tissues. Such nucleus markers to achieve these requirements, however, has not developed yet in spite of tremendous efforts, although it has long history of developing DNA staining dye.

Current nucleus markers heavily rely on blue emitting Hoechst 33342^14,15^ despite the needs of photo-toxic UV light for photo-excitation^16^. To overcome this drawback, Hoechst tagging strategy, which fluorescent dye excited by visible wavelength was fused to Hoechst through linker, has been reported recently^17,18^. Especially, silicon-rhodamine (Sir)-Hoechst, which fused a far-red emissive Sir to Hoechst 33342, stained nucleus with no significant photo-toxicity and showed compatibility with stimulated emission depletion (STED) nanoscopy^18,19^. In exchange for these preferable properties, however, the high DNA-binding strength of Hoechst has been lost by fusing Sir and the increase of molecular weight is inevitable in the Hoechst tagging strategy which are considered disadvantage for cell permeability.

On the other hand, unsymmetrical cyanine fluorescent dyes, such as SYBR-Green^20^, Pico-Green^21^, TO-PRO-1/TOTO^22, 23^, TO-PRO-3/TOTO-3^23–25^, are also the most widely used DNA/nucleus markers. The most remarkable properties represent their high fluorescence jump upon binding nucleic acid and availability of visible wavelength light for their excitation^22–24,26^. Out of all of unsymmetrical cyanine dyes, SYBR-Green and Pico-Green have a favorable character unlike Hoechst that can stain mitochondrial DNA (mt-DNA)^9,27–30^ as well as nuclear DNA^31,32^ in living cells. Unfortunately, however, these dyes are mostly excited with photo-toxic short wavelength laser such as 488 nm^13^ and do not have high DNA selectivity over RNA^25,33^. The excitation/emission wavelength in these monomethine cyanine dyes can be easily red-shifted by extending π-conjugation to synthesize trimethine dyes. The thus obtained red-shifted dyes, however, showed low fluorescence jump upon DNA binding and needed high concentration to stain cell nucleus in fixed cells^34^. So far, fluorescent DNA marker which fulfills above all requirements is still awaited.

To this end, in the present study, we synthesized unprecedented series of symmetrical cyanine DNA markers based on N-aryl pyrido cyanine (PC). The produced novel N-aryl PC dyes provided expanded options of excitation wavelength with the exceptional DNA/RNA selectivity. In addition to these great properties, N-aryl PC dyes has an outstanding cell permeability to clearly stain nuclear and mt-DNA in concentration dependent manner in various cell types. Therefore, here we demonstrated their versatile abilities for various optical microscopies such as two-photon excitation microscopy (2PEM), FLIM, and STED-FLIM nanoscopy.

## Results

### Molecular design and in vitro characterization of N-aryl PC derivatives

To achieve long absorption DNA selective markers with a small-molecule fluorophore, we envisioned that PC derivatives could give an advantage for making longer wavelength excitable dyes since PC has an extended π-conjugation with a monomethine cyanine unit^35,36^. Firstly, we designed and synthesized N-Phenyl Pyrido Cyanine (**PC1**) (**Fig. 1a**). UV-Vis and fluorescence spectra measurements revealed that the maximum absorption wavelength of free **PC1** was 510 nm and it was red-shifted to 532 nm when binding upon DNA (**Table 1**, **Fig. S1a**), which are effectively longer than those in previously reported unsymmetrical monomethine dyes (460-520 nm)^20–23^. When fluorogenic properties (I^dsDNA^/I^free^ or I^RNA^/I^free^) of **PC1** on binding to nucleic acids (DNA and RNA) were evaluated, it was extremely high and jumped 1600-fold upon binding DNA (**Table 1, Fig. S2a**). On the other hand, favorably for DNA selective property, the fluorogenicity (I^RNA^/I^free^) was only up to 110-fold upon binding to RNA. Actually, the value of DNA/RNA selectivity (I^dsDNA^/^RNA^) of **PC1**, which calculated by dividing I^dsDNA^ by I^RNA^, was extremely higher than that of most popular commercialized nucleus probes, Hoechst 33342^14^ (**Fig. S2i**) and Pico-Green^21^ (**Fig. S2j**). To know the nucleotide sequence specificity of **PC1**, we performed fluorescence titration with three hairpin oligonuclotides (^AATT^DNA, ^CGCG^DNA, ^AAUU^RNA)^18,37^ and found that **PC1** preferentially binds to ^AATT^DNA, while no noticeable bindings to ^GCGC^DNA and ^AUAU^RNA (**Fig. 1c, Fig. S3**). These results suggest that **PC1** specifically bind to AT base-pair of nucleic acids. For tuning toward further optical red-shift, we also synthesized N-aryl PC dye derivatives by replacing N-phenyl group to electron donating N-aryl groups^38^ such as anisole (**PC2**), N,N-dimethylaniline (**PC3**), and N,N-diethylaniline (**PC4**) (**Fig. 1d**). It should be noted that despite the structural modifications, exceptional DNA/RNA selectivity and high DNA-based fluorogenic properties are not impaired by these structural modifications (**Table1**, **Fig. S1 and S2**). Furthermore, we synthesized additional four trimethine PC dyes which possess N-Phenyl (**PC5**), N,N-dimethylaniline (**PC6**), N,N-diethylaniline (**PC7**) and 1-methyl-4-phenylpiperazine (**PC8**) which possess extended the π-conjugation of methylene chain (**Fig. 1d**). In these deep-red and far-red emissive dyes, comparable DNA/RNA selectivity were remained, although the fluorogenicity (I^dsDNA^/I^free^) were lower than that in monomethine PC dyes. Thus, we succeeded to synthesize the new series of PC based symmetrical cyanine DNA probes providing researchers with expanded options of excitation wavelength from 500-700 nm (**Fig. 1e-f**).

**Figure 1.**
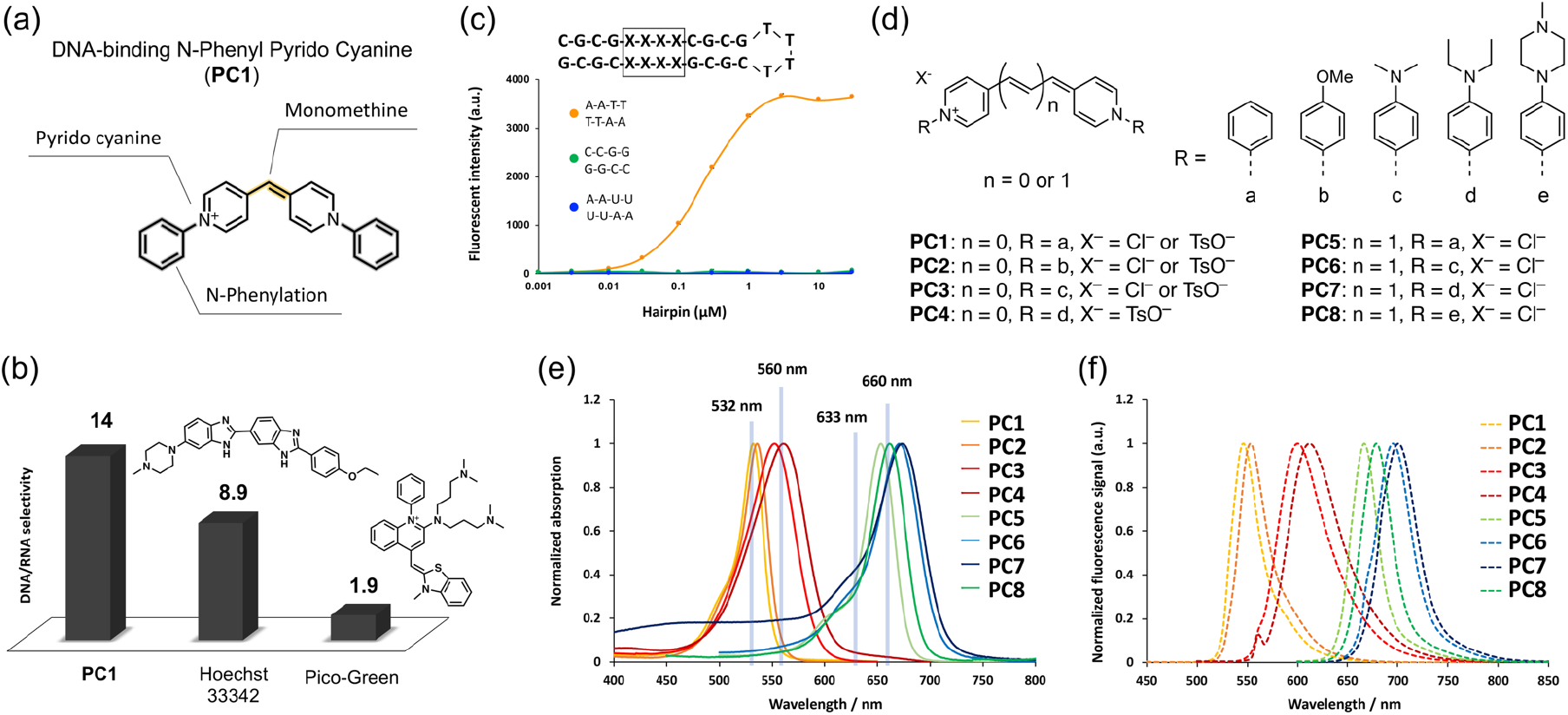
Molecular design and in vitro characterization of N-aryl PC derivatives. a) The structure and structural components of **PC1**, b) The comparison of DNA/RNA selectivity of **PC1**, Hoechst 33342, and Pico-Green, c) The titration curve of 100 nM **PC1** with various concentration of hair-pin oligonucleotides. The error bars indicate means ± s.d. of three independent replicates. d) The general structure of PC dyes and their substituent patters with corresponding compound names; the unit of methylene length and N-aryl groups are represented as “n” and “R” respectively. The normalized absorption (e) and fluorescence spectra (f) of all PC dyes when complexed with calf thymus double stranded DNA (dsDNA) in tris-EDTA buffer solution. (pH= 8.0); see details in Fig. S1 and Fig. S2.

**Table 1.** Photophysical properties of all synthesized N-aryl PC derivatives.

| Name | $\lambda_{\text{abs}}^{\text{dsDNA}}/\text{nm}$ <sup>(a)</sup><br>($\lambda_{\text{abs}}^{\text{free}}/\text{nm}$) <sup>(b)</sup> | $\epsilon^{\text{dsDNA}}/10^4$<br>$\text{M}^{-1}\text{cm}^{-1}$ <sup>(c)</sup> | $\lambda_{\text{em}}^{\text{dsDNA}}/\text{nm}$ <sup>(d)</sup><br>($\lambda_{\text{em}}^{\text{free}}/\text{nm}$) <sup>(e)</sup> | $\Phi_{\text{F}}^{\text{dsDNA(f)}}$<br>$\Phi_{\text{F}}^{\text{free (g)}}$ | $I^{\text{dsDNA}}$<br>$/I^{\text{free (h)}}$ | $I^{\text{RNA}}$<br>$/I^{\text{free (i)}}$ | $I^{\text{dsDNA}}$<br>$/I^{\text{RNA (j)}}$ | $\tau^{\text{dsDNA}}$<br>$/\text{ns}$ <sup>(k)</sup> |
| --- | --- | --- | --- | --- | --- | --- | --- | --- |
| PC1 | 532 (510) | 14 | 546 (535) | 0.09<br>(0.0004) | 1600 | 110 | 14 | 0.62 |
| PC2 | 536 (512) | 13 | 553 (537) | 0.22<br>(0.0006) | 2500 | 170 | 15 | 0.86 |
| PC3 | 552 (520) | 8.6 | 600 (592) | 0.42<br>(0.0018) | 3900 | 130 | 31 | 1.5 |
| PC4 | 561 (524) | 7.7 | 612 (605) | 0.44<br>(0.0026) | 1700 | 53 | 32 | n.d. |
| PC5 | 654 (630) | 15 | 666 (652) | 0.4<br>(0.039) | 140 | 25 | 5.7 | 2.7 |
| PC6 | 671 (639) | 10 | 695 (666) | 0.26<br>(0.038) | 54 | 2.1 | 26 | 1.7 |
| PC7 | 674 (642) | 9.6 | 700 (683) | 0.20<br>(0.038) | 57 | 2.3 | 25 | 1.8 |
| PC8 | 662 (637) | 9.7 | 678 (666) | 0.33<br>(0.075) | 39 | 20 | 2 | 2.3 |

Tris-EDTA buffer solution (TE buffer, pH 8.0) was used for all optical measurements. <sup>(a)</sup> Maximum absorption wavelength of dye-dsDNA complexes. <sup>(b)</sup> Maximum absorption wavelength of dyes in their free states. <sup>(c)</sup> Molar absorption coefficient of dye-dsDNA complexes. <sup>(d)</sup> Maximum fluorescence wavelength of dye-dsDNA complexes and <sup>(e)</sup> maximum fluorescence wavelength of free dyes. <sup>(f)</sup> Fluorescence quantum yields of dye-dsDNA complexes and <sup>(g)</sup> fluorescence quantum yield in their free states. <sup>(h)(i)</sup> Ratio of fluorescence increases of each dye upon binding to dsDNA and RNA; see the SI for the detailed changes in fluorescence spectra. <sup>(j)</sup> The value of calculated DNA/RNA selectivity (see main text). <sup>(k)</sup> Fluorescence lifetime of dye-dsDNA complexes.

### Applicability of N-aryl PC dyes in various living-cell types

To test the utility of N-aryl PC dyes for nucleus marker, we stained HeLa cells with all synthesized PC dyes at 1 μM concentration and found that they are capable of labeling nuclear DNA by confocal laser scanning microscopy (CLSM) (**Fig. S4**). As representative case study, we also confirmed that **PC1** and **PC3** specifically bind AT pairs in nucleus by co-staining with Hoechst 33342 which is well known to bind minor groove of AT-rich sequences^37^ (**Fig. S5**). Consequently, these N-aryl PC dyes are excluded from nucleus by Hoechst 33342 in dose dependent manner. These results indicate that they scramble for AT pairs sequences with Hoechst and are consistent with in vitro study using hairpin oligonucleotides (**Fig. 1c, Fig. S3**). The talented ability of N-aryl PC dyes for selective DNA stain without washing process was obvious when compared with commercially available cell permeable SYTO dyes, which have similar excitation and emission spectrum. **PC1** and **PC3** clearly stain nucleus and chromosome with no substantial background from cytosol, whereas SYTO dyes failed to stain nucleus even at higher concentration and they rather stained cytosol and nucleolus which have a large amount of RNA (**Fig. 2a, Fig. S6**). Moreover, cell proliferation analysis as well as its time lapse imaging revealed that **PC1** and **PC3** showed no substantial cytotoxicity and photo-toxicity compared with those of SYTO dyes (**Fig. 2b**). The applicability of **PC1** and **PC3** was also investigated to other mammalian cell types (U-2OS, C6, NIH3T3) and found that they could uniformly stain nuclear DNA without any reagent for facilitating cell permeability (**Fig. 2c**). Considering that SiR-Hoechst needs voltage-dependent calcium channel inhibitor, verapamil, for homogeneous staining of nuclear DNA in U-2OS cells^18^, these results indicate that cell permeability of N-aryl PC dyes is substantially higher than that of SiR-Hoechst. Then, we investigated the applicability of N-aryl PC dyes to stain nucleus in plant tissue which are composed of cells stacking with thick cell wall in layers, and succeeded in this challenge using Arabidopsis leaf and root tissues (**Fig. 2d**). In addition to the epidermal cells including stomata, mesophyll cells underneath epidermis were perfectly stained in the nucleus in leaf tissue without the washing process (**Movie S1** and **S2**). On the other hand, time-lapse analysis revealed that root hairs as well as main root tissue grew normally and were also clearly stained in their nucleus by **PC1** (**Movie S3**). We next employed 2PEM excited with 1000 nm to investigate the dye penetration in the main root. Consequently, 2PEM enabled to observe whole nuclei of root tip region by overall cross sections (~100 μm) while CLSM limited to visualize only the half cross sections (~50 μm) (**Fig. 2e**, **Movie S4**). Furthermore, we did not detect any cytotoxicity and photo-toxicity during the study of plant cells stained by PC dyes and cell division with root growth was frequently observed by 2PEM time-lapse imaging (**Fig. 2f, Movie S5**). These results indicate that the PC dyes penetrated deep into the root layer with no apparent toxicity and are very suitable for 2PEM possibly because of the symmetrical donor-acceptor-donor molecule^39^.

**Figure 2.**
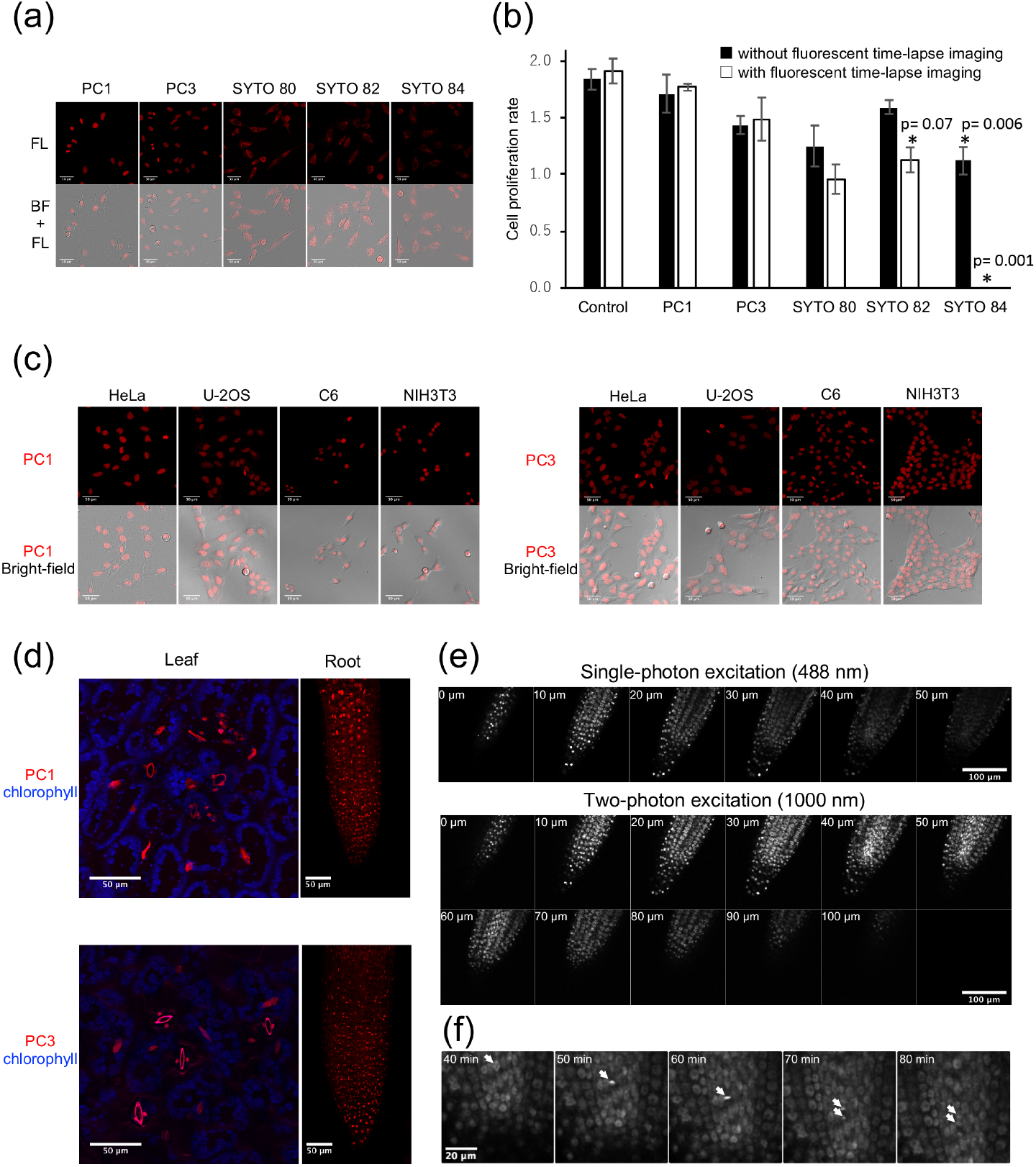
Live cell imaging with PC1 and PC3. (a) Live cell fluorescent microscopy with **PC1, PC3** and commercialized red fluorescent DNA dyes. HeLa cells were stained with each dye at 100 nM. Images are maximum z-projections of total planes (1-μm intervals) (b) Quantification of cell proliferation rate of cells stained with DNA labeling dyes. HeLa cells were stained with each dye at 30 nM and observed every 5 min with z-sectioning (6 frames at 3 μm steps) for 24 h. The proliferation rate was quantified as fold changes based on the number of cells between the first frame (0 h) and the last frame (24 h) of bright-field images. Black bars indicate the results from only bright-field time-lapse imaging without fluorescent time-lapse imaging and white bars indicate the results from both bright-field and fluorescent time-lapse imaging. Error bar shows mean ± s.d. from three independent biological replicates (>26 cells per replicate). Statistical significance (p-value <0.01) of difference from control condition was examined bytwo-sided student t-test. (c) Live cell fluorescent images of different culture cell types with 500 nM **PC1** and **PC3**. The images are maximum z-projections of total planes (1 μm intervals) (d) Live cell fluorescent images of Arabidopsis leaf and root cells with 1 μM **PC1** and **PC3**. The images are maximum z-projections of total planes (1.1 μm intervals) (e) Comparison of imaging penetration for single and two-photon excitation microscopy in Arabidopsis root tip stained with 1 μM **PC1**. Images were shown every 10 μm steps from z-sectioning images at 1 μm interval. (f) Time-lapse observation by two-photon microscopy excited with 1000 nm in Arabidopsis root stained with 5 μM **PC1**. Root tip was observed every 2 min with z-sectioning (50 frames at 2 μm steps).

### Discrimination between nuclear DNA and organellar DNA with fluorescence lifetime of N-aryl PC dyes

During the exploration of the minimized concentration for N-aryl PC dyes, we found that **PC1** obviously stained nucleus with very low cytoplasmic background even at only 10 nM (**Fig. 3a**). In contrast, many cytoplasmic spots as well as nucleus were also observed at 1 nM and the signal from the spots instead from nucleus became predominant by further dilution at 100 pM (**Fig. 3b** and **3c**). In our previous study, the similar changes of staining pattern were observed using SYBR-Green which stained not only nucleus but mitochondrial nucleoids (mt-nucleoids), the core complexes of mitochondrial DNA (mt-DNA) replication and transcription^30^. We also confirmed that the fluorescence spots resided in mitochondria by co-staining of **PC1** and MitoTracker (**Fig. S7**). These results indicate that **PC1** enables to clearly stain mt-DNA as well as nuclear DNA in dose dependent manner. Furthermore, we found that the fluorescent lifetime of **PC1** in nucleus (~ 1.1 ns) was substantially longer than that in mt-nucleoids (~ 0.5 ns) and they were clearly discriminated with different pseudo colors using FLIM combined with phasor plot analysis in various cell types (**Fig. 3d-l**). Similar results were obtained in the **PC3** stained cells (**Fig. S8**). Furthermore, plant cells stained with **PC1** were also applied to extend the concept of FLIM based separation. We envisioned that **PC1** could also distinguish chloroplast DNA (ch-DNA), another organelle having own genome, from mt-DNA and nuclear DNA if the fluorescent lifetime of **PC1** in chloroplasts was quite different from that in mt-DNA and nuclear DNA. Then, we employed FLIM analysis of **PC1** in living stomata of Arabidopsis (**Fig. 3k, Movie S6**). Consequently, the **PC1** in chloroplasts (yellow) had an intermediate fluorescence lifetime (~0.9 ns) compared with that in mitochondria (cyan) and in nucleus (red) and they could be displayed with different pseudo colors. Therefore, FLIM analysis stood out the talented ability of PC dyes and we successfully demonstrated that **PC1** achieved to discern all three DNA storages (nucleus, mitochondria, plastid) in eukaryote without any help of other fluorescent probes.

**Figure 3.**
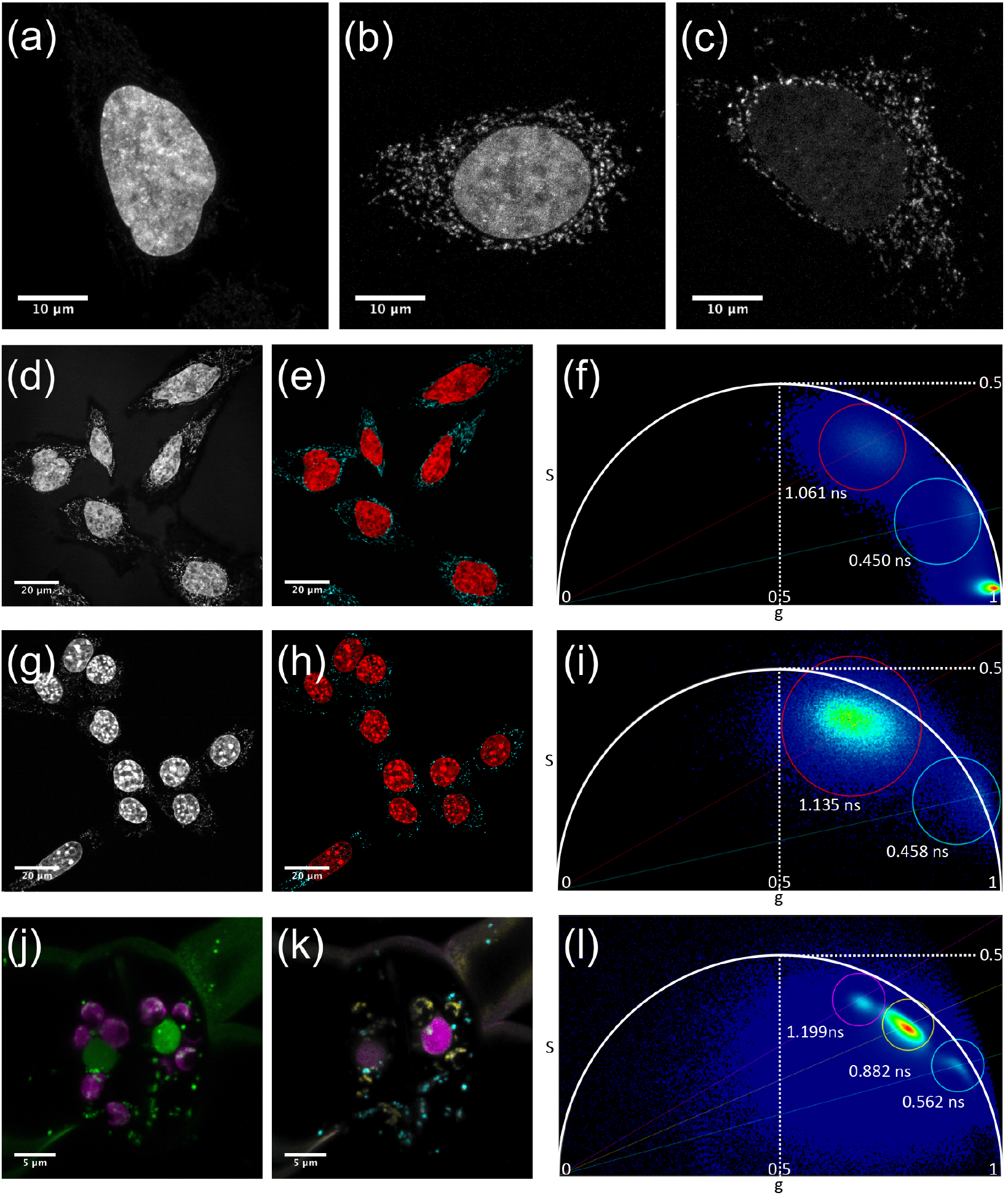
Discrimination between nuclear DNA and mt-DNA with fluorescence lifetime of PC1. **(a-c)** Concentration dependence of staining pattern with **PC1**. (a) 10 nM, (b) 1 nM, (c) 100 pM. The images are maximum z-projections of total planes (0.3 μm intervals). **(d-l)** Fluorescent intensity images (d, g, j) and FLIM based separation images of nuclear DNA and mitochondrial DNA (e, h, k) by phasor plot analysis (f, i, l). The pseudo colors of (e, h, k) is correspond to the colors of circles in (f, i, l). The nuclear DNA, mt-DNA, and ch-DNA are shown in red, cyan, and yellow, respectively. HeLa cells (d-f) and NIH3T3 (g-i) were stained with 1 nM and 10 nM **PC1**, respectively and the fluorescent spectrum were collected between 540-650 nm excited at 532 nm. Stomata in Arabidopsis leaf cells was stained with 300 nM **PC1** and excited at 532 nm. The fluorescent spectrum of **PC1** and chlorophyll autofluorescence were collected between 540-620 nm and 680-700 nm shown in green and magenta in (k), respectively.

### Applicability of PC dyes for Live-cell STED-FLIM nanoscopy

Since maintenance of mt-nucleoid is essential for proper mt-DNA segregation and replication^40,41^, probes directly staining mt-DNA as well as nuclear DNA is very effective tool for mitochondrial biomedical research. As far as we know, useful probes compatible with live cell super-resolution microscopy, however, have not been reported till now in spite of great needs for elucidating mt-nucleoids of which their size are less than a diffraction limit^42.43^. Therefore, we assessed the compatibility of PC dyes to STED-FLIM nanoscopy and finally found that **PC3** was a promising PC dye. HeLa cells stained with 10 nM **PC3** were excited with 561 nm with or without 660 nm STED laser by sequential imaging using between lines. Super-resolution images were obtained by two component separations of STED-FLIM image using n-exponential reconvolution model (**Fig. 4b**). Comparable results were also obtained not only in living NIH/3T3 cells but also in plant cells (**Fig. S9** and **S10**). Then, we also calculated a full width at half maximum (FWHM) of minor axis of mt-nucleoids to estimate the size of mt-nucleoids in HeLa cells (**Fig. 4c**). The FWHM value was 100±9 nm at 3 ns of delay time, which is good agreement with the previous study^44^. Taken together, **PC3** is a unique DNA probe which has a great compatibility with STED nanoscopy in various cell types.

**Figure 4.**
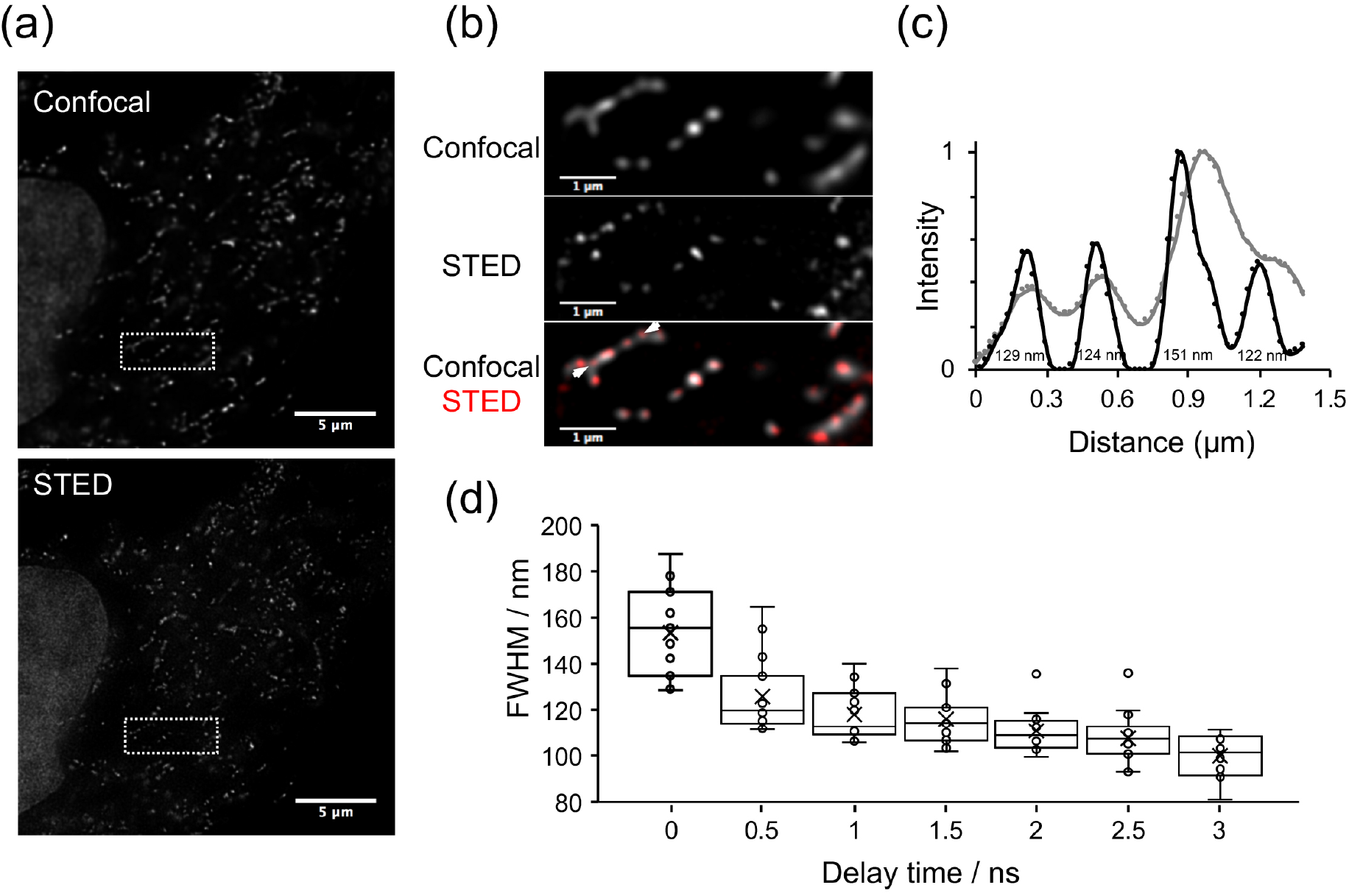
Comparison of confocal and STED-FlIM imaging in living HeLa cells stained with PC3. **(a)** Confocal and STED-FLIM images of mt-DNA stained with 10 nM **PC3** in living HeLa cells. **(b)** Enlarged images of the white dotted square region of (a). **(c)** An example of normalized fluorescence intensity profiles obtained from the region between arrows in (b). Line profiles in STED and confocal image are shown in black and gray, respectively. FWHM values estimated by fitting with a Gaussian function are also indicated in the black line profile. **(d)** Box plot of FWHM value as a function of delay time in the minor axis of mt-nucleoids (n=15 from independent cells).

## Discussion

In this study, we developed new series of DNA markers based on N-aryl pyrido cyanine (PC) to fulfill all three required properties described above. Firstly, we actualized that all N-aryl PC dyes were applicable to long wavelength excitation over 532 nm as planned. This achievement is due to our molecular design focusing on PC^35^, possessing an extended π-conjugation with a monomethine cyanine unit compared with hitherto unsymmetrical cyanine dyes such as SYBR-Green and Pico-Green. Secondly, fortunately, all eight PC dyes including trimethine dyes have high DNA selectivity and are capable of staining nucleus. Among them, **PC1** and **PC3** have an unexpectedly great property of DNA sequence selectivity at AT-pairs rich region and a high fluorogenic property upon binding DNA, succeeding in staining nuclear DNA at very low concentration. And thirdly, these dyes exhibited extremely high cell and tissue permeability to stain nuclear DNA in various cell types. Especially in **PC1** stained plant root, 2PEM revealed whole nuclei without apparent toxicity in root growth and cell division.

We also succeeded to bring out other key potentials of PC dyes except for basic ideal properties described above. We found that these PC dyes obviously stained mt-DNA at ultralow concentration and also labelled both mt-DNA and nuclear DNA by optimizing the staining dye concentration. Since the synthetic probes for mt-DNA have been limited to staining DAPI^45^, SYBR-Green^9,27^, and Pico-Green^28^, our red fluorescent PC dyes will provide researchers with new channel. More importantly, taking advantage of this character, we also successfully applied FLIM into mammalian cells for discrimination between mt-DNA and nuclear DNA and into plant cells for separation with ch-DNA as well as mt-DNA and nuclear DNA by itself. Furthermore, we demonstrated that at least one PC dye (**PC3**) were applicable to STED-FLIM nanoscopy. Our dye will be useful for the biology field in mt-nucleoid as well as nuclear chromatin dynamics. Recently, dynamic structure of mitochondrial cristae has been visualized by STED nanoscopy^46, 47^. In regards to the dynamic structural relationship between mt-nucleoid and mitochondrial cristae, however, dual STED imaging has not been reached yet since PicoGreen which stains mt-nucleoids were not be applied to STED nanoscopy while super-resolution of cristae was perfectly performed using SNAP-Cell Sir labeling system in COX8A-SNAP expressed cells^46^. We believe that N-aryl PC dyes should enable the challenging study. Last of all, together with these outstanding characters and talented applicability to various microscope techniques, our PC dyes have both the best of Hoechst and Pico-Green. Although we here did not validate the effectiveness, we also believe that PC dyes would be also useful tools for molecular biology such as real-time PCR and fluorescent cytometry.

## Methods

### Binging of Hair-pin DNA and RNA Oligonucleotides

For the oligo-nucleotides binding studies, synthetic DNA oligonucleotides, 5’-CGCGAATTCGCGTTTTCGCGAATTCGCG-3’ (28 bp) and 5’-CGCGCCGGCGCGTTTTCGCGCCGGCGCG-3’ (28 bp) were purchased by eurofins, whereas RNA 5’-CGCGAAUUCGCGUUUUCGCGAAUUCGCG-3’ (28 bp) was obtained from FASMAC. Each oligonucleotide was dissolved in 1 × TBS buffer (50 mM Tris HCl, 150 mM NaCl, pH 7.4) at 200 μM concentration and adjusted various concentrations by serial dilution with TBS. These oligonucleotide solutions were heated at 95°C for 1 min followed by cooling down at room temperature. On the other hand, each PC dye was dissolved in TBS with 2 mg/mL BSA at 200 nM concentration. The fluorescent intensity of equal amount mixture of oligonucleotide and PC dye solution was measured by EnSpire (PerkinElmer).

### Animal and plant cell cultures for fluorescence imaging

Cell culture lines (HeLa, U-2OS, C6, NIH3T3) were cultured in Dulbecco’s modified Eagle’s medium (DMEM, Wako) containing 10% fetal bovine serum at 37 °C in a 5% CO_2_/95% air incubator. These lines (2 × 10^4^ cells /mL) were transferred on each well of a glass-bottom 8-well slide and cultured 1day before imaging. DNA staining was performed in DMEM (-) containing 10 mM HEPES (pH 7.4) without washing. The *Arabidopsis thaliana* wild-type (Col-0) was also used. After keeping at 4°C for 3 days on Murashige and Skoog medium, seeds were cultured under continuous white light at 22-23°C for germination and cultured for 11-12 days.

### Fluorescent titration

Calf thymus double stranded DNA (dsDNA), purchased from Sigma-Aldrich Co, and Ribonucleic acid from torula yeast (RNA), purchased from Wako Pure Chemical Industries, were used in the fluorescence titration^48,49^. DNA/RNA selectivity of **PC1** was compared with commercialized nucleus markers (Pico-Green, Hoechst 33342; ThermoFisher Scientific). All chemicals are used without additional treatment or further purifications. UV/Vis absorption spectra were recorded on a Shimadzu UV-3510 spectrometer with a resolution of 0.5 nm and emission spectra were measured with an FP-6600 Hitachi spectrometer with a resolution of 0.2 nm. Circular dichroism spectra were measured with a JASCO FT/IR6100. 1.0 cm square quartz cell was used for all optical measurements. 1.0 g/L dsDNA solution (1.0 mL) or 2.0 g/L RNA solution (1.0 mL) were added to dye solutions (2.0 mL) with absorbance around 0.2 at each maximum wavelength at room temperature by using a micro pipet. After titration, the combined solution was gently shaken several times to stabilize the absorbance and fluorescence intensities of all samples. (see detailed results in the supporting information)

### Wide-field microscopy

For assessment of cytotoxicity and photo-toxicity of PC dyes, we also used commercialized red fluorescence nucleus markers (SYTO 80, SYTO 82, SYTO 84; ThermoFisher Scientific). The time-lapse observation was performed by an inverted microscope system (IX-71; Olympus) equipped with an UPlanSApo IR 20x/0.75 objective lens (Olympus), and a CMOS camera (ORCA Flash 4.0 V3 C13440; Hamamatsu photonics). The TRITC-A-Basic fluorescent filter set (FF01-542/20, FF570-Di01, FF01-620/52; Semrock Inc.) was used for all nucleus markers. The stage incubator system (Tokai Hit Co, Ltd.) was used to keep temperature at 37 °C and 5% CO_2_/95% air condition. The fluorescence and bright-field time lapse images were taken with or without excitation and cell proliferation rate was assessed by visual inspection from bright-field time-lapse images.

### Confocal microscopy

A confocal laser scanning microscopy system (TCS-SP8 FALCON gSTED; Leica) equipped with a pulsed white light laser (WLL; 80 MHz) and a HyD detector was used for fluorescence imaging of nuclear DNA in various animal cultured cells at 37 °C in a 5% CO_2_/95% air condition (Fig. 2a and 2c). For low and high magnification observation, HC PL APO CS2 20× 0.75 and HC PL APO CS2 100× 1.40 oil objective were used, respectively. Cells stained with **PC1** or SYTO 80 were excited with a 532 nm and their emission was collected at 540 – 670 nm. Cells stained with **PC3** were excited with a 552 nm or a 561 nm and their emission was collected at 560 - 670 nm or 570-670 nm. When stained at ultralow concentration (100 pM), cells were excited with 561 nm and their emission was detected at 570-769 nm. Cells stained with SYTO 82 were excited with a 543 nm and their emission was detected at 550-670 nm. Cells stained with SYTO 84 were excited with a 561 nm and their emission was detected at 570-670 nm. Gated detection between 0.1-12 ns was performed for all fluorescence imaging. A confocal microscope system (LSM 780; Zeiss) equipped with a 20×/0.8 Plan-Apochromat lens and 32-channel gallium arsenide phosphide (GaASP) detector array was used for Arabidopsis leaf and root imaging. Cells stained with **PC1** were excited with a 514 nm and their fluorescence were collected at 517-614 nm in leaf cells and 517-693 nm in root cells, respectively. Cells stained with **PC3** were excited with a 560 nm and their fluorescence were collected at 561-605 nm in leaf cells and 570-693 nm in root cells, respectively. In leaf cell imaging, chlorophyll autofluorescence was also detected at 675-693 nm. Collected images were further processed using open-source software Image J (http://imagej.nih.gov/ij/).

### Two-photon excitation microscopy

Two-photon imaging was performed using a laser scanning microscope (LSM-780; Zeiss) equipped with a widely tunable Ti: Sapphire femtosecond pulse laser (Chameleon; Coherent) and LD C-Apochromat 40×/1.1 water immersion lens. The same Arabidopsis root stained with **PC1** were excited with 1000 nm as well as 488 nm and their fluorescence were detected at 500-690 nm and 490-596 nm, respectively.

### FLIM and STED-FLIM microscopy

A confocal laser scanning microscopy system (TCS-SP8 FALCON gSTED; Leica) equipped with a pulsed white light laser (WLL; 80 MHz), 660 STED laser, HC PL APO CS2 100൹/1.40 oil objective lens, and a HyD detector was used for fluorescence imaging of nuclear and mitochondrial DNA in various animal cultured cells at 37°C in a 5% CO2/95% air condition (Fig. 4a-i). For observation of Arabidopsis stomata, HC PL APO CS2 93×/1.30 GLYC objective lens was used and z-sectioning image was obtained from 26 frames at 0.26 μm steps. For STED-FLIM imaging, cells were excited with 561 nm and their emission was corrected at 570 – 650 nm with or without 660 nm STED laser. STED image was obtained by separation of a FLIM image to two exponential components thorough n-exponential reconvolution model or τ-STED function. Confocal and STED imaging were acquired alternately between lines. Fluorescent life time based separation images were displayed with different pseudo colors by phaser plot analysis^50^. Collected images were deconvoluted by default setting of Huygens; signal-to-noise ratio and quality threshold were set to 7 and 0.05 for STED images, 20 and 0.05 for conventional CLSM images, respectively. Images were further processed using ImageJ. Full-width-half-maximum (FWHM) was estimated by fitting with a Gaussian function described before^47^.

## Acknowledgements

We are deeply grateful to Prof. K. Itami, Prof. and Prof. T. Higashiyama at Nagoya University for enormous help of this work such as invaluable comments, discussion and financial support. We thank Dr. H. Ito, Dr. Y. Segawa, Dr. T. Fujikawa and Dr. K. Kato at Nagoya University for many helpful comments. We also thank M. Tsuzuki (Nagoya University) for supporting cell culture and cell toxicity assessment. This work was supported by Grant-in-Aid for JSPS Research Fellow (14J03652 to K.U.), Japan Society for Scientific Research (19H05364, 20H05412 to Y.S., 16H06464, 16K21727, JP16H06280 to T.H), Toyoaki scholarship Foundation and Ohsumi Frontier Science Foundation to Y.S. and the ERATO program from JST (JPMJER1302 to K.I.).

## Author contributions

K.U. and Y.S conceived and designed this research. K.U performed synthesis of all PC dyes and most of the spectroscopic measurements. Y.S. and N.S. performed fluorescence titration experiments using hairpin oligonucleotides and all imaging experiments. U.K. and Y.S wrote the manuscript. All authors read and approved the manuscript.

**Additional information Supplementary information is available for this paper at https://doi.org/\*\*\*\*\***

