## Supplemental Information for "N-Aryl Pyrido Cyanine derivatives: nuclear and organelle DNA markers for two-photon and super-resolution imaging"

**Suuplementary Information for**

This PDF file includes:

**1. Supplementary methods**
**2. Chemical synthesis**
**3. Figure S1 to S9**
**4. Caption for movies S1 to S6**
**5. NMR Charts.**
**6. References for SI reference citations**

Other supplementary materials for this manuscript include the following:
Movies S1 to S6

### 1. Supplementary methods

**Fluorescence lifetime.** The light source used was a wavelength-tunable optical parametric amplifier based on a regeneratively amplified mode-locked Ti: sapphire laser with a pulse duration of 200 fs and a repetition rate of 200 kHz. The excitation wavelengths were set around the maximum absorption wavelengths for each sample. Emitted photons were detected using a single monochromator with an avalanche photodiodes (SPD-050-CTE-N1; MPD). The detection wavelengths were tuned to the emission peak wavelengths for each sample. Each photon arrival times were recorded using time-correlated single-photon counting boards (SPC-130EM-N1; Becker & Hickl GmbH).

**Fluorescence quantum yield.** The fluorescence quantum yields of dye-dsDNA complexes were determined with a Hamamatsu C9920-02 calibrated by the integrating sphere system. Measurements of absolute fluorescence quantum yields ($\Phi_F^{dsDNA}$) of PC dyes were performed with scan excitation mode at around maximum absorption wavelength ($\lambda_{abs}^{dsDNA}$), and values of $\Phi_F^{dsDNA}$ of monomethine dyes were calculated by using averaged values. Relative method was applied to determine the intrinsic fluorescence quantum yields ($\Phi_F^{free}$). The values of $\Phi_F^{free}$ for monomethine and trimethine PC are deterimined in relative to the of Rhodamine 6G ethanol in solution ($\Phi_F = 0.94$)[1] and Cresyl violet in methanol solution ($\Phi_F = 0.54$)[2] according to following equations.

$$\Phi_F^{free} = \Phi_F^{referece} \times (F^{free\text{-}dye}/F^{reference}) \times (A^{reference}/A^{free\text{-}Dye}) \times (n^{free\text{-}dye}/n^{reference})^2$$

where $\Phi_F^{referece}$ represents fluorescence quantum yield of reference samples, $F^{free\text{-}dye}$ and $F^{reference}$ stand for the area of corrected fluorescence spectra of dyes and reference sample using same excitation wavelength, $A^{reference}$ and $A^{free\text{-}Dye}$ are the recorded absorbance of reference sample and dyes at the wavelength for the photo-excitation, and $n^{free\text{-}dye}$ and $n^{reference}$ represent the reflective index of the solvents for the measurements. Reflective index value of TE buffer solution was calculated using that of water ($n = 1.333$). The values of reflective index for ethanol ($n = 1.359$)[3] and methanol ($n = 1.327$)[3] were used for the detailed calculation.

**Co-staining of N-aryl PC dyes with Hoechst 33342 in living HeLa cells.** HeLa cells were co-stained with 100 nM PC dyes with various concentration of Hoechst 33342 (0 nM, 100 nM, 1 µM, 3 µM). The confocal images were obtained with a TCS SP8 (Leica) equipped with HC PL APO CS2 100×/1.40 oil objective. The image channels used were Ex 405 nm and Em 420-480 nm for Hoechst 33342, Ex 532 nm / Em 540-670 nm for **PC1**, and Ex 561 nm / Em 570-670 nm for **PC3**, respectively.

**Co-staining of PC dyes with a mitochondrial staining dye.** HeLa cells were co-stained with 1 nM PC dyes with 20 nM MitoTracker Deep Red. The confocal images were obtained with a TCS SP8 (Leica) equipped with HC PL APO CS2 100×/1.40 oil objective. The image channels used were Ex 514 nm and Em 520-610 nm for **PC1**, Ex 561nm / Em 570-610 nm for **PC3**, and Ex 633 nm / Em 640-680 nm for MitoTracker Deep Red, respectively.

**Time-lapse analysis of Arabidopsis root and root hairs**Arabidopsis root stained with 1 µM **PC1** was observed every 5 min excited with 488 nm and the emission spectrum was collected through band-pass filter BP525/50. The fluorescence images are maximum z-projections of 20 planes (4.3-µm intervals).

### 2. Chemical synthesis

Unless otherwise noted, all materials including dry solvents were obtained from commercial suppliers and used without further purification. The starting materials such as 1-(2,4-dinitrophenyl)-4-methylpyridin-1-ium chloride[4] and **5a**[5] were prepared according to the reported procedures. All work-up and purification procedures were carried out with reagent-grade solvents. High-resolution mass spectra analyses were carried out using FT-ESI mass analyzer. Melting points of all compounds were measured on a MPA100 Optimelt automated melting point system. Column chromatography was performed with silica gel 60N (Kanto Chemical Co., spherical, neutral, 40–50 mesh) for the purification of **PC1**, **PC2**, **PC3**, **PC4**, **PC5**, **PC6, PC7**, and **PC8**. Column chromatography was performed with amino silica gel (Fuji Sylysia Chemical LTD, Cat. No. Hu 41003) for the purification of **8a**, **8b**, **8c**, **8d**, and **PC8**. Purifications of the other compounds were performed by silica gel 60N (Kanto Chemical Co., spherical, neutral, 40–100 mesh).

Nuclear magnetic resonance (NMR) spectra were recorded on JEOL JNM-ECA-400 ($^1$H 400 MHz, $^{13}$C 100 MHz) spectrometers in dimethyl dimethyl sulfoxide-d6 (d-DMSO) ($(CD_3)_2SO$). Chemical shifts for $^1$H NMR are expressed in parts per million (ppm) relative to $(CD_3)_2SO$ (δ 2.49 ppm). Chemical shifts for $^{13}$C NMR are expressed in ppm relative to $(CD_3)_2SO$ (δ 39.5 ppm). Data are reported as follows: chemical shift, multiplicity (s = singlet, d = doublet, dd = doublet of doublets, ddd = doublet of doublet of doublets, dt = doublet of triplets, td = triplet of doublets, t = triplet, q = quartet, quin = quintet, m = multiplet, br = broad), coupling constant (Hz), and integration.

Synthesis of **PC1**, **PC2**, **PC3**, and **PC4**.

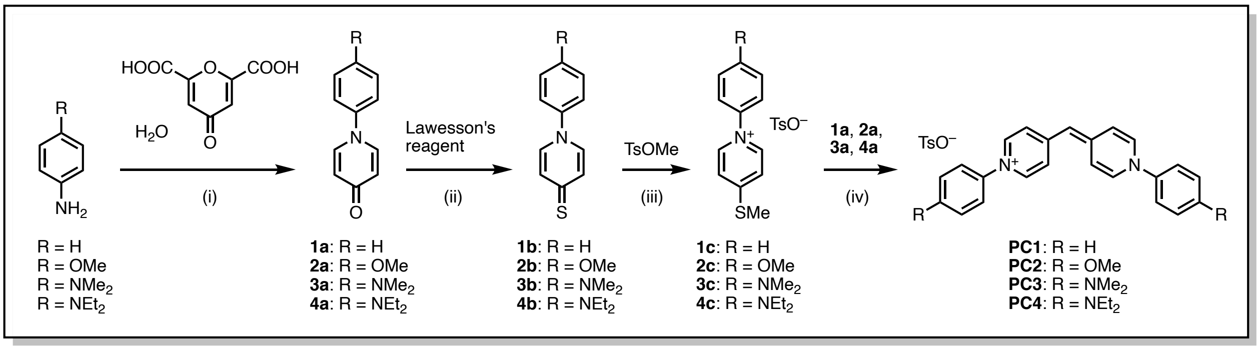


Scheme S1. (i) Chelidonic acid monohydrate, aniline derivatives (1.1 equiv.), DMSO, 170 °C, 2 h; (ii) Lawesson's reagent, toluene/1,4-dioxane (= 10/1), reflux, 3–4 h; (iii) TsOMe, 1,4-dioxane, reflux, 2 h; (iv) **1a, 2a, 3a**, and **4a** in dry THF, MeLi, –0 °C. $CH_2Cl_2$/triethylamine, 70 °C, reflux, 1 d.

#### 1-Phenylpyridin-4(1*H*)-one (1a)

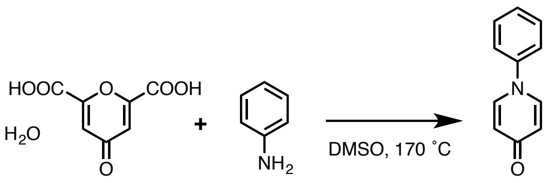


**1a**

Chelidonic acid monohydrate (3.00 g, 14.8 mmol) and aniline (1.45 g, 15.6 mmol, 1.05 equiv.) were dissolved in DMSO (45 mL). The mixture was stirred at 170 °C under open air for 2 h. After cooling back to room temperature, DMSO was removed *in vacuo*. The resulting dark residue was subjected to column chromatography on silica gel (eluent: $CH_2Cl_2$/MeOH = 95/5 to 80/20; *v*/*v*). Recrystallization from methanol and ether gave **1a** as a pale yellow solid (1.45 g, 57%).

$^1$H NMR (400 MHz, $(CD_3)_2SO$): δ 6.23 (d, *J* = 7.6 Hz, 2H), 7.40–7.48 (m, 1H), 7.54 (s, 2H), 7.55 (s, 2H), 7.99 (d, *J* = 8.4 Hz, 2H). $^{13}$C NMR (100 MHz, $(CD_3)_2SO$): δ 117.96, 122.60, 127.87, 129.98,

139.91, 142.76, 177.42. HRMS (ESI positive mode) *m/z* calculated for $C_{11}H_9NNaO$ $[M+Na]^+$: 194.0576, found: 194.0574. Mp = 78.2–79.0 °C.

#### 1-Phenylpyridine-4(1*H*)-thione (1b)

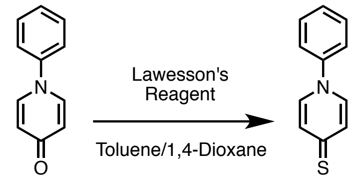


**1a** **1b**

A solution of **1a** (1.00 g, 5.84 mmol) and Lawesson's reagent (2.48 g, 6.13 mmol, 1.05 equiv.) in toluene (50 mL) and 1,4-dioxane (5.0 mL) was stirred at refluxing temperature for 4 h. After cooling back to room temperature, hexane (20 mL) was added to the mixture and the formed solid was filtered. The crude product was purified by column chromatography on silica gel (eluent: $CH_2Cl_2$/MeOH = 99/1 to 70/30; *v/v*). Recrystallization from methanol and ether gave **1b** as a yellow solid (569 mg, 52%).

$^1$H NMR (400 MHz, $(CD_3)_2SO$): δ 7.26 (d, *J* = 7.6 Hz, 2H), 7.51 (t, *J* = 7.0 Hz, 1H), 7.54–7.56 (m, 4H), 7.92 (d, *J* = 7.2 Hz, 2H). $^{13}$C NMR (100 MHz, $(CD_3)_2SO$): δ122.83, 128.83, 130.11, 130.32, 134.84, 142.39, 191.16. HRMS (ESI positive mode) *m/z* calculated for $C_{11}H_9NNaS$ $[M+Na]^+$: 210.0348, found: 210.0345. Mp = 149.5–150.2 °C.

#### 4-(Methylthio)-1-phenylpyridin-1-ium 4-methylbenzenesulfonate (1c)

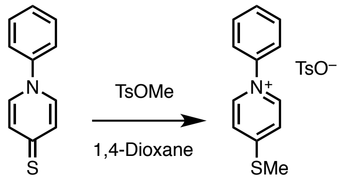


**1b** **1c**

A solution of **1b** (420 mg, 2.24 mmol) and methyl *p*-toluenesulfonate (438 mg, 2.35 mmol, 1.05 equiv.) in 1,4-dioxane (20 mL) was stirred at refluxing temperature for 2 h. After cooling back to room temperature, ether (20 mL) was added to the mixture. The upper layer was separated, and the bottom layer containing **1c** was washed with ether twice. The resulting oil was dried *in vacuo* for 2 h. The obtained crude **1c** (820 mg, 98%) was used to the next reaction without further purification.

#### 1-Phenyl-4-((1-phenylpyridin-4(1*H*)-ylidene)methyl)pyridin-1-ium4-methylbenzenesulfonate (PC1)

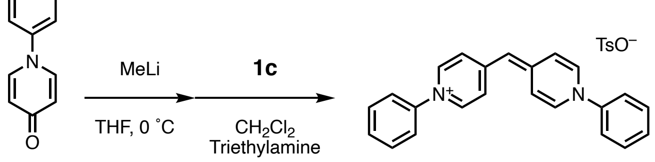


**1a** **PC1**

A solution of **1a** (300 mg, 1.75 mmol) in THF (20 mL) was cooled to 0 °C by an ice bath under nitrogen atmosphere. 3 M MeLi solution in diethoxymethane (1.2 mL, 3.6 mmol, 2.0 equiv.) was slowly added dropwise using a syringe. The ice bath was removed and the mixture was warmed to room temperature gradually. After additional stirring for 20 min, a solution of **1c** (720 mg, 1.93 mmol, 1.1 equiv.) and triethylamine (4.0 mL) in $CH_2Cl_2$ (20 mL) was added using a syringe at 0 °C. Then, the mixture was stirred at 70 °C for 1 day under nitrogen atmosphere. After cooling back to room temperature, the solvent was removed *in vacuo*. The resulting dark red solid was subjected to column chromatography on silica gel (eluent: $CH_2Cl_2$/MeOH = 99/5 to 80/20; *v*/*v*). The obtained red solid was suspended to minimum amount of acetone and sonicated for 5 min, and then ether was further added. The formed solid was filtered to give **PC1** as a red solid (355 mg, 41%).

$^1$H NMR (400 MHz, $(CD_3)_2SO$): δ 2.27 (s, 3H), 5.82 (s, 1H), 7.09 (d, *J* = 8.4 Hz, 2H), 7.18–7.40 (br, 4H), 7.46 (d, *J* = 8.4 Hz, 2H), 7.54 (t, *J* = 6.8 Hz, 2H), 7.58–7.72 (m, 8H), 8.19 (d, *J* = 7.6 Hz, 4H). $^{13}$C NMR (100 MHz, $(CD_3)_2SO$): δ 20.74, 100.91, 117.04 (br), 122.77, 125.46, 127.98, 128.82, 130.11, 137.46, 138.39, 142.16, 145.87, 149.61. HRMS (ESI positive mode) calculated for $C_{23}H_{19}N_2$ $[M]^+$: 323.1543, found: 323.1537. Mp = 214.1–215.0 °C.

**1-(4-Methoxyphenyl)pyridin-4(1*H*)-one (2a)**

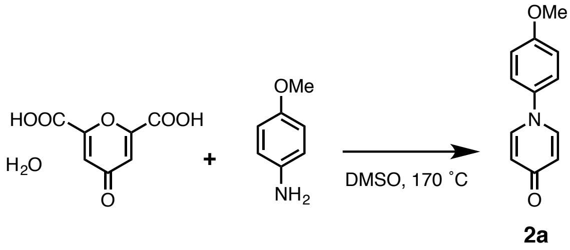


Chelidonic acid monohydrate (1.00 g, 4.95 mmol) and *p*-anisidine (640 mg, 5.2 mmol, 1.05 equiv.) were dissolved in DMSO (20 mL). The mixture was stirred at 170 °C under open air for 2 h. After cooling back to room temperature, DMSO was removed *in vacuo*. The resulting solid was suspended in ethyl acetate (30 mL) and vigorously stirred for 2 h. After addition of ether (30 mL), the solid was filtered and dried to give **2a** as a pale yellow solid (863 mg, 87%).

$^1$H NMR (600 MHz, $(CD_3)_2SO$): δ 3.80 (s, 3H), 6.19 (d, *J* = 7.8 Hz, 2H), 7.07 (d, *J* = 8.4 Hz, 2H), 7.46 (d, *J* = 9.0 Hz, 2H), 7.88 (d, *J* = 7.2 Hz, 2H). $^{13}$C NMR (150 MHz, $(CD_3)_2SO$): δ 55.54, 114.86, 117.69, 124.08, 136.09, 140.11, 158.62, 177.10. HRMS (ESI positive mode) *m*/*z* calculated for $C_{12}H_{11}NNaO_2$ $[M+Na]^+$: 224.0682, found: 224.0679. Mp = 182.0–183.4 °C.

**1-(4-Methoxyphenyl)pyridine-4(1*H*)-thione (2b)**

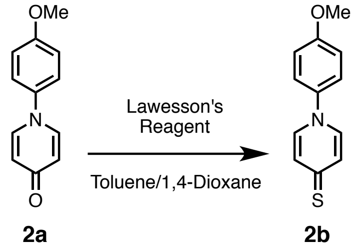


A solution of **2a** (500 mg, 2.48 mmol) and Lawesson's reagent (1.06 g, 2.61 mmol, 1.05 equiv.) in toluene (20 mL) and 1,4-dioxane (2.0 mL) was stirred at refluxing temperature for 4 h. After cooling back to room temperature, ether (20 mL) was added to the mixture. The formed yellow solid was filtered. The obtained solid was subjected to column chromatography on silica gel (eluent: $CH_2Cl_2$/MeOH = 95/5 to 50/50; *v*/*v*) to give **2b** as a yellow solid (252 mg, 47%).

$^1$H NMR (600 MHz, $(CD_3)_2SO$): δ 3.81 (s, 3H), 7.11 (d, *J* = 8.4 Hz, 2H), 7.24 (d, *J* = 6.6 Hz, 2H), 7.55 (d, *J* = 9.0 Hz, 2H), 7.83 (d, *J* = 6.6 Hz, 2H). $^{13}$C NMR (150 MHz, $(CD_3)_2SO$): δ 55.64, 114.99, 124.23, 130.14, 135.05, 135.63, 159.29, 190.42. HRMS (ESI positive mode) *m*/*z* calculated for $C_{12}H_{11}NNaOS$ $[M+Na]^+$: 240.0454, found: 240.0450. Mp = 190.9–192.1 °C.

**1-(4-Methoxyphenyl)-4-(methylthio)pyridin-1-ium 4-methylbenzenesulfonate (2c)**

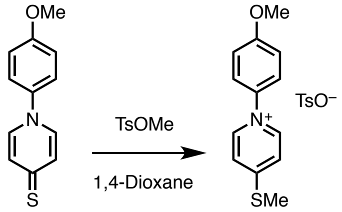


**2b** **2c**

A solution of **2b** (230 mg, 1.06 mmol) and methyl *p*-toluenesulfonate (216 mg, 1.16 mmol, 1.10 equiv.) in 1,4-dioxane (10 mL) was stirred at refluxing temperature for 2 h. After cooling back to room temperature, ether (30 mL) were added to form the precipitation which was then filtered, collected, and dried to give **2c** as a white solid (420 mg, 98%).

$^1$H NMR (600 MHz, $(CD_3)_2SO$): δ 2.73 (s, 3H), 3.22 (s, 3H), 4.32 (s, 3H), 7.55 (d, *J* = 7.8 Hz, 2H), 7.68 (d, *J* = 8.4 Hz, 2H), 7.92 (d, *J* = 7.8 Hz, 2H), 8.20 (d, *J* = 8.4 Hz, 2H), 8.47 (d, *J* = 6.6 Hz, 2H), 9.38 (d, *J* = 6.0 Hz, 2H). $^{13}$C NMR (150 MHz, $(CD_3)_2SO$): δ 14.17, 20.73, 55.85, 115.07, 122.32, 125.45, 125.67, 127.97, 135.16, 137.46, 141.92, 145.85, 160.68, 164.25. HRMS (ESI positive mode) *m*/*z* calculated for: $C_{13}H_{14}NOS$ $[M]^+$: 232.0791, found: 232.0787. Mp = 155.0 –156.0 °C.

**1-(4-Methoxyphenyl)-4-((1-(4-methoxyphenyl)pyridin-4(1*H*)-ylidene)methyl)pyridin-1-ium 4-methylbenzenesulfonate (PC2)**

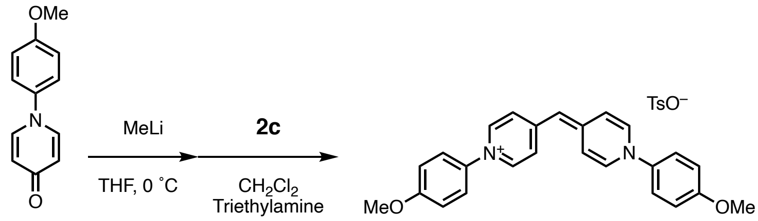


**2a** **PC2**

A solution of **2a** (200 mg, 0.99 mmol) in THF (10 mL) was cooled to 0 °C by ice bath under nitrogen atmosphere. 3 M MeLi solution in diethoxymethane (0.66 ml, 2.0 mmol, 2.0 equiv.) was slowly added dropwise for 5 min. The ice bath was removed and the reaction mixture was warmed to room temperature gradually. After additional stirring for 20 min, a solution of **2c** (400 mg, 0.99 mmol) and triethylamine (2 mL) in $CH_2Cl_2$ (10 mL) was added using a syringe at 0 °C. The reaction solution was refluxed at 70 °C for 24 h under nitrogen atmosphere. After cooling back to room temperature, the solvent was removed *in vacuo*. Then column chromatography on silica gel (eluent: $CH_2Cl_2$/MeOH = 95/5 to 80/20; *v*/*v*) was performed for purification. The obtained red solid was suspended to minimum amount of acetone and sonicated for 5 min and then ether was further added. The solid was filtered to give **PC2** as a red powder (153 mg, 28%).

$^1$H NMR (600 MHz, $(CD_3)_2SO$): δ 2.27 (s, 3H), 3.82 (s, 6H), 5.75 (s, 1H), 7.09 (d, *J* = 7.2 Hz, 2H), 7.14 (d, *J* = 7.2 Hz, 4H), 7.18–7.34 (br, 4H), 7.47 (d, *J* = 6.6 Hz, 2H), 7.58 (d, *J* = 7.8 Hz, 4H), 8.08 (d, *J* = 6.6 Hz, 4H). $^{13}$C NMR (150 MHz, $(CD_3)_2SO$): δ 20.75, 55.69, 100.36, 115.08, 124.21, 125.48, 128.00, 135.47, 137.50, 138.49, 145.85, 149.24, 159.35. HRMS (ESI positive mode) *m*/*z* calculated for $C_{25}H_{23}N_2O_2$ $[M]^+$: 383.1754, found: 383.1745. Mp = 196.0–196.8 °C.

**1-(4-(Dimethylamino)phenyl)pyridin-4(1*H*)-one (3a)**

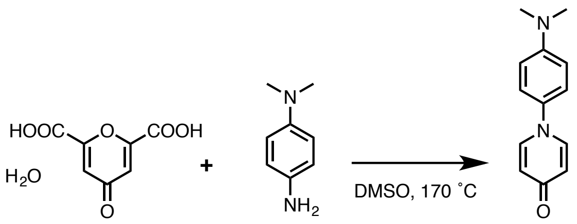


**3a**

Chelidonic acid monohydrate (1.00 g, 4.95 mmol) and *N*,*N*-dimethyl-1,4-phenylenediamine (741 mg, 5.44 mmol, 1.10 equiv.) were dissolved in DMSO (15 mL). The mixture was stirred at 170 °C under open air for 2 h. After cooling back to room temperature, DMSO was removed *in vacuo*. The dark residue was subjected to column chromatography on silica gel (eluent: $CH_2Cl_2$/MeOH = 95/5 to 80/20; *v*/*v*). Recrystallization from methanol and ether gave **3a** as pale yellow solid (792 mg, 75%).

$^1$H NMR (400 MHz, $(CD_3)_2SO$): δ 2.93 (s, 6 H), 6.19 (d, *J* = 8.0 Hz, 2H), 6.80 (d, *J* = 9.2 Hz, 2H), 7.31 (d, *J* = 9.2 Hz, 2H), 7.85 (d, *J* = 7.6 Hz, 2H). $^{13}$C NMR (150 MHz, $(CD_3)_2SO$): δ 40.10, 112.57, 117.53, 123.43, 132.25, 140.35, 149.82, 176.86. HRMS (ESI positive mode) *m*/*z* calculated for $C_{13}H_{14}N_2NaO$ $[M+Na]^+$: 237.0998, found: 237.0994. Mp = 181.5–182.2 °C.

**1-(4-(Dimethylamino)phenyl)pyridine-4(1*H*)-thione (3b)**

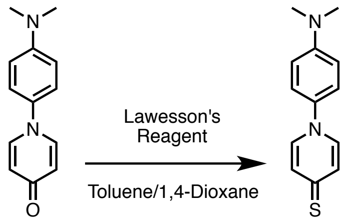


**3a** **3b**

A solution of **3a** (600 mg, 2.80 mmol) and Lawesson's reagent (1.19 g, 2.94 mmol, 1.05 equiv.) in toluene (25 mL) and 1,4-dioxane (2.5 mL) was stirred at refluxing temperature for 3 h. After cooling back to room temperature, ether (30 mL) was added to form the precipitation that was then filtered. The solid was subjected to column chromatography on silica gel (eluent: $CH_2Cl_2$/MeOH = 99/1 to 60/40; *v*/*v*). Recrystallization from methanol and ether gave **3b** as a yellow solid (347 mg, 54%).

$^1$H NMR (600 MHz, $(CD_3)_2SO$): δ 2.95 (s, 6H), 6.82 (d, *J* = 9.0 Hz, 2H), 7.23 (d, *J* = 6.6 Hz, 2H), 7.39 (d, *J* = 9.0 Hz, 2H), 7.80 (d, *J* = 7.2 Hz, 2H). $^{13}$C NMR (150 MHz, $(CD_3)_2SO$): δ 40.02, 112.47, 123.35, 130.19, 131.63, 135.03, 150.26, 189.49. HRMS (ESI positive mode) *m*/*z* calculated for $C_{13}H_{14}N_2NaS$ $[M+Na]^+$: 253.0770, found: 253.0768. Mp = 268.0–269.6 °C.

**1-(4-(Dimethylamino)phenyl)-4-(methylthio)pyridin-1-ium 4-methylbenzenesulfonate (3c)**

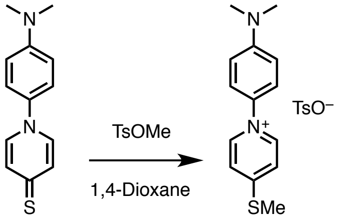


**3b** **3c**

A solution of **3b** (300 mg, 1.30 mmol) and methyl *p*-toluenesulfonate (245 mg, 1.32 mmol, 1.01 equiv.) in 1,4-dioxane (20 mL) was stirred at refluxing temperature for 2 h. After cooling back to room temperature, ether (30 mL) was added to form the precipitation. The resulting solid were collected and dried to give **3c** as a yellow solid (540 mg, 99%).

$^{1}$H NMR (600 MHz, $(CD_3)_2SO$): δ 2.27 (s, 3H), 2.75 (s, 3H), 3.00 (s, 6H), 6.89 (d, $J$ = 9.0 Hz, 2H), 7.09 (d, $J$ = 7.8 Hz, 2H), 7.46 (d, $J$ = 7.2 Hz, 2H), 7.58 (d, $J$ = 9.0 Hz, 2H), 7.96 (d, $J$ = 6.6 Hz, 2H), 8.87 (d, $J$ = 6.0 Hz, 2H). $^{13}$C NMR (150 MHz, $(CD_3)_2SO$): δ 14.12, 20.74, 112.17, 122.40, 124.53, 125.46, 127.99, 130.82, 137.48, 137.48, 141.39, 145.85, 151.31, 162.92. HRMS (ESI positive mode) *m/z* calculated for $C_{14}H_{17}N_2S$ $[M]^+$: 245.1107, found: 245.1107. Mp = 112.3–113.7 °C.

**1-(4-(Dimethylamino)phenyl)-4-((1-(4-(dimethylamino)phenyl)pyridin-4(1*H*)-ylidene)methyl)pyridin-1-ium 4-methylbenzenesulfonate (PC3)**

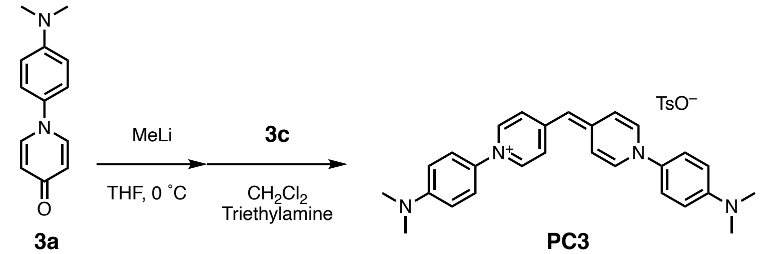


**3a** **PC3**

A solution of **3a** (250 mg, 1.17 mmol) in THF (20 mL) was cooled to 0 °C by ice bath under nitrogen atmosphere. 3 M MeLi solution in diethoxymethane (0.78 mL, 2.3 mmol, 2.0 equiv.) was slowly added dropwise using a syringe. The ice bath was removed and worm to room temperature gradually. After additional stirring for 20 min, a solution of crude **3c** (486 mg, 1.17 mmol) and triethylamine (4 mL) in $CH_2Cl_2$ (20 mL) was added using a syringe at 0 °C. The mixture was stirred at 70 °C for 24 h under nitrogen atmosphere. After cooling back to room temperature, the solvent was removed *in vacuo*. The resulting black solid was subjected to column chromatography on silica gel (eluent: $CH_2Cl_2$/MeOH = 99/1 to 85/15; *v*/*v*) for purification. The obtained purple solid was suspended to minimum amount of acetone and sonicated for 5 min and then ether was further added. The obtained solid was filtered to give **PC3** as a purple powder (227 mg, 34%).

$^{1}$H NMR (400 MHz, $(CD_3)_2SO$): δ 2.27 (s, 3H), 2.96 (s, 12H), 5.70 (s, 1H), 6.84 (d, $J$ = 9.2 Hz, 4H), 7.10 (d, $J$ = 7.6 Hz, 2H), 7.04–7.36 (br, 4H), 7.43 (d, $J$ = 9.2 Hz, 4H), 7.46 (d, $J$ = 8.4 Hz, 2H), 8.04 (d, $J$ = 6.8 Hz, 4H). $^{13}$C NMR (150 MHz, $(CD_3)_2SO$): δ 11.80, 31.04, 92.12, 104.40, 114.84, 117.48, 120.28, 123.80, 129.64, 132.10, 134.15, 141.60, 142.73. HRMS (ESI positive mode) *m/z* calculated for $C_{27}H_{29}N_4$ $[M]^+$: 409.2387, found: 409.2381. Mp = 194.8–195.7 °C.

**1-(4-(Diethylamino)phenyl)pyridin-4(1*H*)-one (4a)**

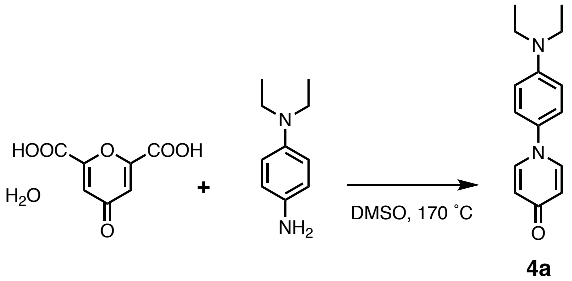


**4a**

Chelidonic acid monohydrate (1.00 g, 4.95 mmol) and *N*,*N*-diethyl-1,4-phenylenediamine (894 mg, 5.44 mmol, 1.10 equiv.) were dissolved in DMSO (15 mL). The mixture was stirred at 170 °C under open air for 2 h. After cooling back to room temperature, DMSO was removed *in vacuo*. The dark residues were subjected to column chromatography on silica gel (eluent: $CH_2Cl_2$/MeOH = 95/5 to 5/1; *v*/*v*) for purification. Recrystallization from methanol and ether gave **4a** as a white solid (917 mg, 77%).

$^{1}$H NMR (600 MHz, $(CD_3)_2SO$): δ 1.08 (t, $J$ = 7.2 Hz, 6H), 3.36 (q, $J$ = 6.6 Hz, 4H), 6.16 (d, $J$ = 7.8 Hz, 2H), 6.72 (d, $J$ = 9.0 Hz, 2H), 7.26 (d, $J$ = 8.4 Hz, 2H), 7.81 (d, $J$ = 7.8 Hz, 2H). $^{13}$C NMR (150 MHz, $(CD_3)_2SO$): δ 12.29, 43.77, 111.64, 117.58, 123.81, 131.32, 140.24, 146.84, 176.96. HRMS (ESI positive mode) *m/z* calculated. for $C_{15}H_{18}N_2NaO$ $[M+Na]^+$: 265.1311, found: 265.1310. Mp = 114.7–

116.3 °C.

**1-(4-(Diethylamino)phenyl)pyridine-4(1*H*)-thione (4b)**

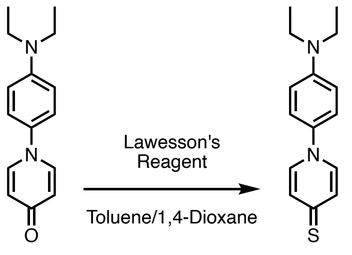


**4a** **4b**

A solution of **4a** (600 mg, 2.48 mmol) and Lawesson's reagent (1.05 g, 2.60 mmol, 1.05 equiv.) in toluene (25 mL) and 1,4-dioxane (2.5 mL) was stirred at refluxing temperature for 3 h. After cooling back to room temperature, ether (20 mL) was added and the formed solid was filtered. The collected solid was subjected to column chromatography on silica gel (eluent: $CH_2Cl_2$/MeOH = 99/1 to 80/20; *v*/*v*) to give **4b** as a yellow solid (427 mg, 67%).

$^1$H NMR (400 MHz, $(CD_3)_2SO$): δ 1.09 (t, *J* = 7.0 Hz, 6H), 3.37 (q, *J* = 7.2 Hz, 4H), 6.75 (d, *J* = 9.2 Hz, 2H), 7.22 (d, *J* = 7.2 Hz, 2H), 7.35 (d, *J* = 9.2 Hz, 2H), 7.79 (d, *J* = 6.8 Hz, 2H). $^{13}$C NMR (100 MHz, $(CD_3)_2SO$): δ 12.72, 43.81, 111.58, 123.72, 130.17, 130.69, 135.10, 147.37, 189.19. HRMS (ESI positive mode) *m*/*z* calculated for $C_{15}H_{18}N_2NaS$ $[M+Na]^+$: 281.1083, found: 281.1080. Mp = 166.2–167.9 °C.

**1-(4-(Diethylamino)phenyl)-4-(methylthio)pyridin-1-ium 4-methylbenzenesulfonate (4c)**

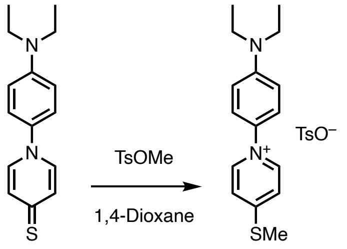


**4b** **4c**

A solution of **4b** (400 mg, 1.55 mmol) and methyl *p*-toluenesulfonate (292 mg, 1.57 mmol, 1.01 equiv.) in 1,4-dioxane (20 mL) was stirred at refluxing temperature for 2 h. After cooling back to room temperature, ether (30 mL) was added to form the precipitation. The resulting solid was filtered, collected and dried to give **4c** as a yellow solid (681 mg, 99%).

$^1$H NMR (600 MHz, $(CD_3)_2SO$): δ 1.11 (t, *J* = 7.2 Hz, 6H), 2.27 (s, 3H), 2.75 (s, 3H), 3.42 (q, *J* = 7.0 Hz, 4H), 6.84 (d, *J* = 9.6 Hz, 2H), 7.09 (d, *J* = 7.8 Hz, 2H), 7.46 (d, *J* = 7.8 Hz, 2H), 7.54 (d, *J* = 9.0 Hz, 2H), 7.96 (d, *J* = 7.8 Hz, 2H), 8.86 (d, *J* = 7.2 Hz, 2H). $^{13}$C NMR (150 MHz, $(CD_3)_2SO$): δ 12.21, 14.10, 20.73, 43.90, 111.42, 122.39, 124.82, 125.46, 127.98, 129.95, 137.48, 141.29, 145.84, 148.53, 162.68. HRMS (ESI positive mode) *m*/*z* calculated for $C_{16}H_{21}N_2S$ $[M]^+$: 273.1420, found: 273.1419. Mp = 140.5–141.6 °C.

**1-(4-(Diethylamino)phenyl)-4-((1-(4-(diethylamino)phenyl)pyridin-4(1*H*)-ylidene)methyl)pyridin-1-ium 4-methylbenzenesulfonate (PC4)**

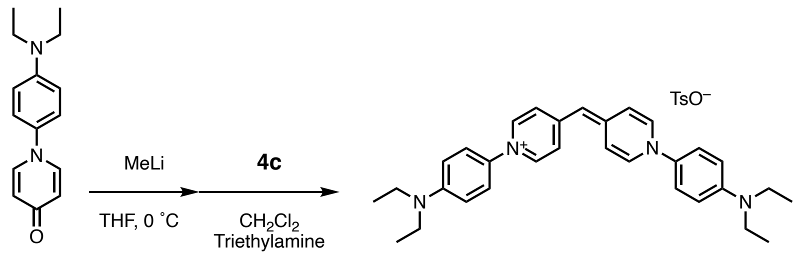


**4a** **PC4**

A solution of **4a** (200 mg, 0.83 mmol) in THF (15 mL) was cooled to 0 °C by ice bath under nitrogen atmosphere. 3 M MeLi solution in diethoxymethane (0.55 mL, 1.7 mmol, 2.0 equiv.) was slowly added dropwise using a syringe. The ice bath was removed and the reaction solution was warmed to room temperature gradually. After additional stirring for 20 min, a solution of **4c** (385 mg, 0.87 mmol, 1.1 equiv.) and triethylamine (2 mL) in $CH_2Cl_2$ (15 mL) was added using a syringe at 0 °C. The mixture was stirred at 70 °C for 24 h under nitrogen atmosphere. After cooling back to room temperature, the solvent was removed *in vacuo*. The resulting dark red solid was subjected to column chromatography on silica gel (eluent: $CH_2Cl_2$/MeOH = 99/1 to 85/15; *v*/*v*) to give **PC4** as a purple solid (53 mg, 10%).

$^1$H NMR (400 MHz, $(CD_3)_2SO$): δ 1.10 (t, *J* = 7.2 Hz, 12H), 2.27 (s, 3H), 3.34–3.42 (m, overlapped with $H_2O$ peak, 8H), 5.68 (s, 1H), 6.77 (d, *J* = 9.2 Hz, 4H), 7.10 (d, *J* = 8.0 Hz, 2H), 7.06–7.32 (br, 4H), 7.37 (d, *J* = 9.2 Hz, 4H), 7.47 (d, *J* = 8.0 Hz, 2H), 8.01 (d, *J* = 7.2 Hz, 4H). $^{13}$C NMR (100 MHz, $(CD_3)_2SO$): δ 12.32, 20.80, 43.86, 99.80, 111.67, 123.60, 125.52, 128.06, 130.50, 137.56, 138.13 (br), 145.82, 147.33, 148.55. HRMS (ESI positive mode) *m*/*z* calculated for $C_{31}H_{37}N_4$ $[M]^+$: 465.3013, found: 465.3007. Mp = 64.0–65.5°C.

Synthesis of **PC5**, **PC6**, **PC7**, and **PC8**.

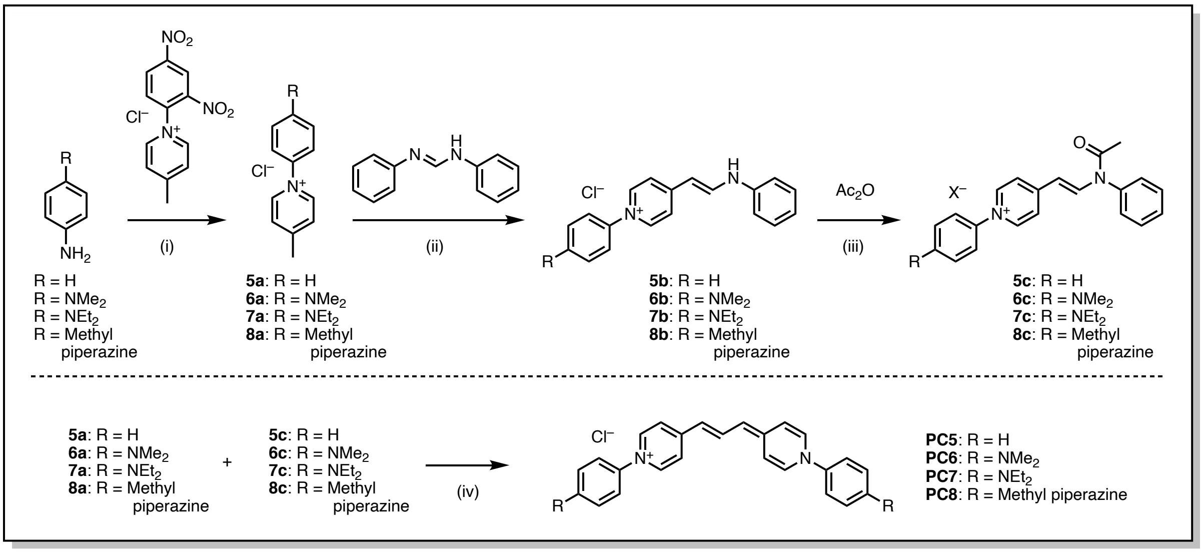


Scheme S2. (i) 1-(2,4-dinitrophenyl)-4-methylpyridin-1-ium chloride,[*] ethanol, reflux, 2.5–12 h; (ii) *N*,*N'*-diphenylformamidine, acetone/ethanol (= 1/1; *v*/*v*), reflux, 6–12 h; (iii) acetone/acetic anhydride (= 1/1; *v*/*v*), 60 °C, 4 h, $X^-$ = $Cl^-$ or $AcO^-$; (iv) $NEt_3$, $CH_2Cl_2$, reflux, 1 day.

#### 1-Phenyl-4-(2-(phenylamino)vinyl)pyridin-1-ium chloride (5b)

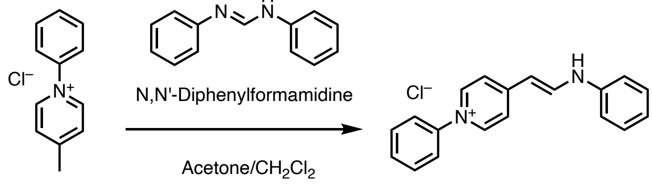


**5a** **5b**

A solution of 4-methyl-1-phenylpyridin-1-ium chloride (**5a**, 150 mg, 0.73 mmol) and *N*,*N'*-diphenylformamidine (286 mg, 1.46 mmol, 2.0 equiv.) in acetone (20 mL) and $CH_2Cl_2$ (20 mL) was stirred at reflux for 12 h. After cooling back to room temperature, the solvent was removed *in vacuo*. The resulting solid was subjected to column chromatography on silica gel (eluent: $CH_2Cl_2$/MeOH = 95/5 to 80/20; *v*/*v*) to give **5b** as a yellow powder (136 mg, 60%). The crude product was used to next reaction without further purification.

#### 1-phenyl-4-(2-(N-phenylacetamido)vinyl)pyridin-1-ium acetate (5c)

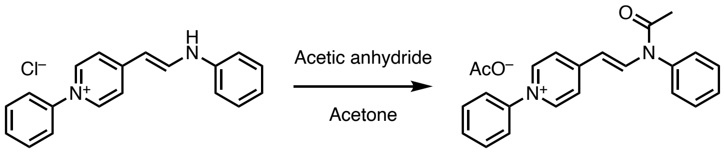


**5b** **5c**

To a solution of **5b** (120 mg, 0.39 mmol) in acetone (20 mL) was added acetic anhydride (20 mL), and the resultant mixture was stirred at 60 °C for 4 h. After cooling back to room temperature, acetone and acetic anhydride were removed *in vacuo*. The resulting solid was purified by column chromatography on silica gel (eluent: $CH_2Cl_2$/MeOH = 95/5 to 80/20; *v*/*v*) to give **5c** as a yellow powder (116 mg, 85%).

$^1$H NMR (400 MHz, $(CD_3)_2SO$): δ 1.88 (s, 3H), 2.05 (s, 3H), 5.53 (d, *J* = 14.4 Hz, 1H), 7.44 (d, *J* = 7.6 Hz, 2H), 7.54–7.74 (m, 6H), 7.76–7.84 (m, 4H), 8.16 (d, *J* = 7.2 Hz, 2H), 8.87 (d, *J* = 14.8 Hz, 1H), 8.96 (d, *J* = 6.8 Hz, 2H). $^{13}$C NMR (100 MHz, $(CD_3)_2SO$): δ 21.36, 23.23, 106.32, 122.33, 124.23, 128.56, 129.54, 130.10, 130.42, 130.62, 137.87, 140.51, 142.17, 143.10, 154.71, 169.78, 172.35. HRMS (ESI positive mode) *m*/*z* calculated for $C_{21}H_{19}N_2O$ $[M]^+$: 315.1492, found: 315.1486. Mp = 151.9–

153.2 °C.

**1-Phenyl-4-(3-(1-phenylpyridin-4(1*H*)-ylidene)prop-1-en-1-yl)pyridin-1-ium chloride (PC5)**

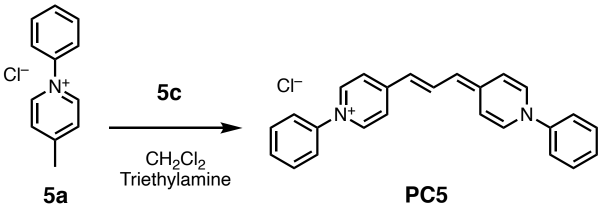


**5a** **PC5**

To a solution of **5a** (55 mg, 267 mmol) and **5c** (103 mg, 0.29 mmol, 1.1 equiv..) in $CH_2Cl_2$ (10 mL) was added triethylamine (2.0 mL) using a syringe under nitrogen atmosphere. The mixture was stirred at refluxing temperature for 1 day. After cooling back to room temperature, the solvent was removed *in vacuo*. The crude product was purified by column chromatography on silica gel (eluent: $CH_2Cl_2$/MeOH = 95/5 to 80/20; *v*/*v*). The blue fractions were collected and concentrated to give solid which was then washed by acetone once to give **PC5** as a blue solid (81.4 mg, 79%).

$^{1}$H NMR (600 MHz, $(CD_3)_2SO$): δ 6.00 (d, $J$ = 13.8 Hz, 2H), 7.32–7.36 (m, 4H), 7.52 (t, $J$ = 6.6 Hz, 2H), 7.56–7.66 (m, 8H), 8.03 (d, $J$ = 7.2 Hz, 4H), 8.27 (t, $J$ = 13.8 Hz, 1H). $^{13}$C NMR (100 MHz, $(CD_3)_2SO$): δ 111.10, 122.63, 128.59, 130.04, 137.85, 141.94, 142.25, 150.27. HRMS (ESI positive mode) *m*/*z* calculated for $C_{25}H_{21}N_2$ $[M]^+$: 349.1699, found: 349.1699. Mp = 194.2–195.0 °C.

**1-(4-(Dimethylamino)phenyl)-4-methylpyridin-1-ium chloride (6a)**

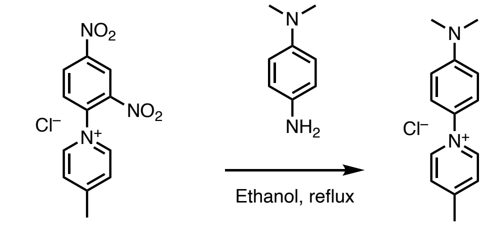


1-(2,4-dinitrophenyl)-4-methylpyridin-1-ium chloride

**6a**

To a solution of 1-(2,4-dinitrophenyl)-4-methylpyridin-1-ium chloride [(*)] (2.0 g, 6.76 mmol) in ethanol (50 mL) was added a solution of *N*,*N*-dimethyl-1,4-phenylenediamine (1.11 g, 8.12 mmol, 1.2 equiv.) in ethanol (10 mL) using a syringe at room temperature. Then the mixture was stirred at refluxing temperature for 4 h. After cooled back to room temperature, solvent was removed *in vacuo*. Then the resulting dark crude product was purified by column chromatography on silica gel (eluent: $CH_2Cl_2$/MeOH = 9/1 to 7/3; *v*/*v*) to give **6a** as a yellow solid (1.28 g, 76%).

$^{1}$H NMR (400 MHz, $(CD_3)_2SO$): δ 2.67 (s, 3H), 3.01 (s, 6H), 6.90 (d, $J$ = 9.6 Hz, 2H), 7.63 (d, $J$ = 9.6 Hz, 2H), 8.05 (d, $J$ = 6.4 Hz, 2H), 9.09 (d, $J$ = 6.8 Hz, 2H). $^{13}$C NMR (100 MHz, $(CD_3)_2SO$): δ 21.33, 112.18, 124.72, 128.33, 131.13, 142.98, 151.47, 158.47. HRMS (ESI positive mode) *m*/*z* calculated for $C_{14}H_{17}N_2$ $[M]^+$: 213.1386, found: 213.1386. Mp = 114.8–115.9 °C.

**1-(4-(Dimethylamino)phenyl)-4-(2-(phenylamino)vinyl)pyridin-1-ium chloride (6b)**

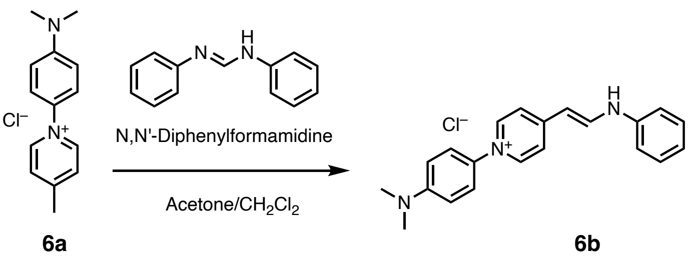


**6a** **6b**

A solution of **6a** (200 mg, 0.80 mmol) and *N*,*N'*-diphenylformamidine (316 mg, 161 mmol, 2.0 equiv.) in acetone (20 mL) and $CH_2Cl_2$ (20 mL) was stirred at refluxing temperature for 6 h. After cooling back to room temperature, the solvent was removed *in vacuo*. The obtained crude solid was subjected to column chromatography on silica gel (eluent: $CH_2Cl_2$/MeOH = 95/5 to 80/20; *v*/*v*) to give **6b** as an orange powder (162 mg, 57%).

$^1$H NMR (400 MHz, $(CD_3)_2SO$): δ 2.98 (s, 6H), 6.00 (d, *J* = 13.2 Hz, 1H), 6.87 (d, *J* = 9.6 Hz, 2H), 7.04 (t, *J* = 7.6 Hz, 1H), 7.30–7.44 (m, 4H), 7.53 (d, *J* = 9.2 Hz, 2H), 7.70–8.00 (br, 2H), 8.51 (d, *J* = 7.2 Hz, 2H), 8.64 (d, *J* = 13.2 Hz, 1H), 10.7–11.0 (br, 1H). $^{13}$C NMR (100 MHz, $(CD_3)_2SO$): δ 39.92, 99.09, 112.31, 116.09, 118.79, 122.96, 124.05, 129.50, 131.23, 140.48, 140.58, 143.52, 150.82, 154.50. HRMS (ESI positive mode) *m*/*z* calculated for $C_{21}H_{22}N_3$ $[M]^+$: 316.1808, found: 316.1808. Mp = 122.8–124.2 °C.

**1-(4-(Dimethylamino)phenyl)-4-(2-(*N*-phenylacetamido)vinyl)pyridin-1-ium chloride (6c)**

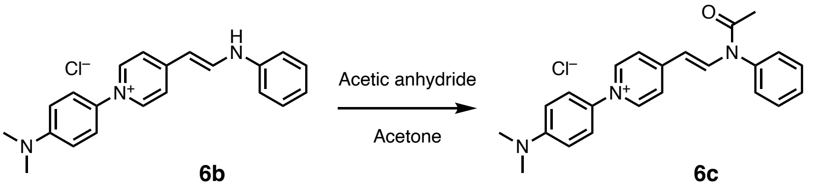

**6b** **6c**

To a solution of **6b** (120 mg, 341 mmol) in acetone (10 mL) was added acetic anhydride (10 mL) at room temperature under nitrogen atmosphere. Then the mixture was stirred at 60 °C for 4 h. After cooling back to room temperature, the solution was removed *in vacuo*. The resulting solid was subjected to column chromatography on silica gel (eluent: $CH_2Cl_2$/MeOH = 95/5 to 80/20; *v*/*v*) to give **6c** as an orange powder (102 mg, 76%).

$^1$H NMR (400 MHz, $(CD_3)_2SO$): δ 2.03 (s, 3H), 3.00 (s, 6H), 5.49 (d, *J* = 14.8 Hz, 1H), 6.87 (d, *J* = 9.2 Hz, 2H), 7.43 (d, *J* = 7.6 Hz, 2H), 7.52–76.8 (m, 5H), 8.07 (d, *J* = 7.2 Hz, 2H), 8.80 (d, *J* = 14.4 Hz, 1H), 8.87 (d, *J* = 7.2 Hz, 2H). $^{13}$C NMR (100 MHz, $(CD_3)_2SO$): δ 23.25, 106.47, 112.18, 122.42, 124.41, 128.60, 129.52, 130.43, 130.93, 137.96, 139.60, 142.36, 151.26, 153.02, 169.71 (two peaks were missing). HRMS (ESI positive mode) *m*/*z* calculated for $C_{23}H_{24}N_3O$ $[M]^+$: 358.1914, found: 358.1909. Mp = 109.0–111.1 °C.

**1-(4-(Dimethylamino)phenyl)-4-(3-(1-(4-(dimethylamino)phenyl)pyridin-4(1*H*)-ylidene)prop-1-en-1-yl)pyridin-1-ium chloride (PC6)**

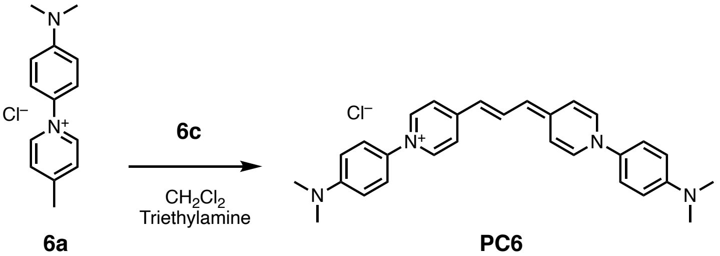

**6a** **PC6**

To a solution of **6a** (50.0 mg, 0.20 mmol) and **6c** (87.4 mg, 0.22 mmol, 1.1 equiv.) in $CH_2Cl_2$ (10 mL) was added triethylamine (2 mL) using a syringe under nitrogen atmosphere. The mixture was stirred at refluxing temperature for 1 day. After cooling back to room temperature, the solvent was removed *in vacuo*. The resulting solid was purified by column chromatography on silica gel (eluent: $CH_2Cl_2$/MeOH = 95/5 to 80/20; *v*/*v*). The blue fractions were collected and concentrated. The obtained solid was dissolved in acetone to which ether was added to form the precipitation, which was then filtered to give **PC6** as a blue solid (28.8 mg, 30%).

$^1$H NMR (600 MHz, $(CD_3)_2SO$): δ 2.96 (s, 12H), 5.88 (d, *J* = 13.2 Hz, 2H), 6.83 (d, *J* = 8.4 Hz, 4H),

7.42 (d, $J$ = 9.0 Hz, 4H), 7.95 (d, $J$ = 7.2 Hz, 4H), 8.23 (t, $J$ = 13.2 Hz, 1H). $^{1}$H NMR (600 MHz, $(CD_3)_2SO$, 60 °C): δ 2.97 (s, 12H), 5.90 (d, $J$ = 13.2 Hz, 2H), 6.84 (d, $J$ = 8.4 Hz, 4H), 7.24 (br, 4H), 7.40 (d, $J$ = 8.4 Hz, 4H), 7.9 (d, $J$ = 7.2 Hz, 4H), 8.17 (t, $J$ = 13.2 Hz, 1H). $^{13}$C NMR (150 MHz, $(CD_3)_2SO$, 80 °C): δ 109.73, 112.30, 116.38, 122.84, 131.27, 137.37, 140.42, 149.19, 150.05. HRMS (ESI positive mode) *m/z* calculated for $C_{29}H_{31}N_4$ $[M]^+$: 435.2543, found: 435.2552. Mp = 143.6–174.7 °C.

**1-(4-(Diethylamino)phenyl)-4-methylpyridin-1-ium chloride (7a)**

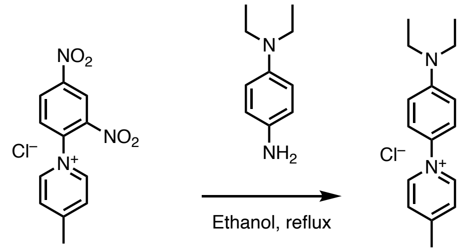


1-(2,4-dinitrophenyl)-4 -methylpyridin-1-ium chloride

**7a**

To a solution of 1-(2,4-Dinitrophenyl)-4-methylpyridin-1-ium chloride (5.0 g, 17 mmol) in ethanol (70 mL) was added a solution of *N*,*N*-diethyl-1,4-phenylenediamine (3.33 g, 20.3 mmol, 1.2 equiv..) in ethanol (20 mL) using a syringe at room temperature. Then the mixture was stirred at refluxing temperature for 12 h. After cooled to room temperature, solvent was removed *in vacuo*. The obtained dark oil was purified by column chromatography on silica gel (eluent: $CH_2Cl_2$/MeOH = 90/10 to 70/30; *v*/*v*) to give **7a** as a yellow solid (3.86 g, 82%).

$^{1}$H NMR (400 MHz, $(CD_3)_2SO$): δ 1.11 (t, $J$ = 7.4 Hz, 6H), 2.66 (s, 3H), 3.43 (q, $J$ = 7.1 Hz, 4H), 6.85 (d, $J$ = 9.2 Hz, 2H), 7.59 (d, $J$ = 9.2 Hz, 2H), 8.04 (d, $J$ = 7.2 Hz, 2H), 9.08 (d, $J$ = 6.8 Hz, 2H). $^{13}$C NMR (100 MHz, $(CD_3)_2SO$): δ 12.25, 21.31, 43.93, 111.42, 125.03, 128.31, 130.27, 142.87, 148.72, 158.27. HRMS (ESI positive mode) *m/z* calculated for $C_{16}H_{21}N_2$ $[M]^+$: 241.1699, found: 241.1697. Mp = 138.5–139.9 °C

**1-(4-(Diethylamino)phenyl)-4-(2-(phenylamino)vinyl)pyridin-1-ium chloride (7b)**

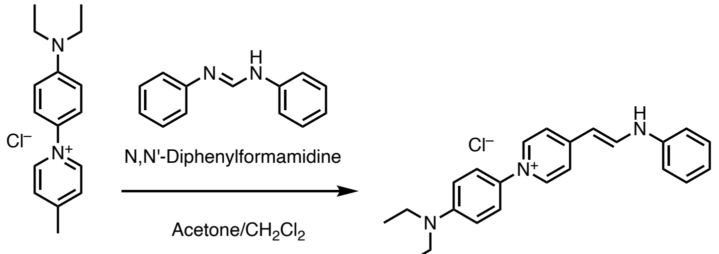


**7a** **7b**

A solution of **7a** (300 mg, 1.08 mmol) and *N*,*N*'-diphenylformamidine (425 mg, 2.17 mmol, 2.0 equiv.) in acetone (30 mL) and $CH_2Cl_2$ (30 mL) was stirred at refluxing temperature for 6 h. After cooling back to room temperature, the solvent was removed. The resulting red solid was subjected to column chromatography on silica gel (eluent $CH_2Cl_2$/MeOH = 95/5 to 80/20; *v*/*v*) to give **7b** as a red powder (287 mg, 70%).

$^{1}$H NMR (400 MHz, $(CD_3)_2SO$): δ 1.11 (t, $J$ = 7.0 Hz, 6H), 3.41 (q, $J$ = 7.2 Hz, 4H), 5.96 (d, $J$ = 13.2 Hz, 1H), 6.80 (d, $J$ = 9.2 Hz, 2H), 7.03 (t, $J$ = 7.8 Hz, 1H), 7.30–7.42 (m, 4H), 7.48 (d, $J$ = 9.2 Hz, 2H), 7.60–7.90 (br, 2H), 8.44 (d, $J$ = 7.2 Hz, 2H), 8.62 (d, $J$ = 12.8 Hz, 1H), 10.5–11.3 (br, 1H). $^{13}$C NMR (100 MHz, $(CD_3)_2SO$): δ 12.27, 43.89, 99.22, 111.51, 116.26, 118.45 (br), 122.91, 124.30, 129.45, 130.37, 140.27, 141.11, 143.95, 147.89, 153.94. HRMS (ESI positive mode) *m/z* calculated for $C_{23}H_{26}N_3$ $[M]^+$: 344.2121, found: 344.2118.

**1-(4-(Diethylamino)phenyl)-4-(2-(*N*-phenylacetamido)vinyl)pyridin-1-ium chloride (7c)**

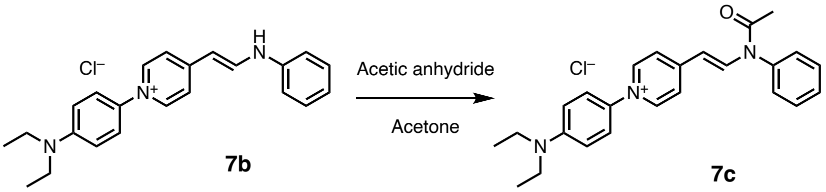


To a solution of **7b** (250 mg, 0.66 mmol) in acetone (10 mL) was added acetic anhydride (20 mL) using a syringe at room temperature under nitrogen atmosphere. The mixed solution was stirred at 60 °C for 4 h under nitrogen atmosphere. Then, the solvent was removed *in vacuo*. The resulting solid was purified by column chromatography on silica gel (eluent: $CH_2Cl_2$/MeOH = 95/5 to 80/20; *v*/*v*) to give **7c** as a red powder (131 mg, 47%).

$^1$H NMR (400 MHz, $(CD_3)_2SO$): δ 1.10 (t, *J* = 7.2 Hz, 6H), 2.03 (s, 3H), 3.41 (q, *J* = 7.2 Hz, 4H), 5.48 (d, *J* = 14.4 Hz, 1H), 6.82 (d, *J* = 9.6 Hz, 2H), 7.43 (d, *J* = 7.2 Hz, 2H), 7.50–7.68 (m, 5H), 8.06 (d, *J* = 6.8 Hz, 2H), 8.79 (d, *J* = 14.8 Hz, 1H), 8.84 (d, *J* = 7.2 Hz, 2H). $^{13}$C NMR (100 MHz, $(CD_3)_2SO$): δ 12.26, 23.24, 43.92, 106.48, 111.44, 122.42, 124.72, 128.60, 129.51, 130.06, 130.42, 137.96, 142.26, 148.51, 152.82, 167.70. One carbon peak was not found. HRMS (ESI positive mode) *m*/*z* calculated for $C_{25}H_{28}N_3O$ $[M]^+$: 386.2227, found: 386.2225. Mp = 243.3–244.6 °C.

**1-(4-(Diethylamino)phenyl)-4-(3-(1-(4-(diethylamino)phenyl)pyridin-4(1*H*)-ylidene)prop-1-en-1-yl)pyridin-1-ium chloride (PC7)**

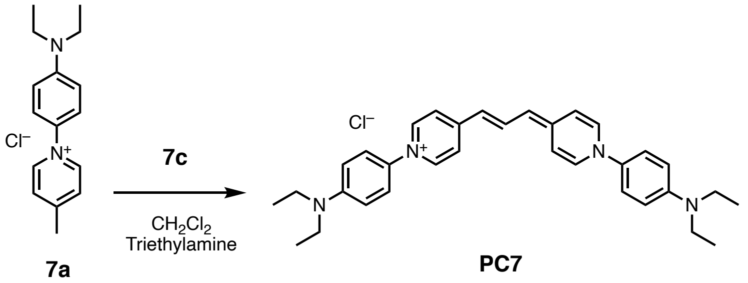


To a solution of **7a** (70 mg, 0.25 mmol) and **7c** (112 mg, 0.27 mmol, 1.05 equiv.) in $CH_2Cl_2$ (10 mL) was added triethylamine (1 mL) using a syringe under nitrogen atmosphere. The mixture was stirred at refluxing temperature for 1 day. After cooling back to room temperature, the solvent was removed *in vacuo*. The resulting dark solid was subjected to column chromatography on silica gel (eluent: $CH_2Cl_2$/MeOH = 95/5 to 80/20; *v*/*v*). The blue fractions were collected and concentrated. The obtained solid was dissolved in acetone, and ether was added to the solution to form the precipitation, which was then filtered to give **PC7** as a blue solid (32.7 mg, 25%).

$^1$H NMR (600 MHz, $(CD_3)_2SO$): δ 1.10 (t, *J* = 6.6 Hz, 12H), 3.34–3.42 (m, 8H), 5.86 (d, *J* = 13.2 Hz, 2H), 6.77 (d, *J* = 8.4 Hz, 4H), 7.37 (d, *J* = 9.0 Hz, 4H), 7.95 (distorted s, 4H), 8.22 (t, *J* = 13.8 Hz, 1H). $^{13}$C NMR (150 MHz, $(CD_3)_2SO$): δ 12.30, 43.86, 109.87, 111.66, 123.55, 130.63, 137.74, 140.72, 147.24, 149.38. One carbon peak was not found. HRMS (ESI positive mode) *m*/*z* calculated for $C_{33}H_{39}N_4$ $[M]^+$: 491.3169, found: 491.3170. Mp = 155.1–156.5 °C.

**4-Methyl-1-(4-(4-methylpiperazin-1-yl)phenyl)pyridin-1-ium chloride (8a)**

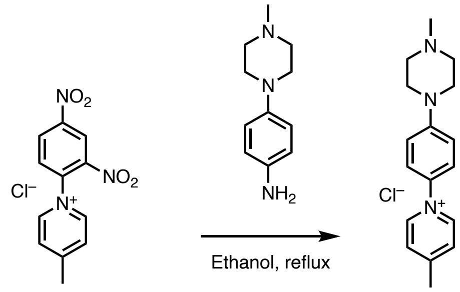


1-(2,4-dinitrophenyl)-4-methylpyridin-1-ium chloride

8a

To a solution of 1-(2,4-dinitrophenyl)-4-methylpyridin-1-ium chloride (3.0 g, 10 mmol) in ethanol (50 mL) was added a solution of 4-(4-methylpiperazin-1-yl)aniline (2.04 g, 1.1 mmol, 1.1 equiv.) in ethanol (10 mL) using a syringe at room temperature. The reaction mixture was stirred 2.5 h. After cooling back to room temperature, the solvent was removed *in vacuo*. Then the resulting dark oil was subjected to column chromatography on silica gel (amino silica gel, eluent: $CH_2Cl_2$/MeOH = 100/1 to 95/5; *v*/*v*) to give **8a** as an orange solid (855 mg, 28%).

$^1$H NMR (400 MHz, $(CD_3)_2SO$): δ 2.22 (s, 3H), 2.38–2.48 (m, 4H), 2.68 (s, 3H), 3.23–3.32 (m, 4H), 7.18 (d, *J* = 8.8 Hz, 2H), 7.66 (d, *J* = 9.2 Hz, 2H), 8.07 (d, *J* = 6.4 Hz, 2H), 9.12 (d, *J* = 5.6 Hz, 2H). $^{13}$C NMR (100 MHz, $(CD_3)_2SO$): δ 21.36, 45.71, 47.15, 54.26, 115.03, 124.84, 128.31, 132.93, 143.12, 152.19, 158.91. HRMS (ESI positive mode) *m*/*z* calculated for $C_{17}H_{22}N_3$ $[M]^+$: 268.1808, found: 268.1807. Mp = 74.1–75.6 °C.

#### 1-(4-(4-Methylpiperazin-1-yl)phenyl)-4-(2-(phenylamino)vinyl)pyridin-1-ium chloride(8b)

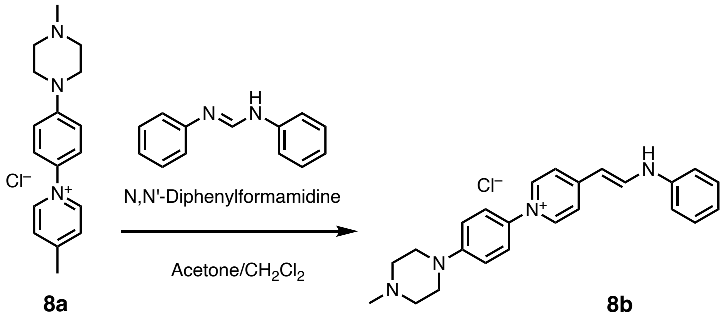


8a 8b

A solution of **8a** (300 mg, 0.99 mmol) and *N*,*N*'-diphenylformamidine (387 mg, 1.97 mmol, 2.0 equiv.) in acetone (25 mL) and $CH_2Cl_2$ (25 mL) was stirred at refluxing temperature for 12 h. After cooling back to room temperature, the solvent was removed *in vacuo*. The resulting solid was subjected to column chromatography on silica gel (amino silica gel, eluent: $CH_2Cl_2$/MeOH = 100/0 to 90/10; *v*/*v*) to give **8b** (208 mg, 52%) as an orange powder.

$^1$H NMR (400 MHz, $(CD_3)_2SO$): δ 2.21 (s, 3H), 2.38–2.47 (m, 4H), 3.08–3.20 (m, 4H), 5.35 (d, *J* = 10.0 Hz, 1H), 6.21 (br s, 1H), 6.90–7.10 (m, 6H), 7.20–7.40 (m, 7H), 8.40 (d, *J* = 10.0 Hz, 1H). $^{13}$C NMR (100 MHz, $(CD_3)_2SO$): δ 45.75, 47.92, 54.46, 103.01, 109.26, 114.96, 115.91, 120.59, 121.78, 122.89, 128.81, 133.89 (br), 134.22, 144.32, 149.61, 153.83, 156.04. One extra carbon peak was observed. HRMS (ESI positive mode) *m*/*z* calculated for $C_{24}H_{27}N_4$ $[M]^+$: 371.2230, found: 371.2225.

#### 1-(4-(4-methylpiperazin-1-yl)phenyl)-4-(2-(N-phenylacetamido)vinyl)pyridin-1-ium chloride (8c)

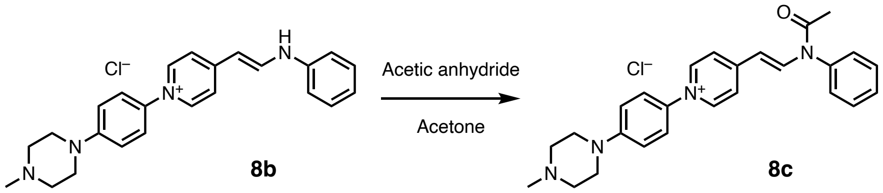


To a solution of **8b** (150 mg, 0.37 mmol) in acetone (20 mL) was added acetic anhydride (20 mL) using a syringe at room temperature under nitrogen atmosphere. Then the mixture was stirred at 60 °C for 4 h. After cooling back to room temperature, the solution was removed *in vacuo*. The resulting solid was purified by column chromatography on silica gel (amino silica gel, eluent: $CH_2Cl_2$/MeOH = 100/0 to 90/10; *v*/*v*) to give **8c** as a red powder (128 mg, 77%). The obtained product was used to the next reaction without further purification.

**1-(4-(4-methylpiperazin-1-yl)phenyl)-4-(3-(1-(4-(4-methylpiperazin-1-yl)phenyl)pyridin-4(1*H*)-ylidene)prop-1-en-1-yl)pyridin-1-ium chloride (PC8)**

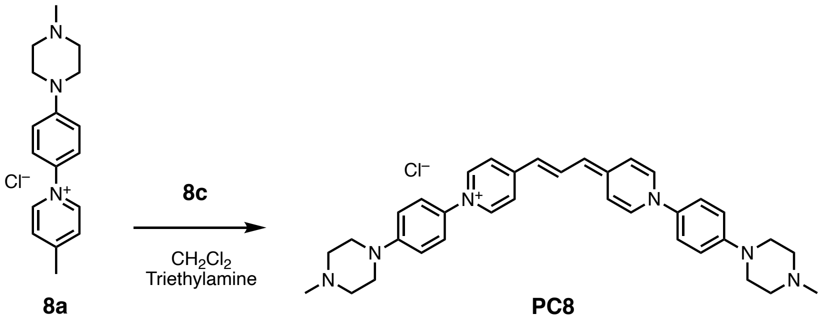


To a solution of **8a** (70 mg, 0.23 mmol) and **8c** (113 mg, 0.25 mmol, 1.1 equiv.) in $CH_2Cl_2$ (10 mL) was added triethylamine (1 mL) using a syringe under nitrogen atmosphere. The reaction mixture was stirred at refluxing temperature for 1 day. After cooling back to room temperature, the solvent was removed *in vacuo*. The resulting solid was subjected to column chromatography on amino silica gel (amino silica gel, eluent: $CH_2Cl_2$/MeOH = 100/0 to 90/10; *v*/*v*). The blue fractions were collected and concentrated. Then the obtained solid was recrystallized from methanol to give **PC8** as a glossy dark green powder (92 mg, 69%).

$^1$H NMR (400 MHz, $(CD_3)_2SO$): δ 2.22 (s, 6H), 2.36–2.47 (m, 8H), 3.14–3.27 (m, 8H), 5.89 (d, *J* = 13.6 Hz, 2H), 7.08 (d, *J* = 9.2 Hz, 4H), 7.45 (d, *J* = 8.8 Hz, 4H), 7.99 (d, *J* = 7.6 Hz, 4H), 8.26 (t, *J* = 13.6 Hz, 1H). $^{13}$C NMR (100 MHz, $(CD_3)_2SO$): δ 45.23, 47.32, 54.02, 110.00, 115.21, 116.41, 122.86, 133.07, 137.34, 140.67, 149.34, 150.53. HRMS (ESI positive mode) *m*/*z* calculated for $C_{35}H_{41}N_6$ $[M]^+$: 545.3387, found: 545.3389. Mp = 65.5–67.0 °C.

**3. Figure S1 to S9**

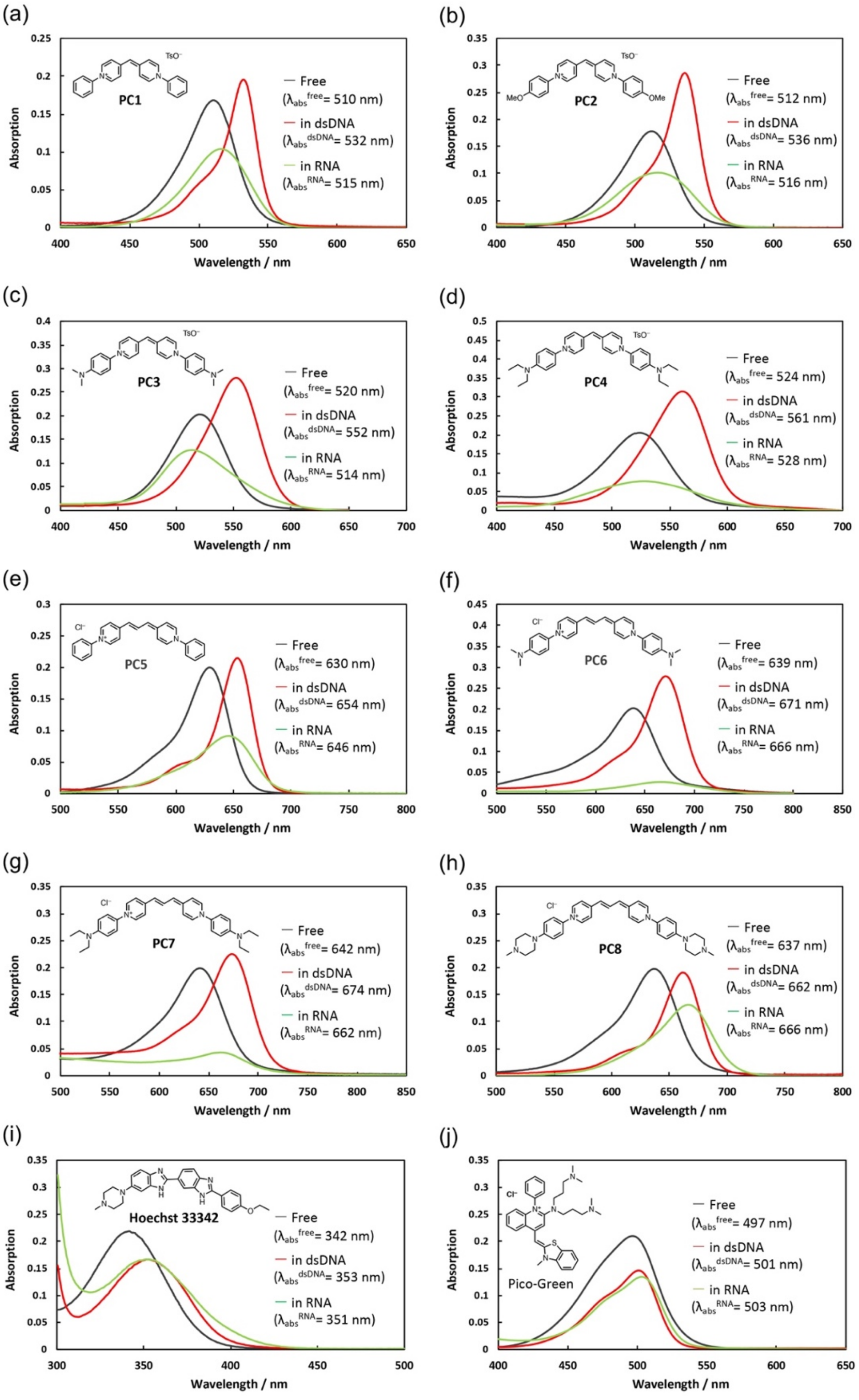


**Figure S1. Absorption spectra of PC derivatives and two popular commercialized DNA probes in EDTA solution upon mixing excess of dsDNA and RNA.** Black lines, red lines, and green lines represent free state, DNA-complexed state, and RNA-complexed state absorption, respectively.

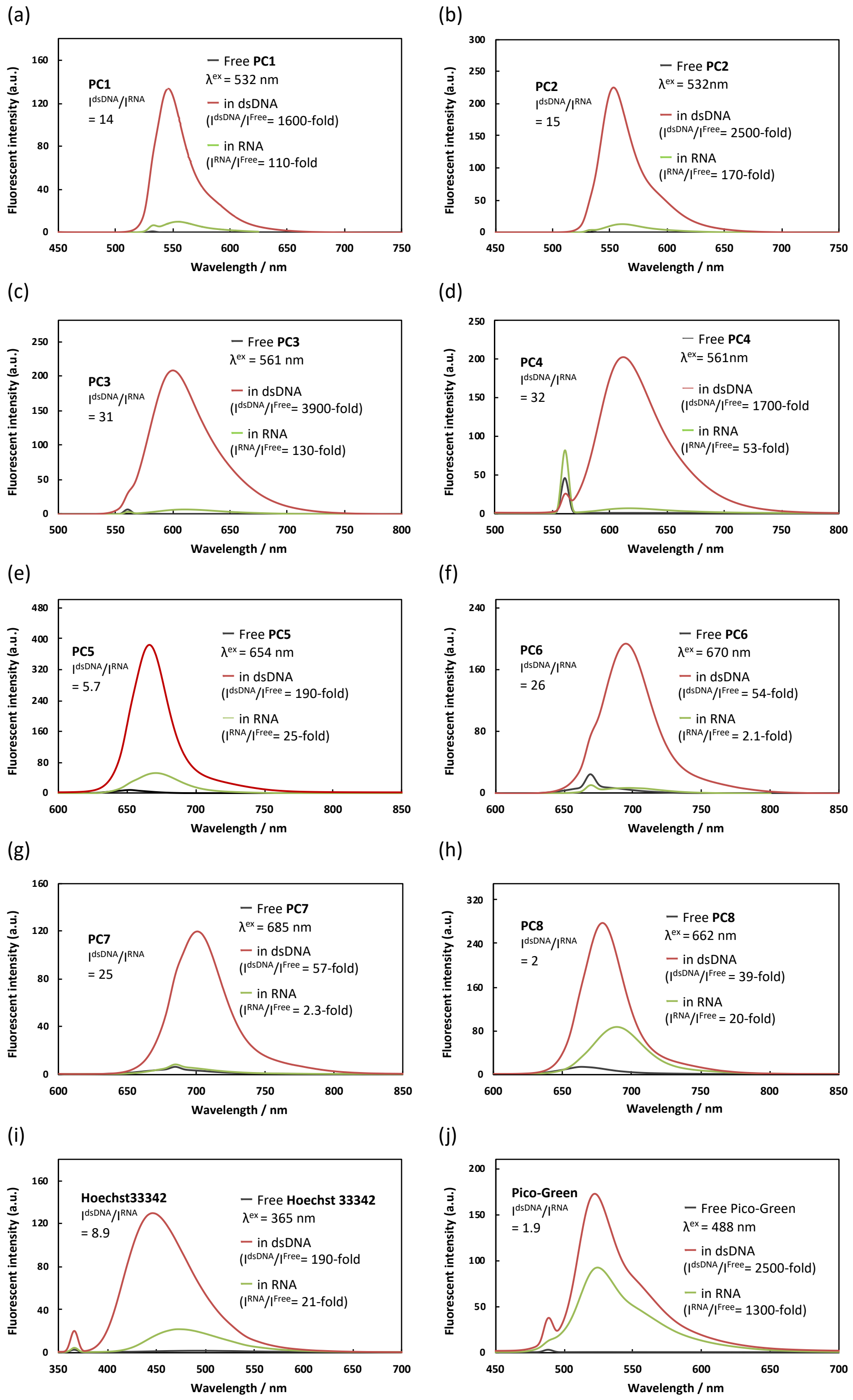


**Fig. S2. Fluorescence spectra of PC derivatives and two popular commercialized DNA probes in EDTA solution upon mixing excess of dsDNA and RNA.** Black lines indicate the fluorescence spectra of noted dyes in EDTA solution. Change of fluorescence spectra and increase of fluorescence intensity upon mixing excess of dsDNA (red-lines) and RNA (green-lines).

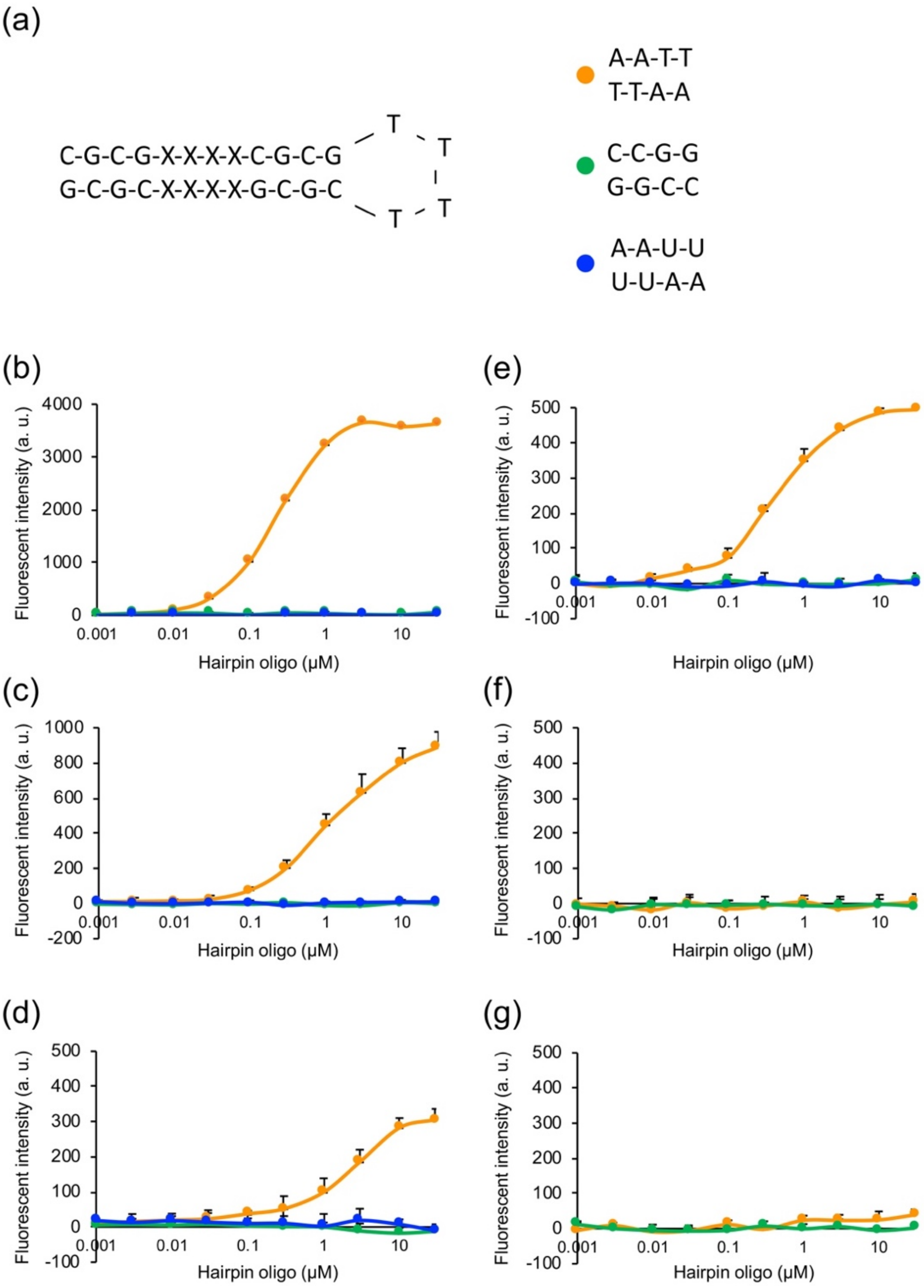


**Fig. S3. Titration of 100 nM N-aryl PC dye derivatives with various concentration of hairpin DNA and RNA. (a)** The sequences of hairpin oligo DNA (orange and green) and RNA (blue) **(b-f)** The titration curve of 100 nM PC dyes, (b) **PC1** emission at 547 nm excited at 532 nm, (c) **PC3** emission at 600 nm excited at 552 nm, (d) **PC4** emission at 612 nm excited at 561 nm, (e) **PC5** emission at 669 nm excited at 654 nm, (f) **PC6** emission at 695 nm excited at 671 nm, (g) **PC7** emission at 700 nm excited at 674 nm. Error bars represent mean ± s.d. from three independent replicates.

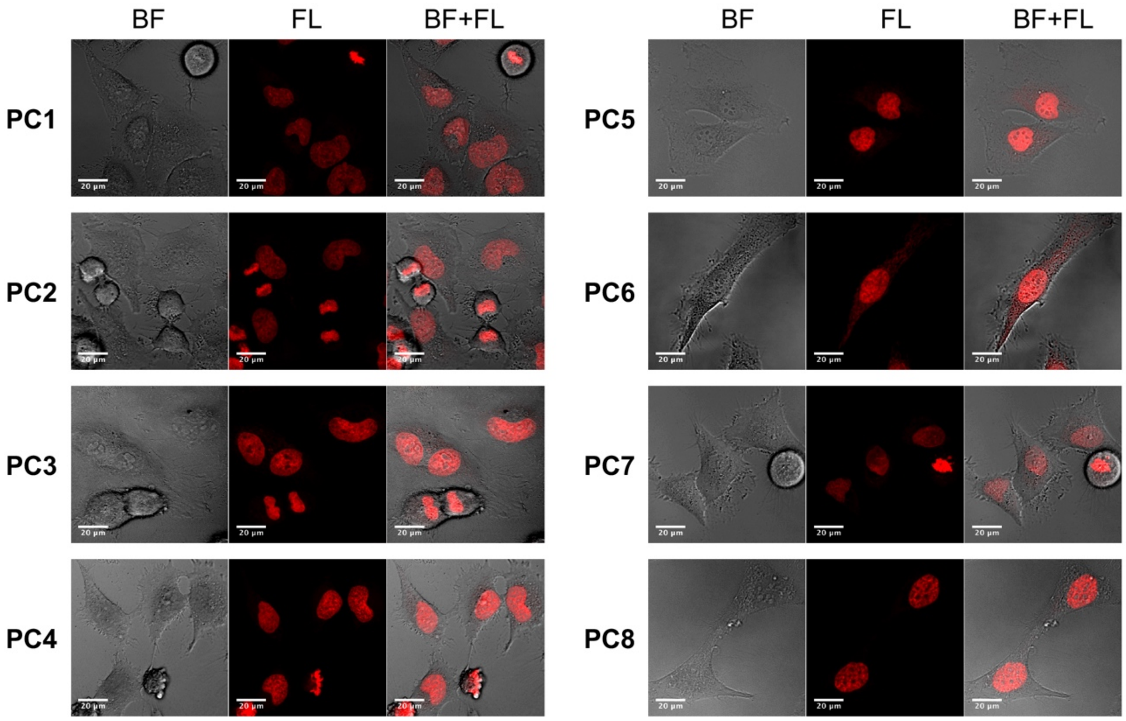


**Fig. S4. N-aryl PC derivatives stain nucleus in living HeLa cells.** HeLa cells were stained with each dye at the concentration of 1 μM. The fluorescence image stained with **PC1** were obtained emission spectrum between 517-693 nm excited with 514 nm. The images stained with **PC2-PC4** were obtained emission spectrum between 570-693 nm excited with 560 nm. The images stained with **PC5-PC8** were obtained the emission spectrum between 640-693 nm excited with 633 nm. Fluorescent images are maximum z-projections of total planes (1 μm intervals). BF; bright-field, FL; fluorescence, BF+FL; overalaid image of BF and FL.

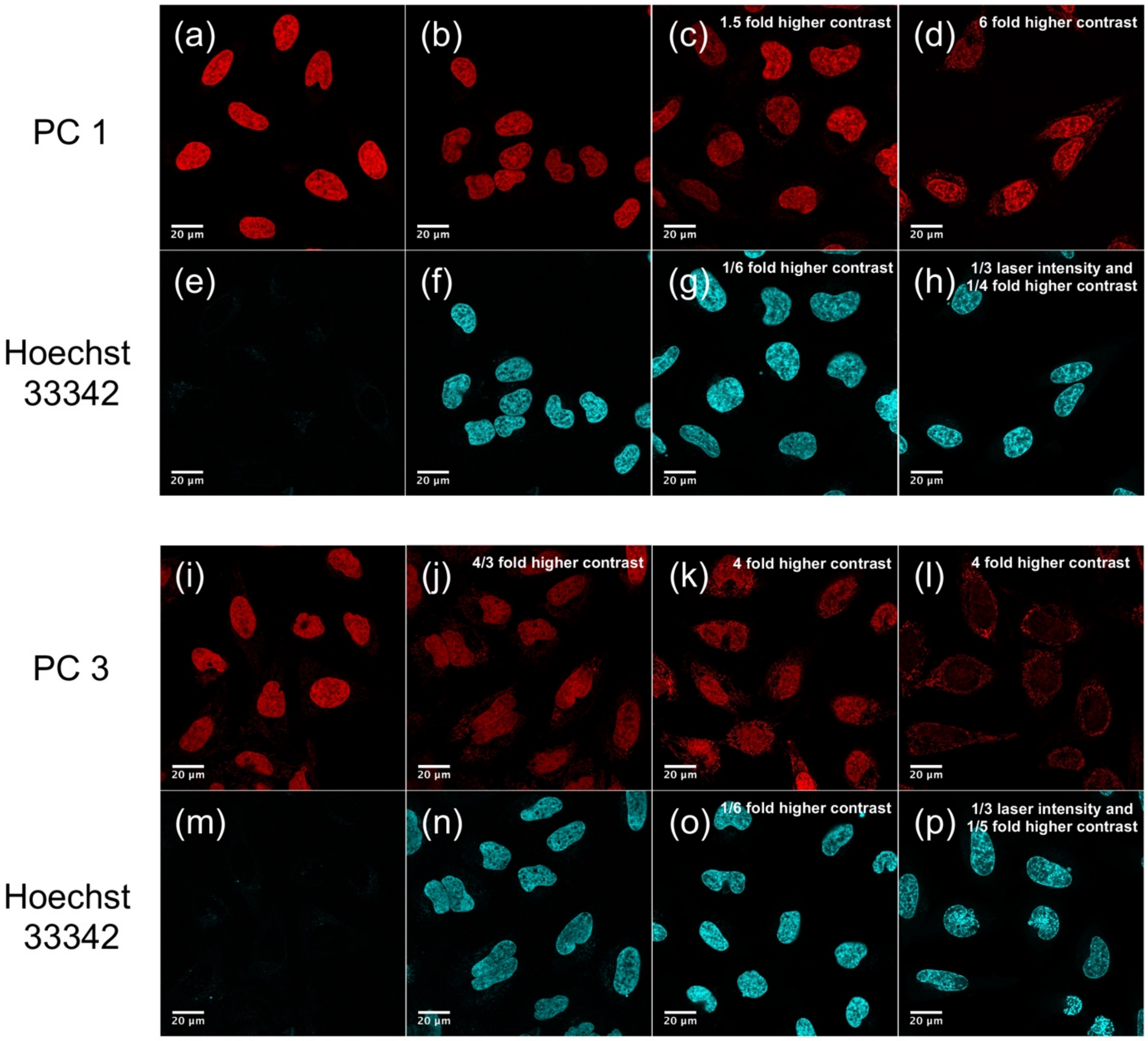


**Fig. S5. Co-staining of N-aryl PC dyes with Hoechst 33342 in living HeLa cells. (a-h)** HeLa cells were co-stained with 100 nM **PC1** with 0 nM (a, e), 100 nM (b, f), 1 μM (c, g), and 3 μM (d, h) Hoechst 33342. **(i-p)** HeLa cells were co-stained with 100 nM **PC3** with Hoechst 33342; 0 nM (i, m), 100 nM (j, n), 1 μM (k, o), 3 μM (l, p). Note that N-aryl PC dyes are excluded from nucleus by Hoechst 33342 in dose dependent manner.

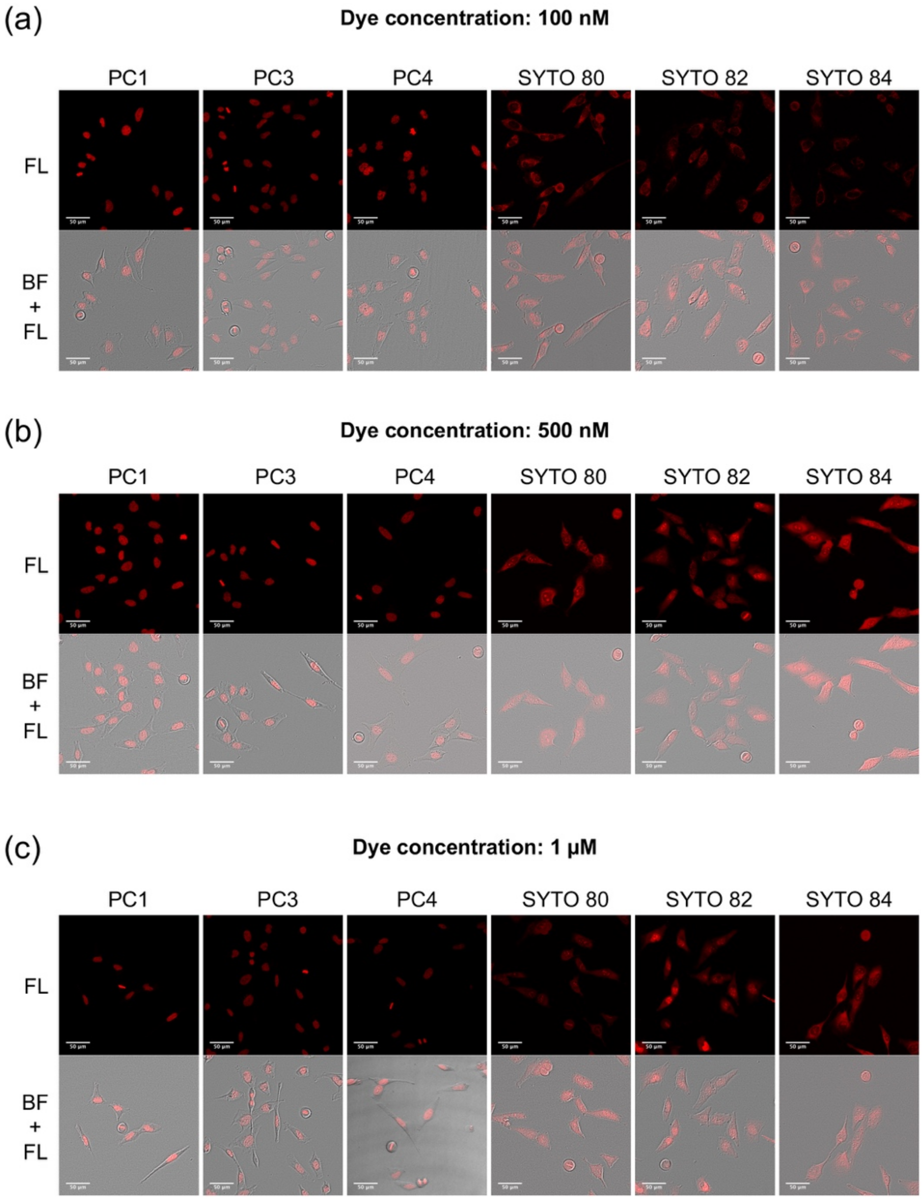


**Fig. S6. Comparison of nuclear DNA stain between N-aryl PC dyes and SYTO orange dyes in living HeLa cells.** HeLa cells were stained with each dye at 100 nM (a), 500 nM (b) and 1 µM (c). The excitation laser lines were used into alignment with absorption peak wavelength of each dye (532 nm for **PC1**, 552 nm for **PC3**, 561 nm for **PC4**, 532 nm for **SYTO 80**, 543 nm for **SYTO 82**, 561 nm for **SYTO 84**) and those emission spectra were collected in 540-670 nm, 560-670 nm, 570-670 nm, 540-670 nm, 550-670 nm, 570-670 nm, respectively. Note that N-aryl PC dyes specifically stain cell nucleus and chromosome in all concentration tested, whereas SYTO series stain cytoplasm and nucleolus instead of specific labelling of nucleus. BF; bright-field, FL; fluorescence, BF+FL; overalaid image of BF and FL. Selected images in (a) and (b) were also used in Fig. 2a and Fig.2b, respectively.

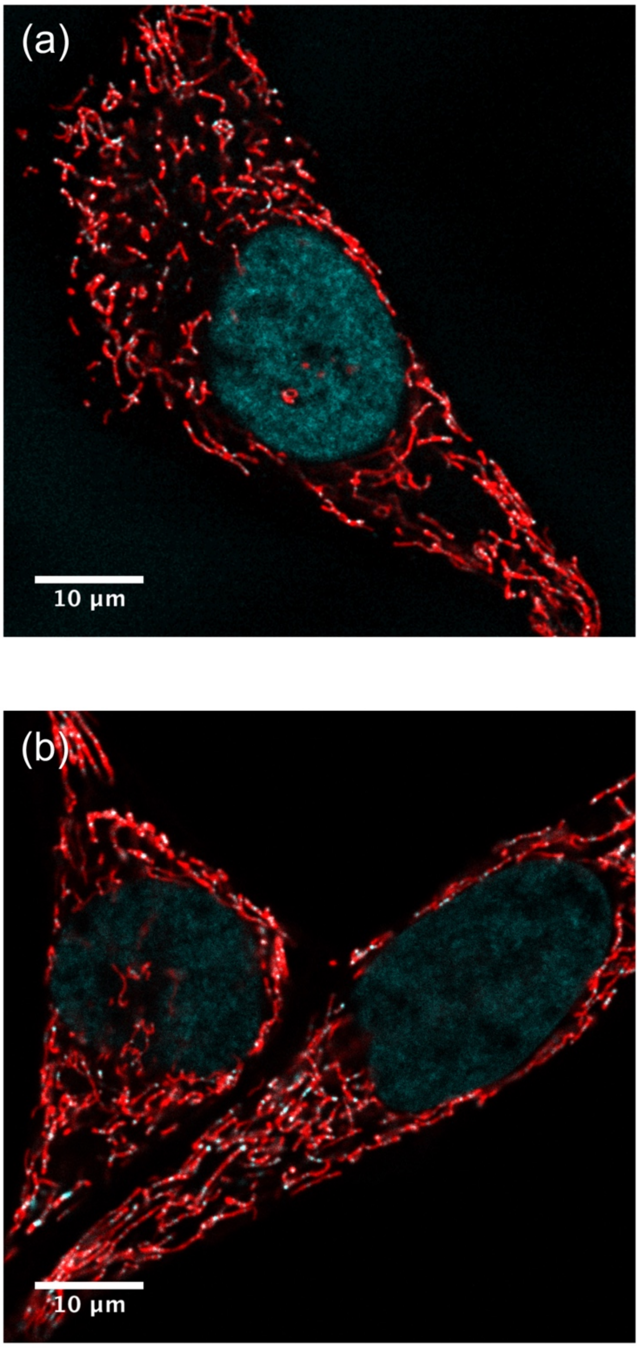


**Fig. S7. Co-staining of N-aryl PC dyes with MitoTracker Deep Red.** (a) Overlaid image of **PC1** (cyan) and MitoTracker Deep red (red). (b) Overlaid image of **PC3** (cyan) and MitoTracker Deep Red (red). Note that the cytoplasmic spots of PC dyes are localized in mitocondorial tubes.

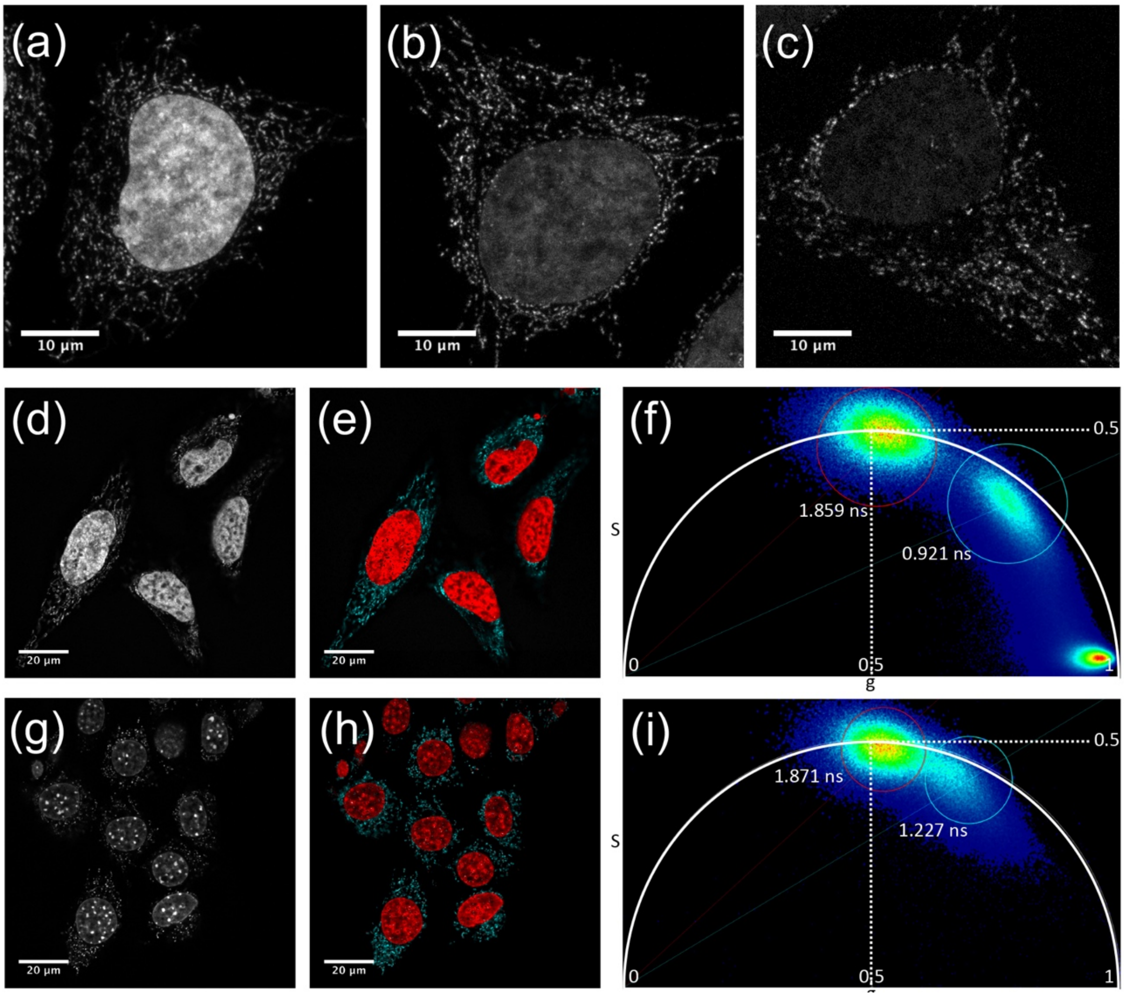


**Fig. S8. Discrimination between nuclear DNA and mt-DNA with fluorescence lifetime of PC3.** Concentration dependence of staining pattern with **PC3**. (a) 10 nM, (b) 1 nM, (c) 100 pM. The images are maximum z-projections of total planes (0.3 µm intervals). **(d-l)** Fluorescent intensity images (d, g) and FLIM based separation images of nuclear DNA and mitochondrial DNA (e, h) by phasor plot analysis (f, I). The pseudo colors of (e, h) is correspond to the colors of circles in (f, I). HeLa cells (d-f) and NIH3T3 (g-i) were stained with 10 nM and 30 nM **PC3**, respectively and the fluorescent spectrum were collected between 570-650 nm excited at 561 nm.

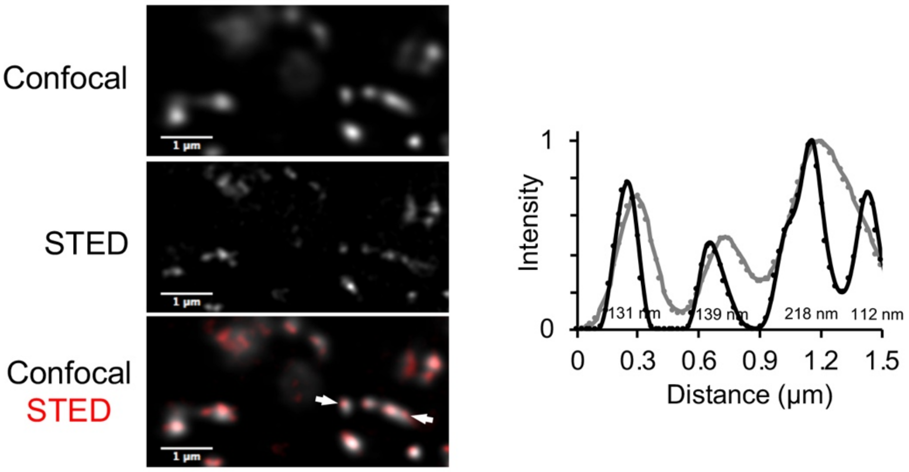


**Fig. S9. Comparison of confocal and STED-FlIM imaging in living NIH3T3 cells stained with PC3.** NIH/3T3 cells were stained with **PC3** at 30 nM concentration. An example of normalized fluorescence intensity profile obtained from the region between arrows. Line profiles in STED and confocal image are shown in black and gray, respectively. FWHM values estimated by fitting with a Gaussian function are also indicated in the black line profile.

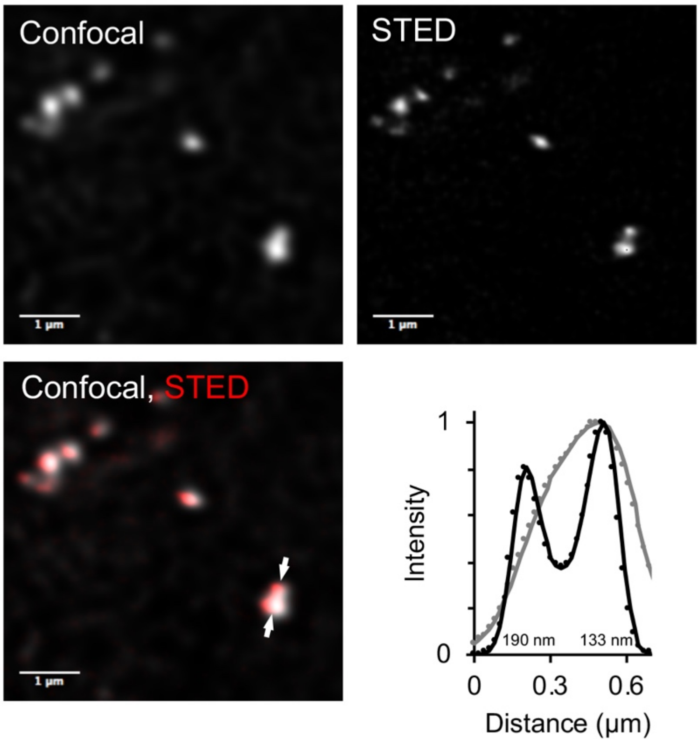


**Fig. S10. Comparison of confocal and STED-FlIM imaging in living Arabidopsis root cells stained with PC3.** Arabidopsis root cells were stained with **PC3** at 300 nM concentration. An example of normalized fluorescence intensity profile obtained from the region between arrows. F Line profiles in STED and confocal image are shown in black and gray, respectively. FWHM values estimated by fitting with a Gaussian function are also indicated in the black line profile.

### 4. Caption for movies S1 to S6

**Movie S1. Staining nuclear DNA in Arabidopsis leaf cells with PC1**
Arabidopsis leaf cells were stained with 1 μM **PC1**. Note that **PC1** stains nuclear DNA in mesophyll cells as well as epidermal cells including stomata.

**Movie S2. Staining nuclear DNA in Arabidopsis leaf cells with PC1**
Arabidopsis leaf cells were stained with 1 μM **PC3**. Note that **PC3** stains nuclear DNA in mesophyll cells as well as epidermal cells including stomata.

**Movie S3. Time-lapse analysis of Arabidopsis root and root hairs with 1 μM PC1.** Arabidopsis root stained with 1 μM **PC1** was observed every 5 min excited with 488 nm and the emission spectrum was collected through band-pass filter BP525/50. The fluorescence images are maximum z-projections of 20 planes (4.3-μm intervals). On the right side, combined images of the **PC1** and bright-field images. Note that the growing root hairs appear after the root tip moved through the image area.

**Movie S4. Comparison of light penetration for CLSM and 2PEM in Arabidopsis root tip stained with 1 μM PC1.**
Selected frames every 10 μm are shown in Fig. 2e

**Movie S5. Time-lapse observation by two-photon microscopy excited with 1000 nm in Arabidopsis root stained with PC1.**
Root tip stained with 5 μM **PC1** was observed every 2 min with z-sectioning (50 frames at 2 μm steps). The images are maximum z-projections of middle 25 planes. Selected frames every 10 min between 40 – 80 min are shown in Fig. 2e

**Movie S6. Discrimination between nuclear DNA, mt-DNA, and chl-DNA with fluorescence lifetime of PC-1.**
Arabidopsis leaf cells were stained with 300 nM **PC1**. The average fluorescent life times of mt-DNA (0.882 ns), chl-DNA (0.562 ns) and nuclear DNA (1.199 ns) are indicated in cyan, yellow and mazenda, respectively. The Maximum projected image was shown in Fig. 3k.

### 5. NMR charts

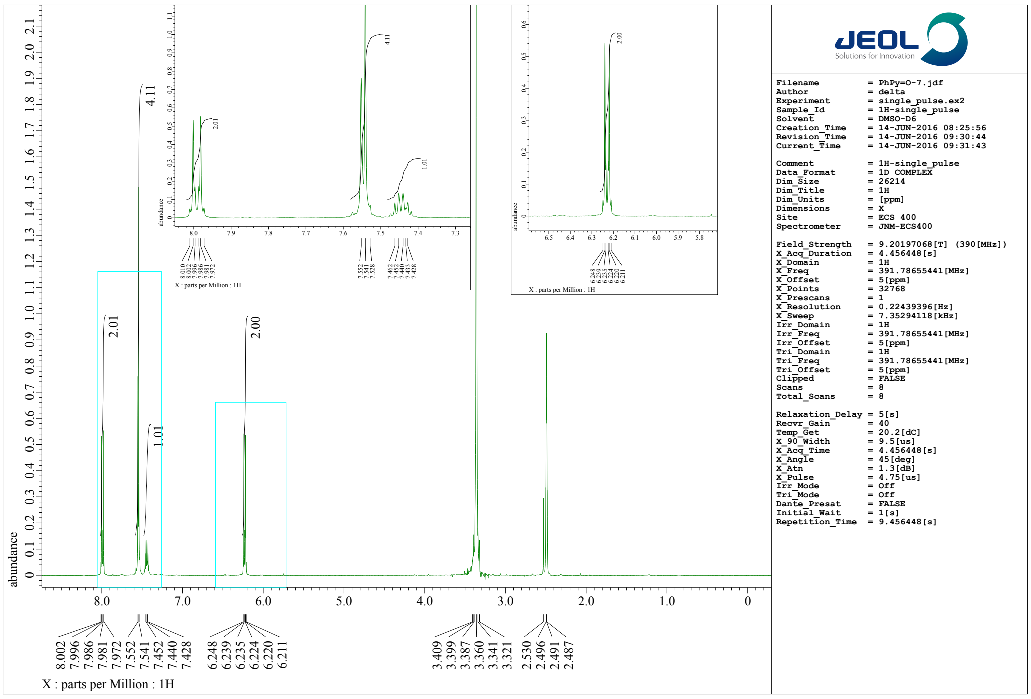


**Fig. S11-a1**. $^{1}$H-NMR (400 MHz) spectrum of **1a** in dimethyl dimethyl sulfoxide-$d_6$ ($d$-DMSO)

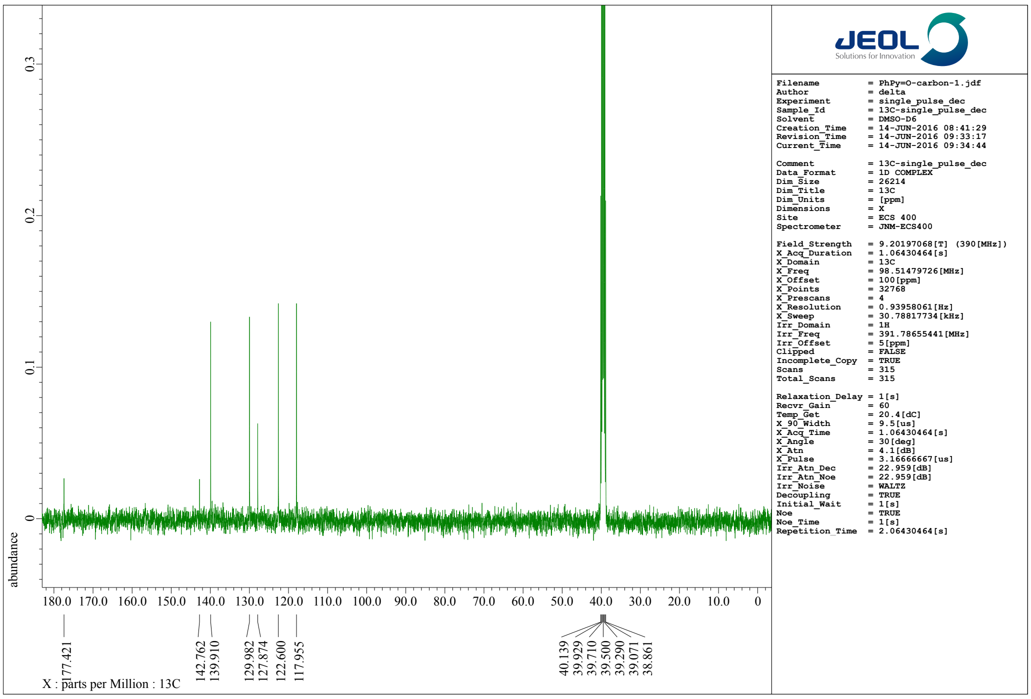


**Fig. S11-a2**. $^{13}$C-NMR (100 MHz) spectrum of **1a** in dimethyl sulfoxide-$d_6$ ($d$-DMSO).

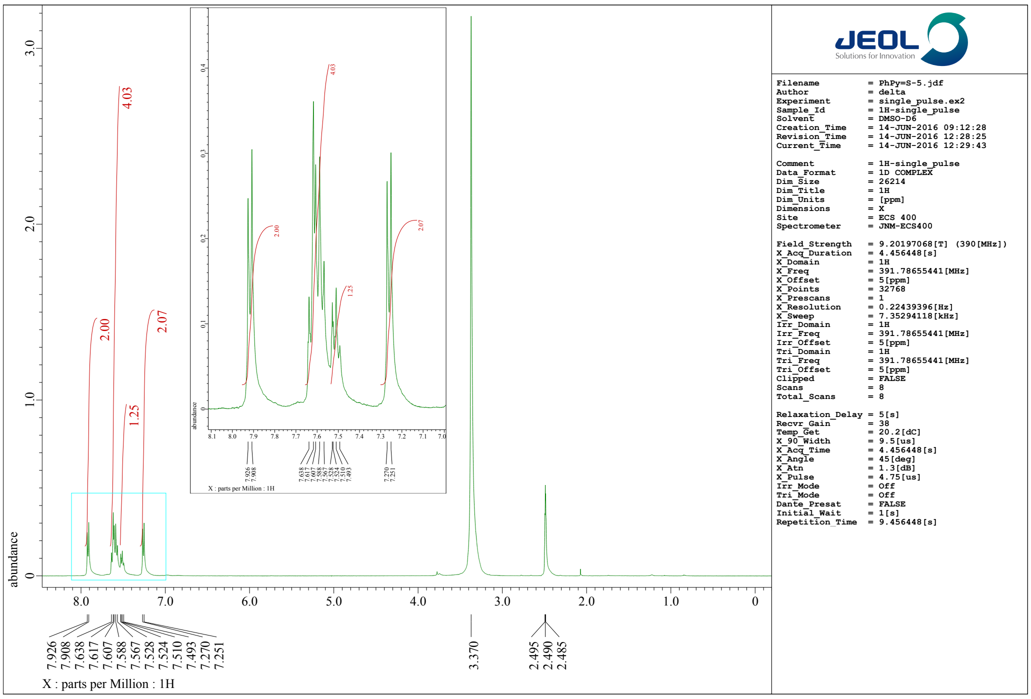


**Fig. S11-a3**. [1]H-NMR (400 MHz) spectrum of **1b** in dimethyl sulfoxide-*d*$_6$ (*d*-DMSO).

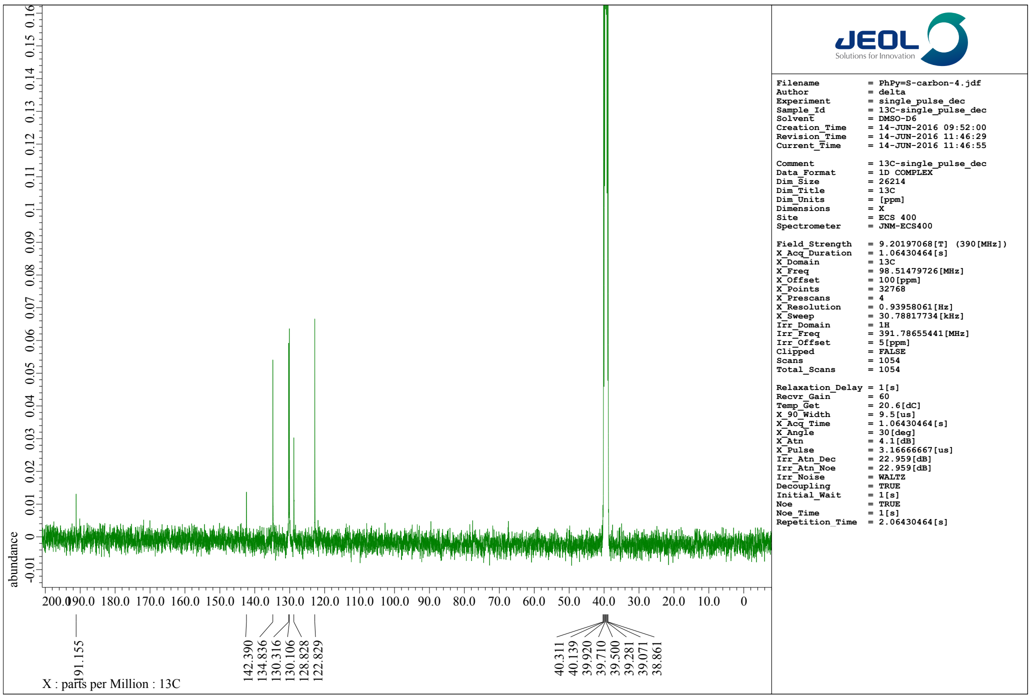


**Fig. S11-a4**. [13]C-NMR (100 MHz) spectrum of **1b** in dimethyl sulfoxide-*d*$_6$ (*d*-DMSO).

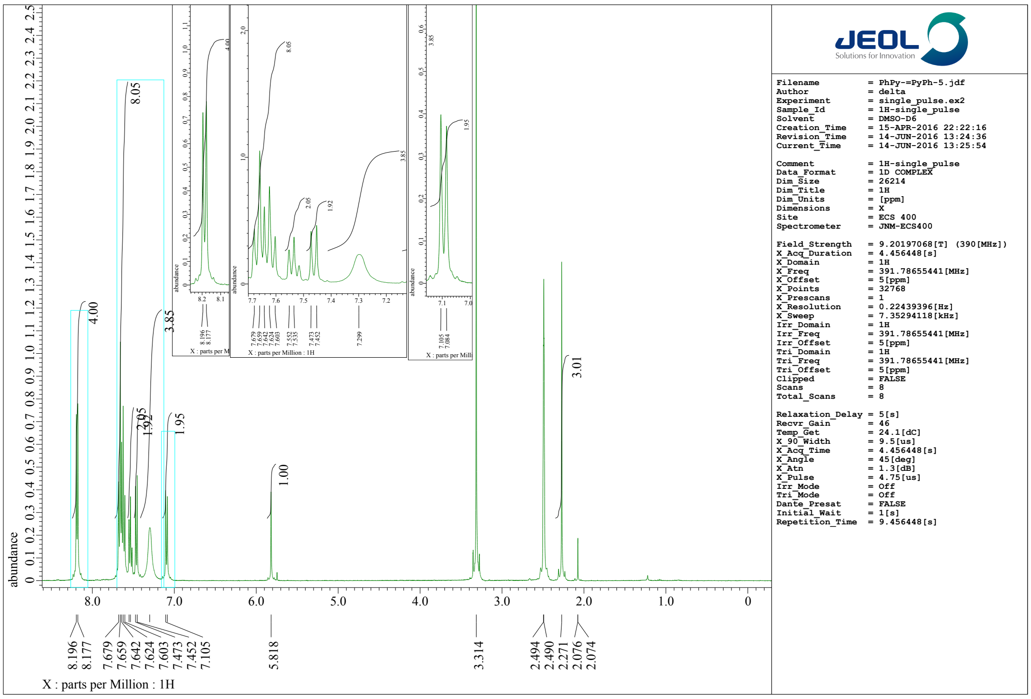


Fig. S11-a5. [1]H-NMR (400 MHz) spectrum of **PC1** in dimethyl sulfoxide-$d_6$ (*d*-DMSO).

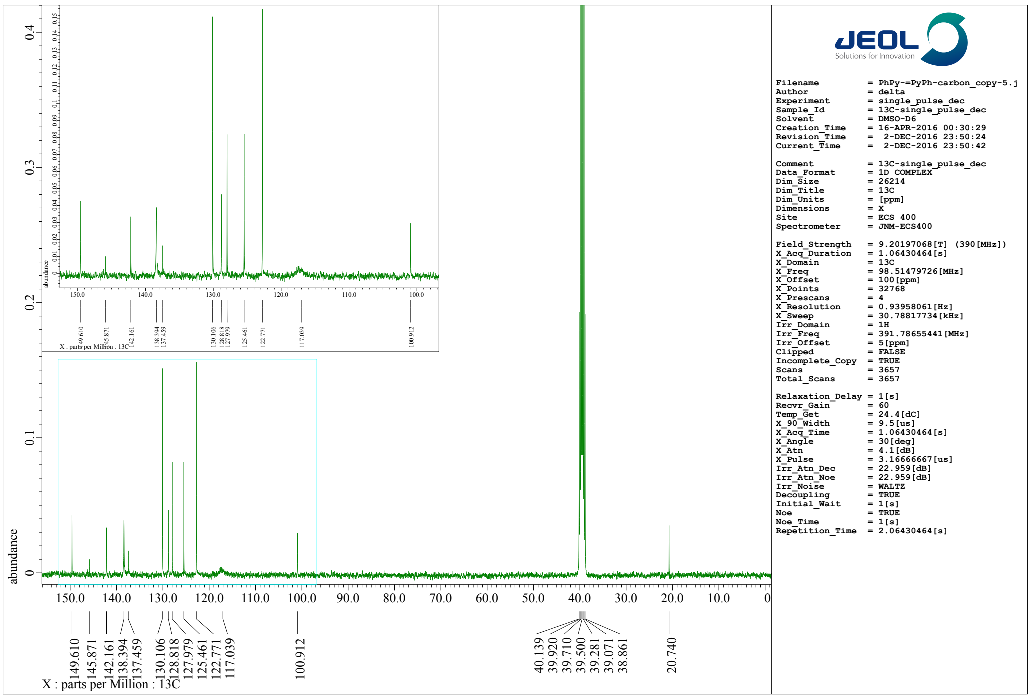


Fig. S11-a6. [13]C-NMR (100 MHz) spectrum of **PC1** in dimethyl sulfoxide-d6 (d-DMSO)

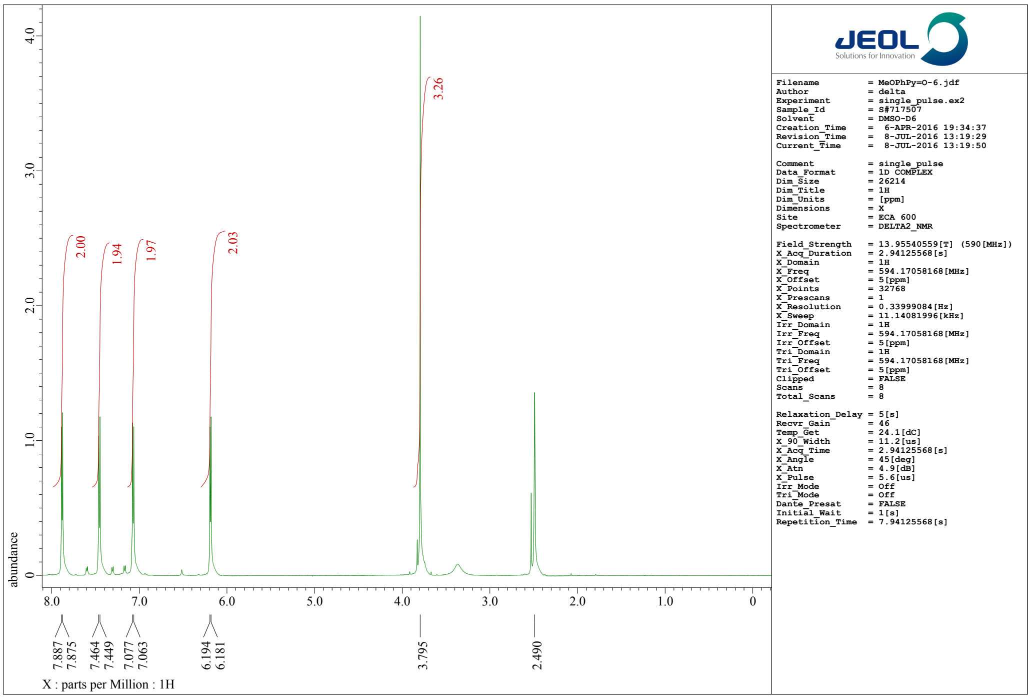


**Fig. S11-b1**. [1]H-NMR (400 MHz) spectrum of **2a** in dimethyl sulfoxide-$d_6$ (*d*-DMSO).

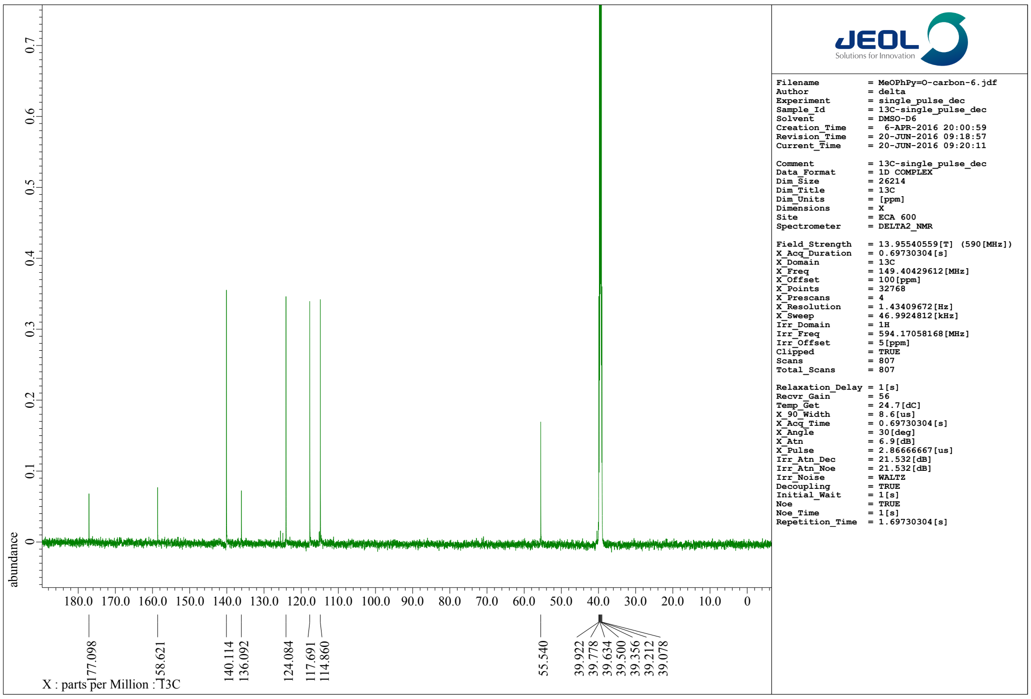


**Fig. S11-b2**. [13]C-NMR (100 MHz) spectrum of **2a** in dimethyl sulfoxide-d6 (d-DMSO)

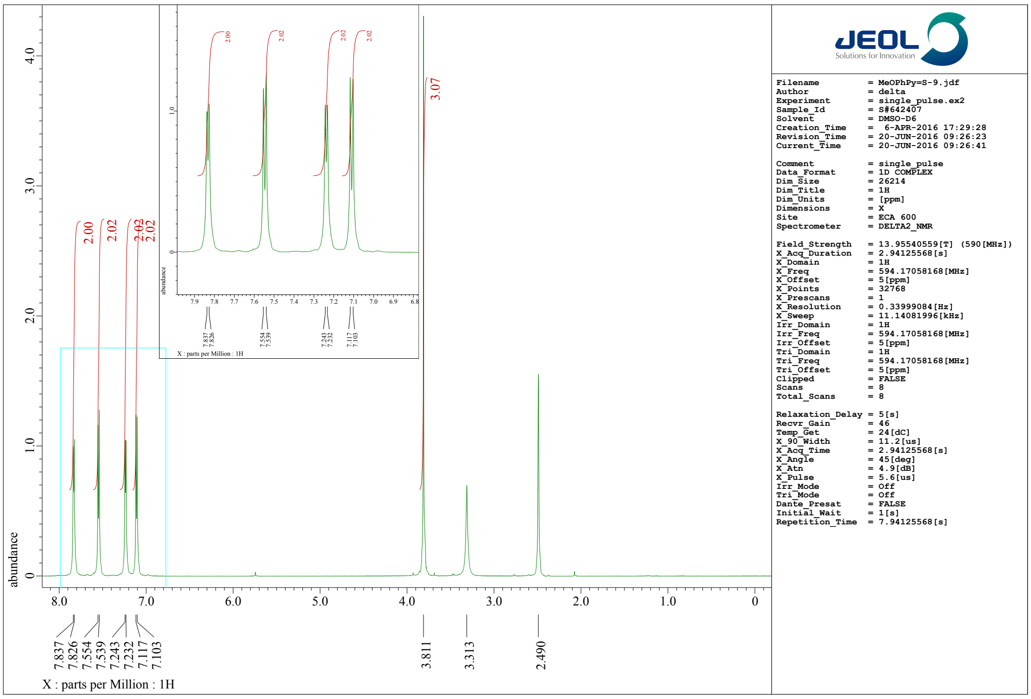


**Fig. S11-b3**. [1]H-NMR (400 MHz) spectrum of **2b** in dimethyl sulfoxide-$d_6$ (*d*-DMSO).

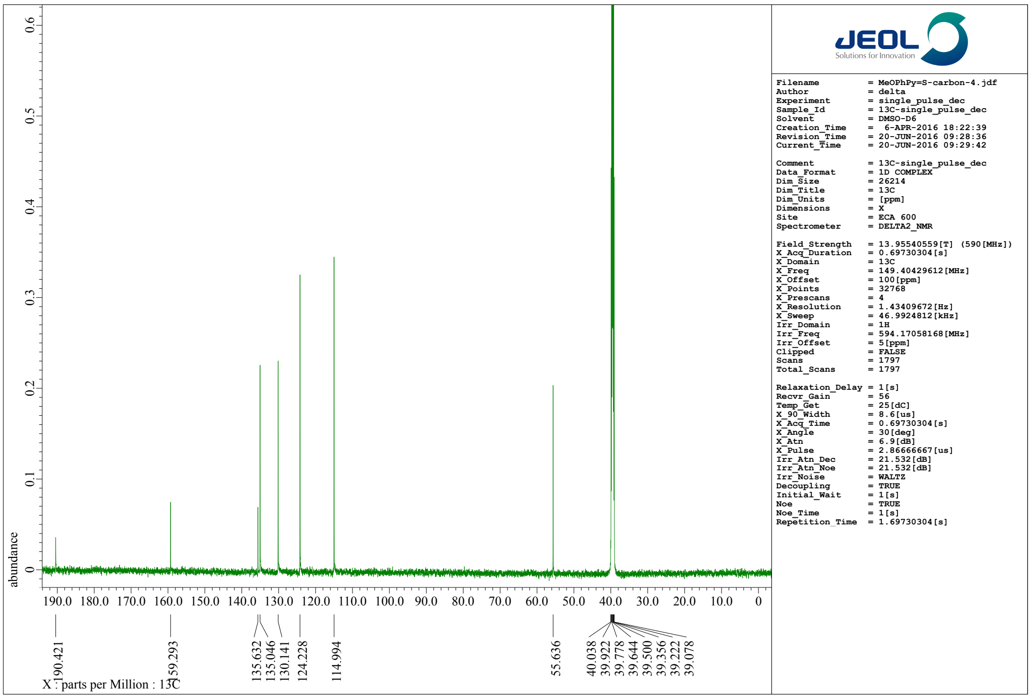


**Fig. S11-b4**. [13]C-NMR (100 MHz) spectrum of **2b** in dimethyl sulfoxide-$d_6$ (*d*-DMSO).

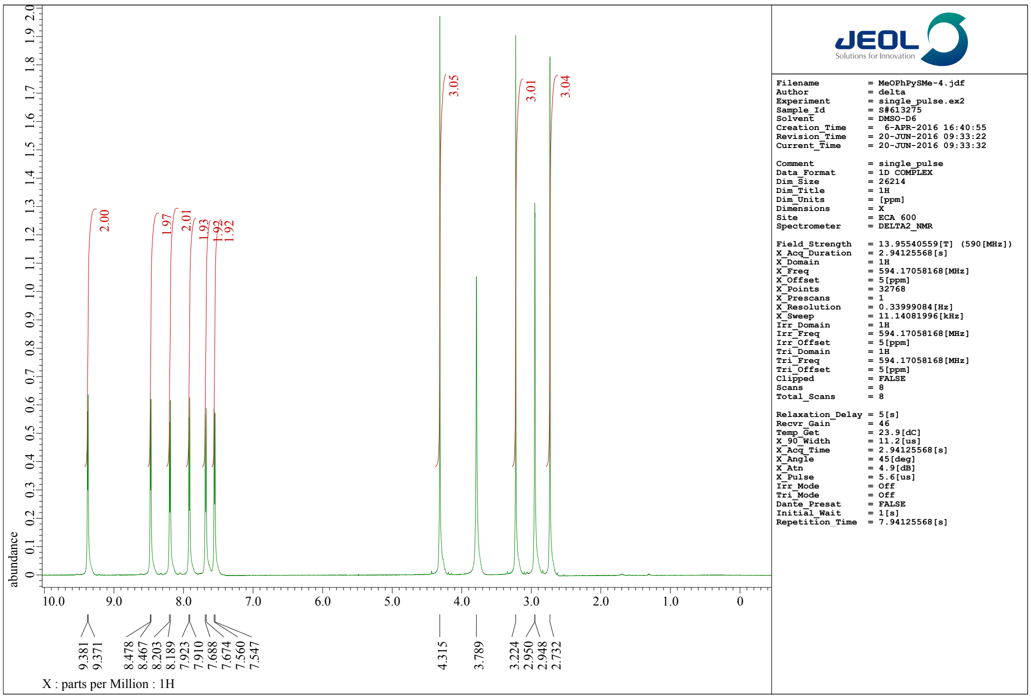


**Fig. S11-b5**. [1]H-NMR (400 MHz) spectrum of **2c** in dimethyl sulfoxide-$d_6$ (*d*-DMSO).

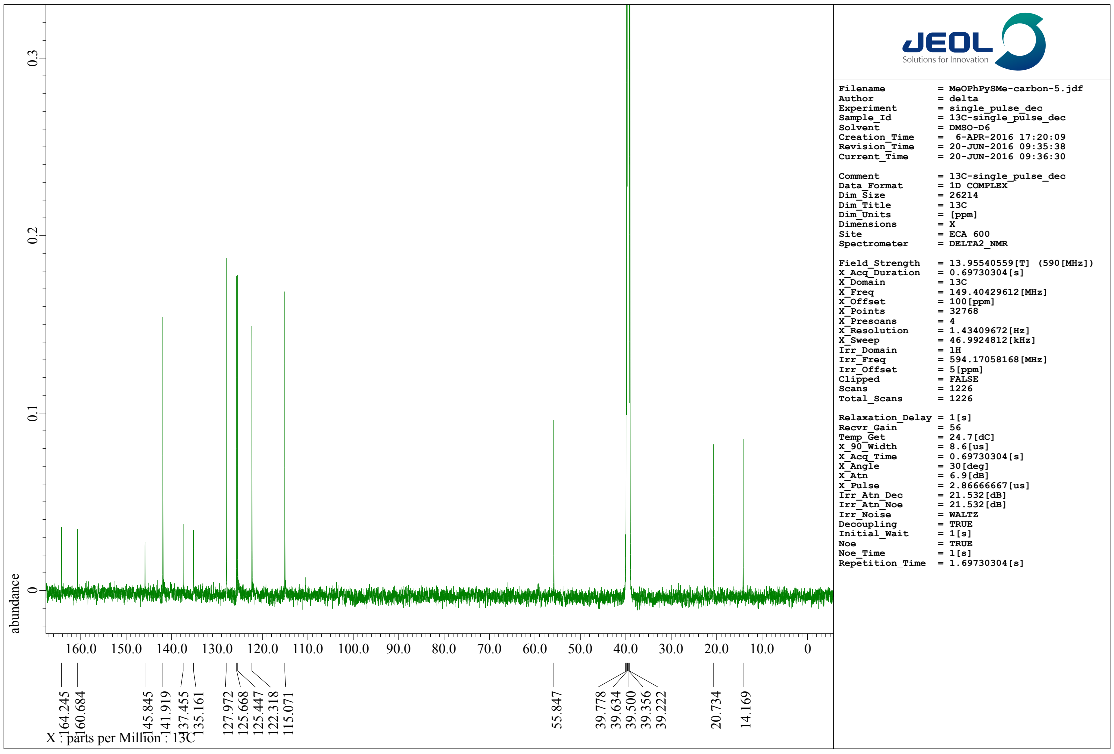


**Fig. S11-b6**. [13]C-NMR (100 MHz) spectrum of **2c** in dimethyl sulfoxide-$d_6$ (*d*-DMSO).

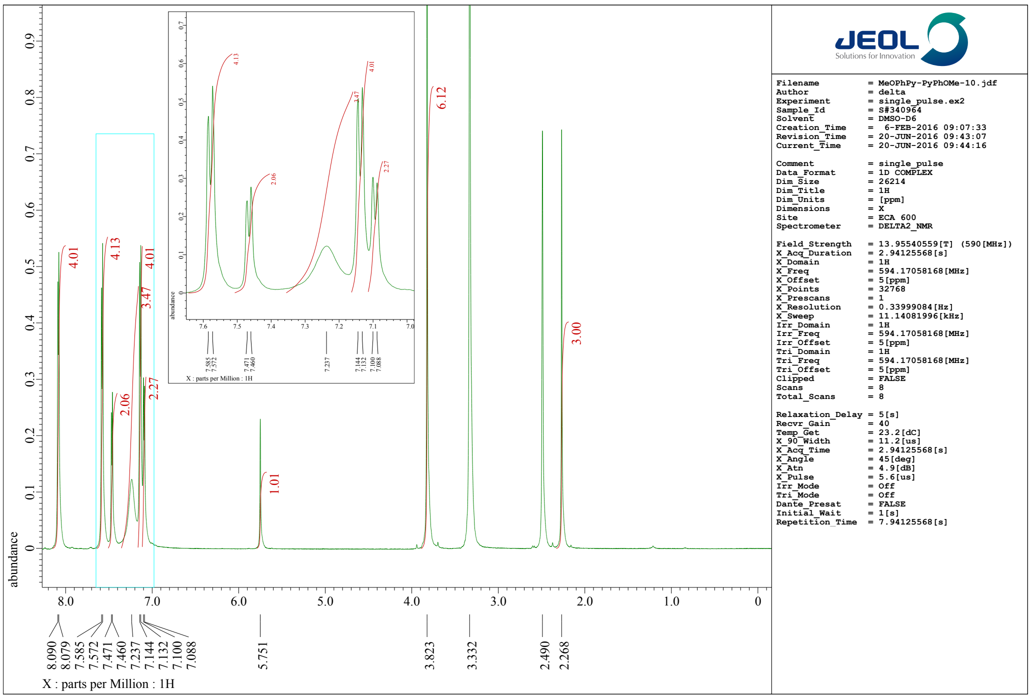


**Fig. S11-b7.** $^{1}$H-NMR (400 MHz) spectrum of **PC2** in dimethyl sulfoxide-$d_6$ ($d$-DMSO).

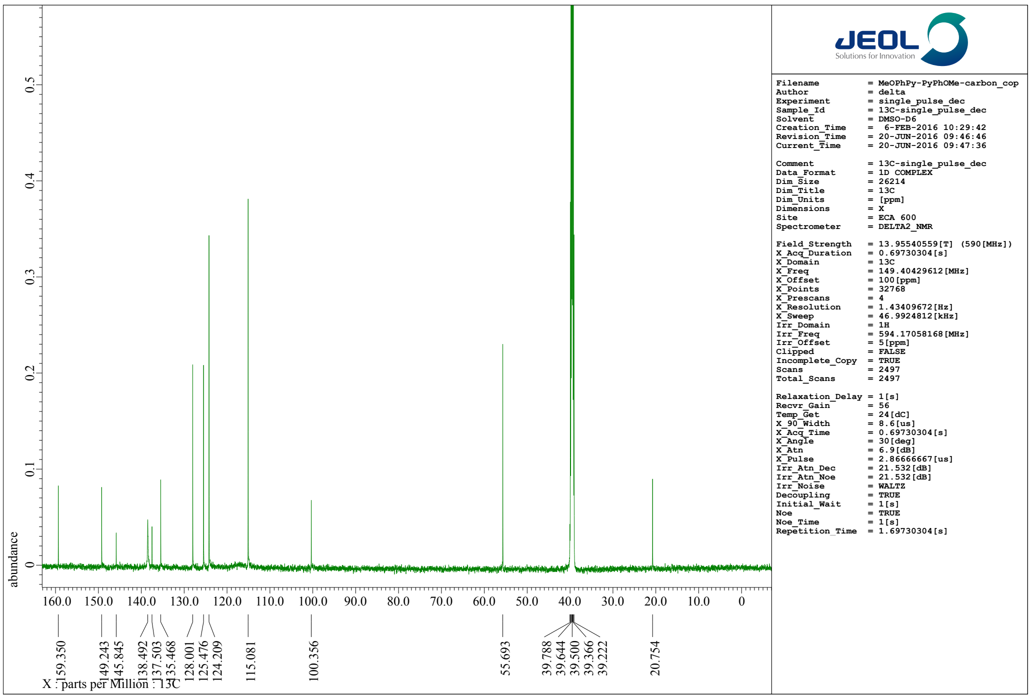


**Fig. S11-b8**. $^{13}$C-NMR (100 MHz) spectrum of **PC2** in dimethyl sulfoxide-$d_6$ ($d$-DMSO).

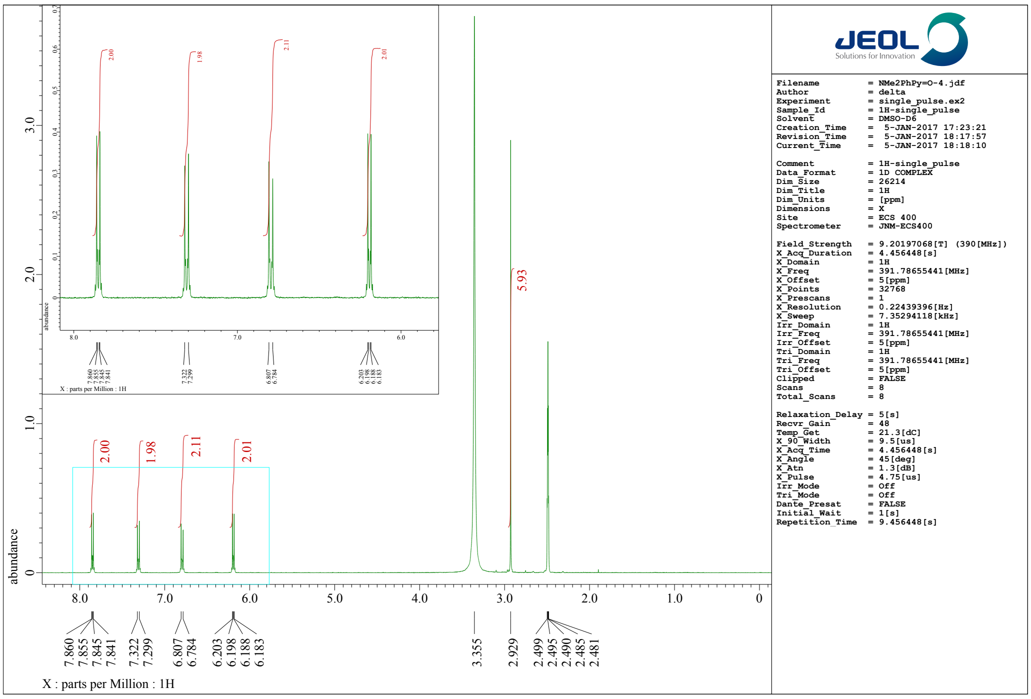


Fig. S11-c1. [1]H-NMR (400 MHz) spectrum of **3a** in dimethyl sulfoxide-$d_6$ (*d*-DMSO).

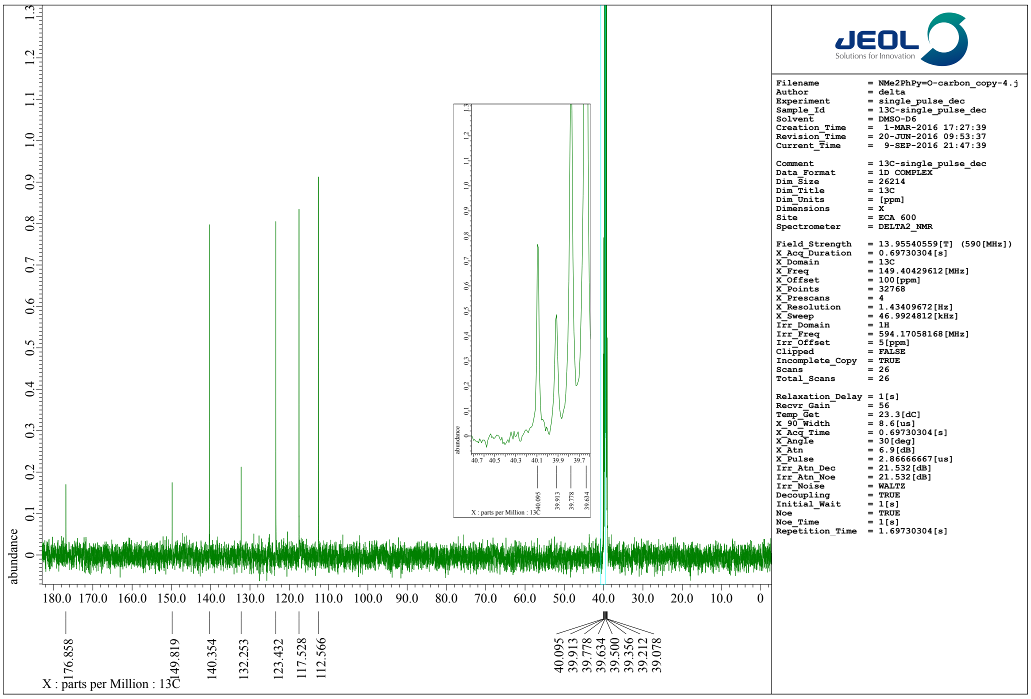


Fig. S11-c2. [13]C-NMR (100 MHz) spectrum of **3a** in dimethyl sulfoxide-$d_6$ (*d*-DMSO).

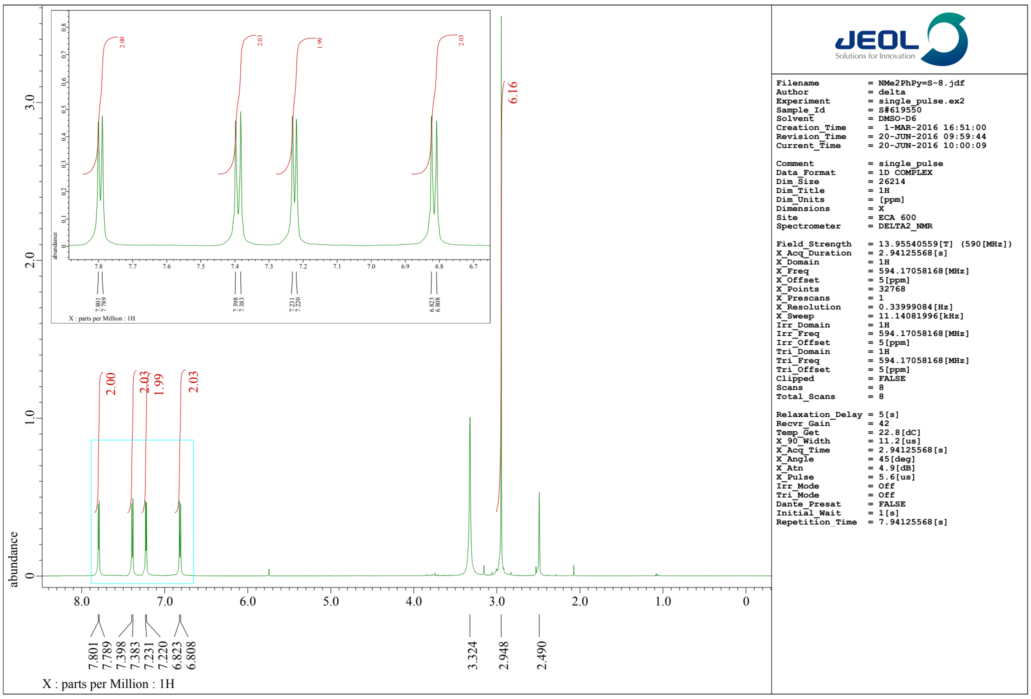


**Fig. S11-c3**. [1]H-NMR (400 MHz) spectrum of **3b** in dimethyl sulfoxide-$d_6$ ($d$-DMSO).

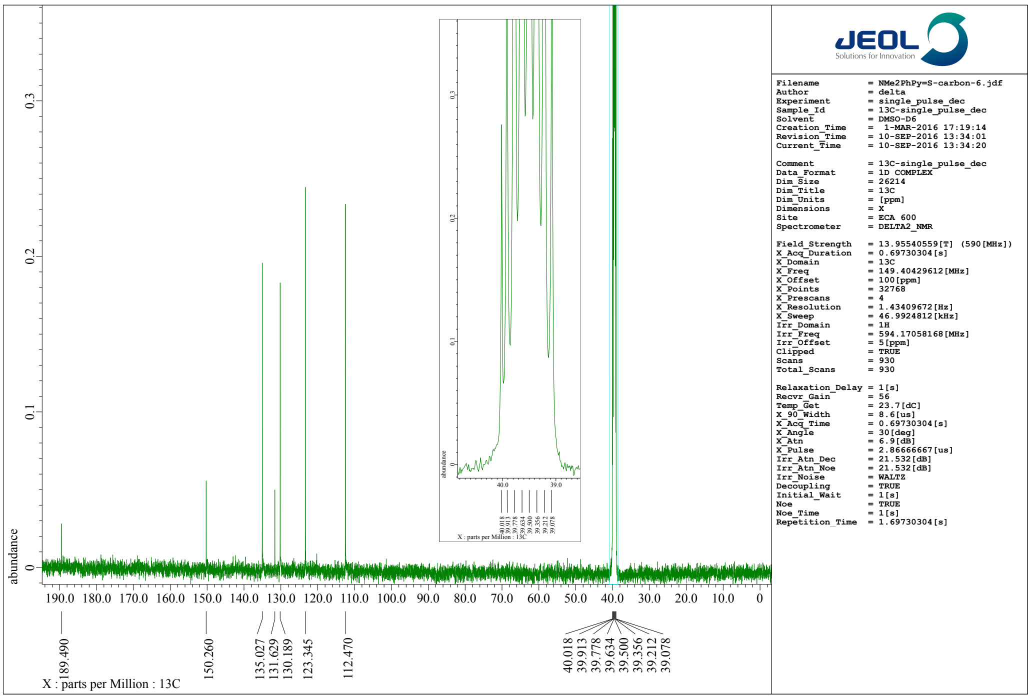


**Fig. S11-c4**. [13]C-NMR (100 MHz) spectrum of **3b** in dimethyl sulfoxide-$d_6$ ($d$-DMSO).

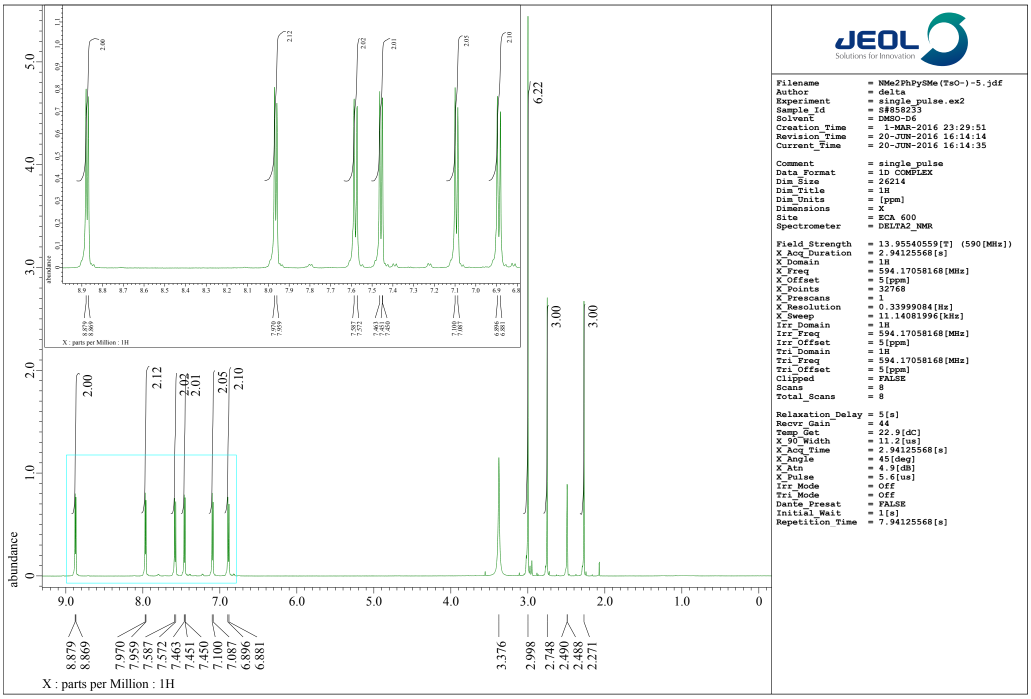


**Fig. S11-c5**. [1]H-NMR (400 MHz) spectrum of **3c** in dimethyl sulfoxide-$d_6$ ($d$-DMSO).

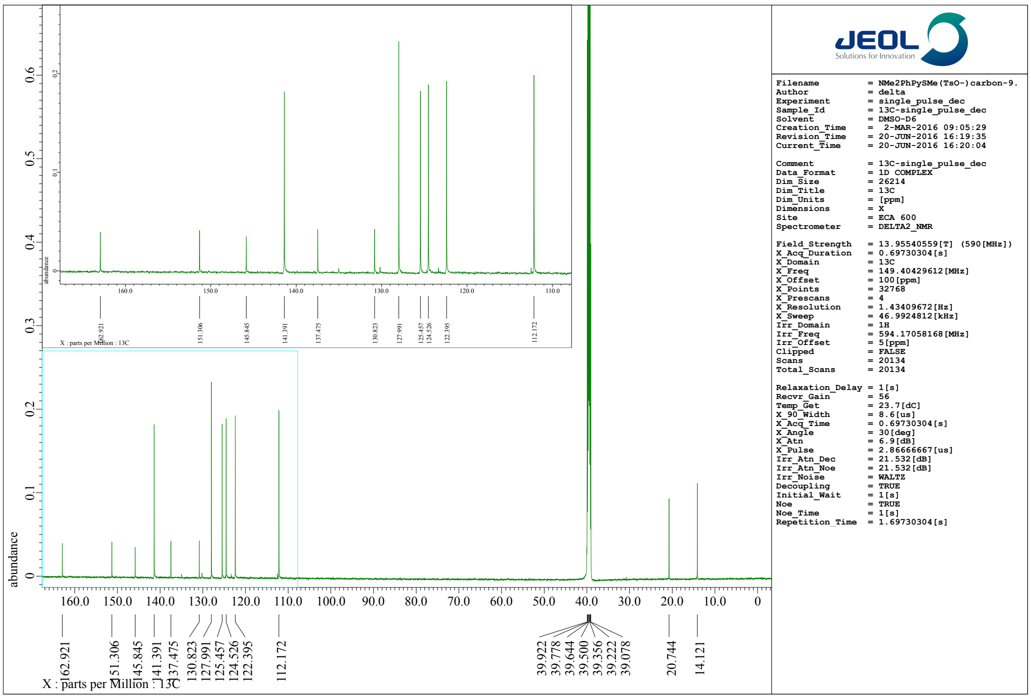


**Fig. S11-c6**. [13]C-NMR (100 MHz) spectrum of **3c** in dimethyl sulfoxide-$d_6$ ($d$-DMSO).

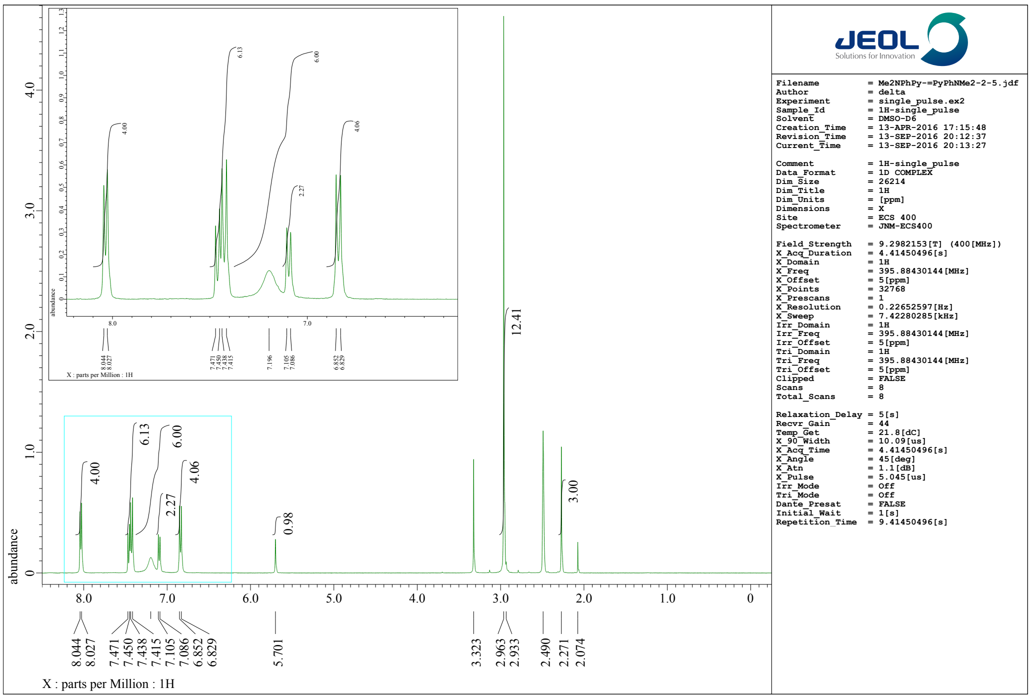


**Fig. S11-c7**. $^{1}$H-NMR (400 MHz) spectrum of **PC3** in dimethyl sulfoxide-$d_6$ ($d$-DMSO).

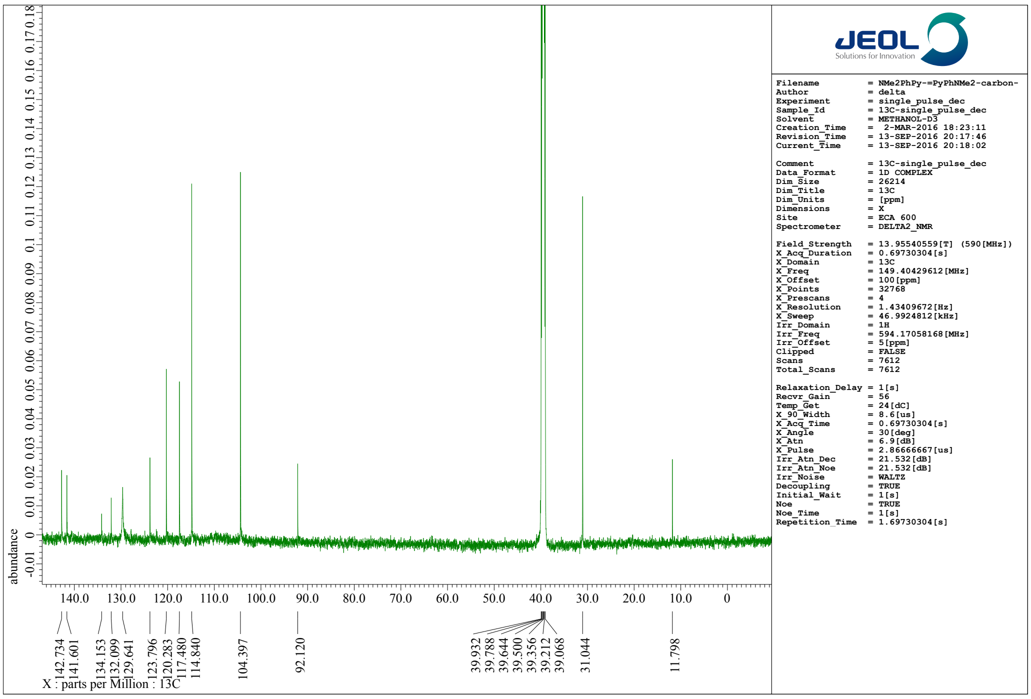


**Fig. S11-c8**. $^{13}$C-NMR (100 MHz) spectrum of **PC3** in dimethyl sulfoxide-$d_6$ ($d$-DMSO).

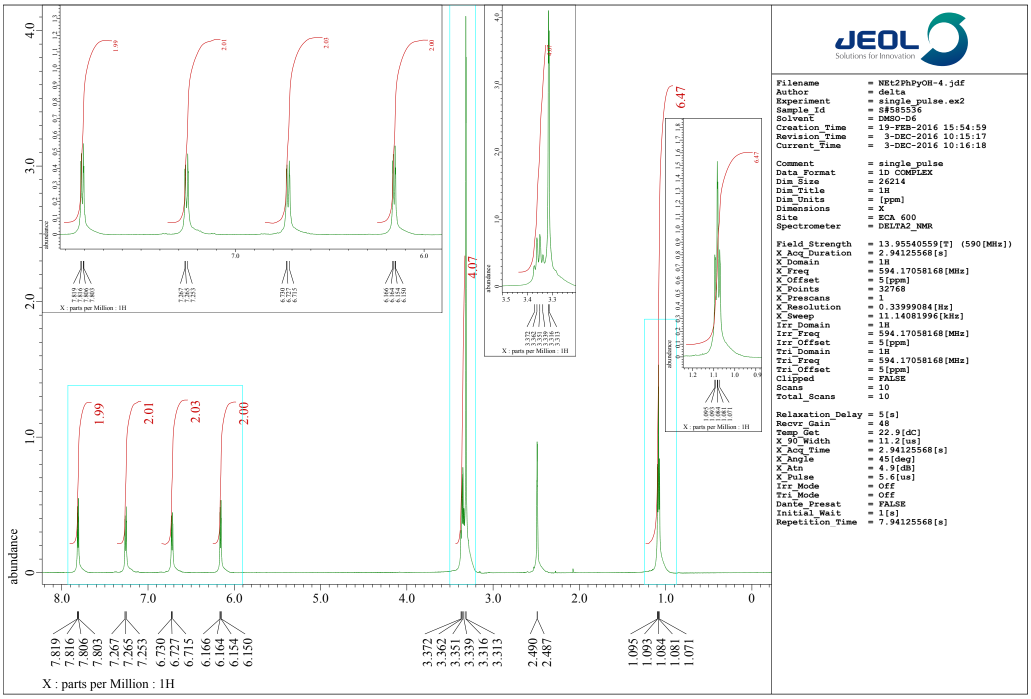


**Fig. S11-d1**. $^{1}$H-NMR (400 MHz) spectrum of **4a** in dimethyl sulfoxide-$d_6$ ($d$-DMSO).

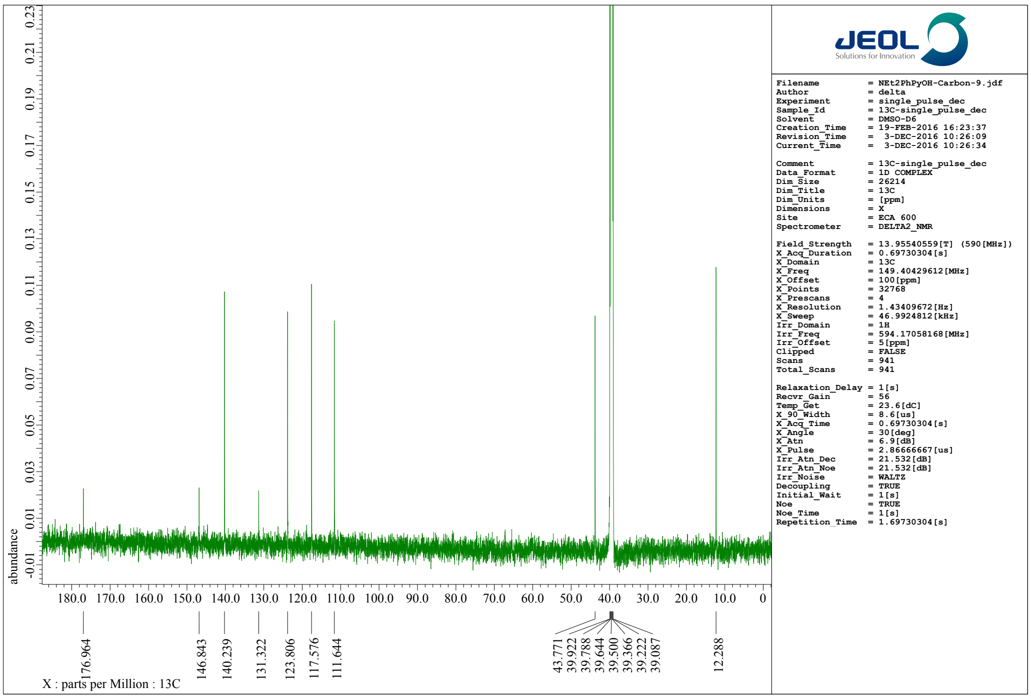


**Fig. S11-d2**. $^{13}$C-NMR (100 MHz) spectrum of **4a** in dimethyl sulfoxide-$d_6$ ($d$-DMSO).

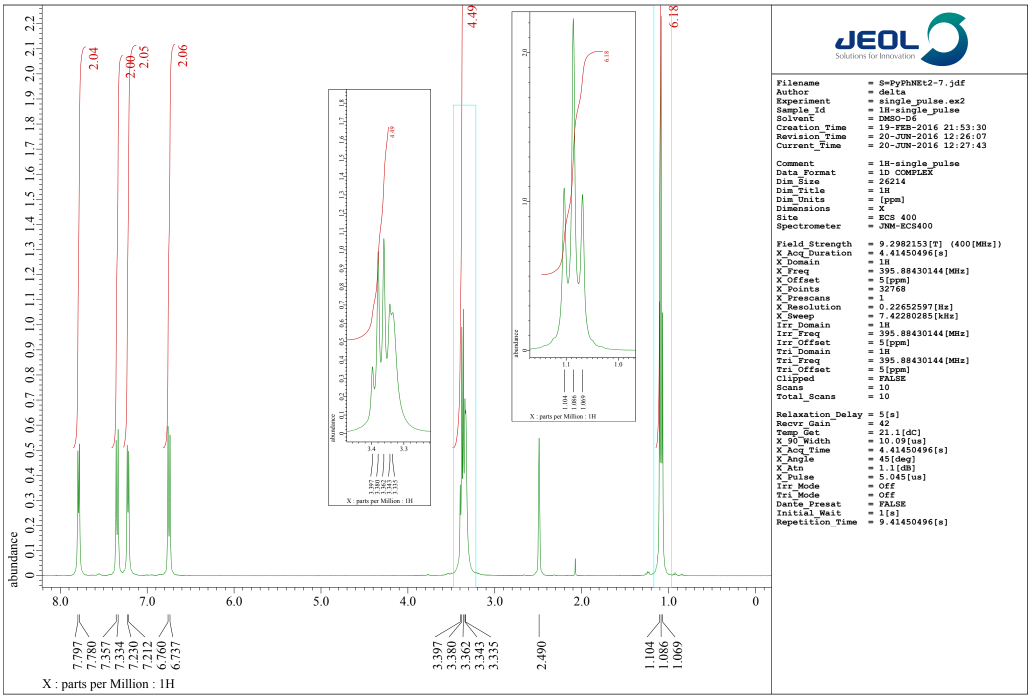


Fig. S11-d3. [1]H-NMR (400 MHz) spectrum of **4b** in dimethyl sulfoxide-$d_6$ (*d*-DMSO).

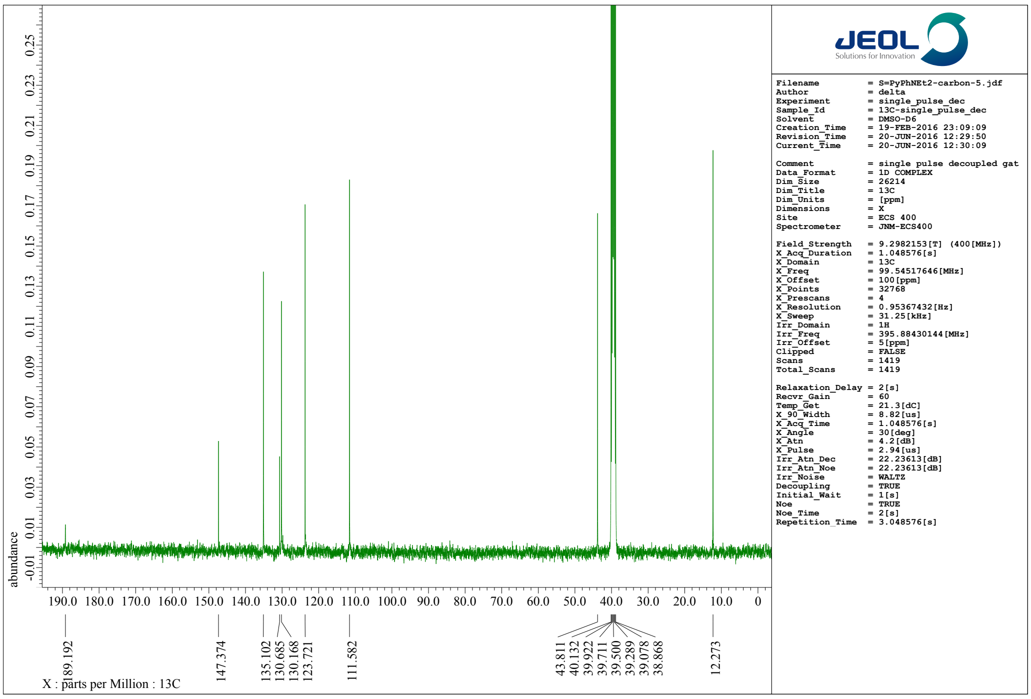


Fig. S11-d4. [13]C-NMR (100 MHz) spectrum of **4b** in dimethyl sulfoxide-$d_6$ (*d*-DMSO).

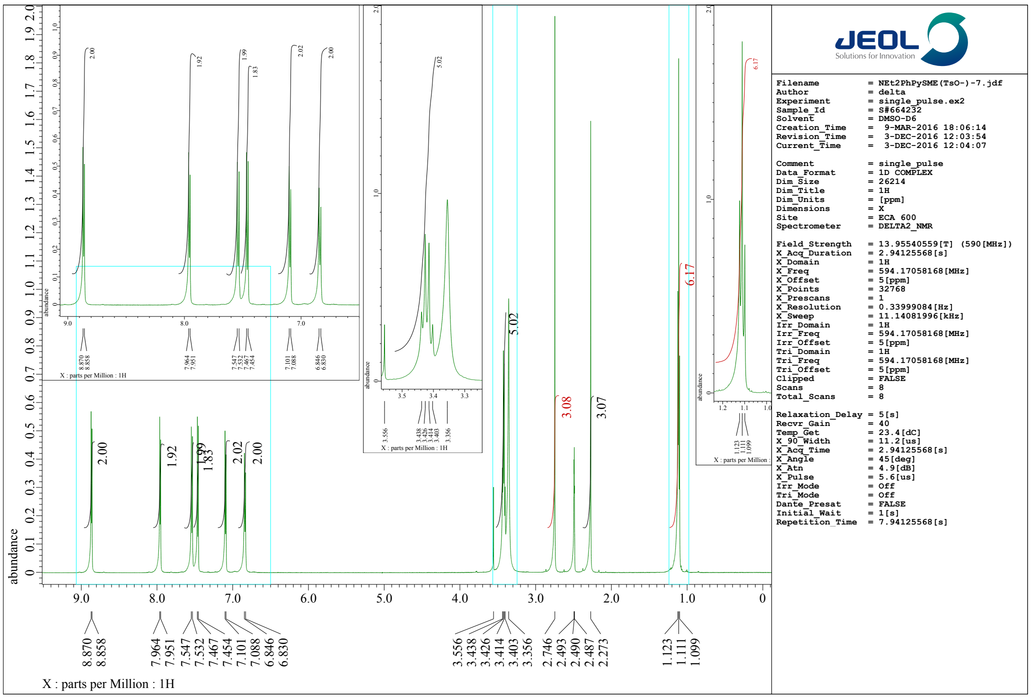


**Fig. S11-d5**. [1]H-NMR (400 MHz) spectrum of **4c** in dimethyl sulfoxide-$d_6$ ($d$-DMSO).

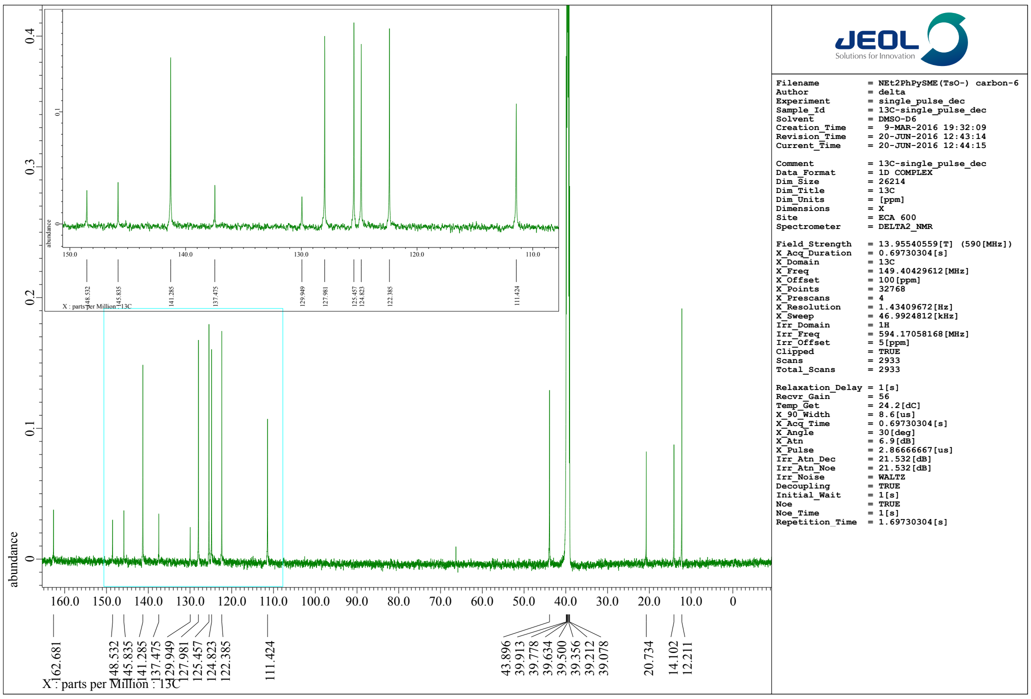


**Fig. S11-d6**. [13]C-NMR (100 MHz) spectrum of **4c** in dimethyl sulfoxide-$d_6$ ($d$-DMSO).

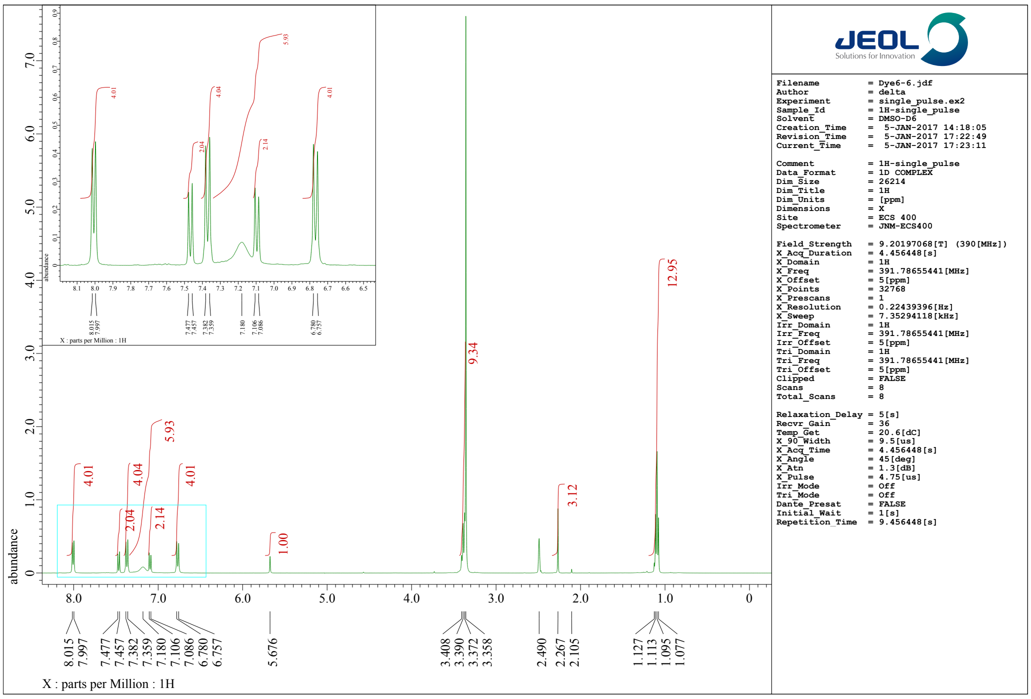


**Fig. S11-d7**. [1]H-NMR (400 MHz) spectrum of **PC4** in dimethyl sulfoxide-$d_6$ (*d*-DMSO).

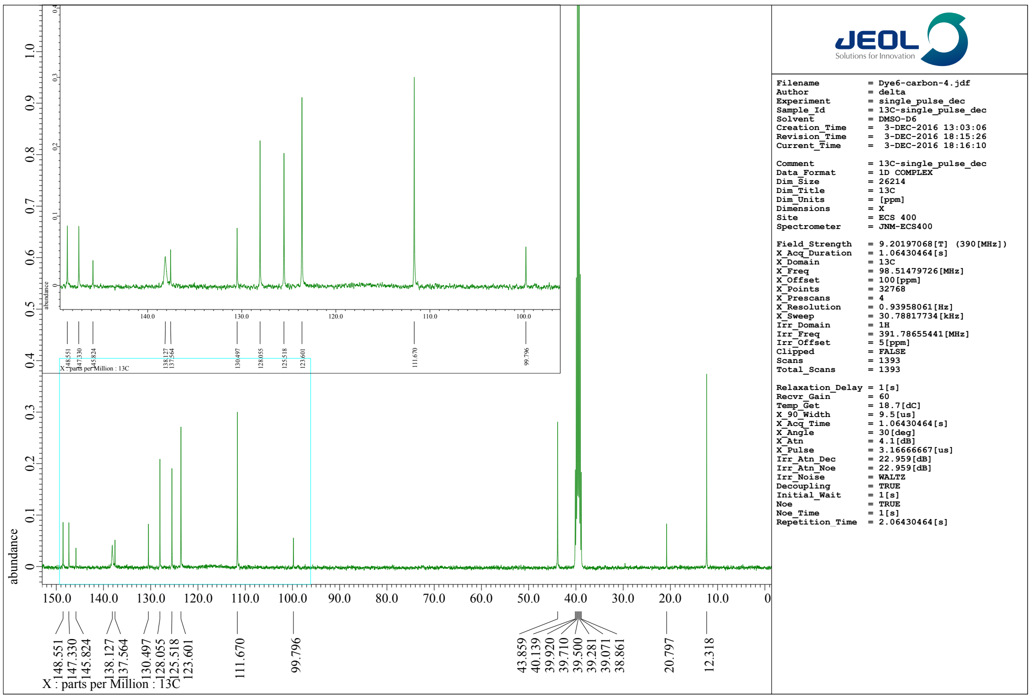


**Fig. S11-d8**. [13]C-NMR (100 MHz) spectrum of **PC4** in dimethyl sulfoxide-$d_6$ (*d*-DMSO).

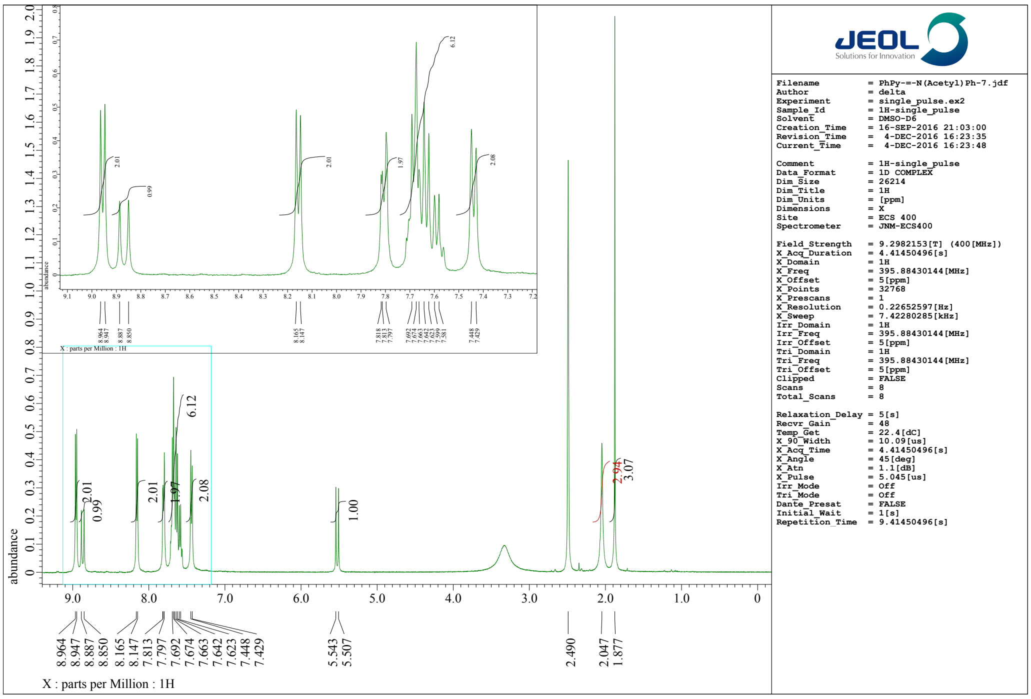


**Fig. S11-e1**. [1]H-NMR (400 MHz) spectrum of **5c** in dimethyl sulfoxide-$d_6$ ($d$-DMSO).

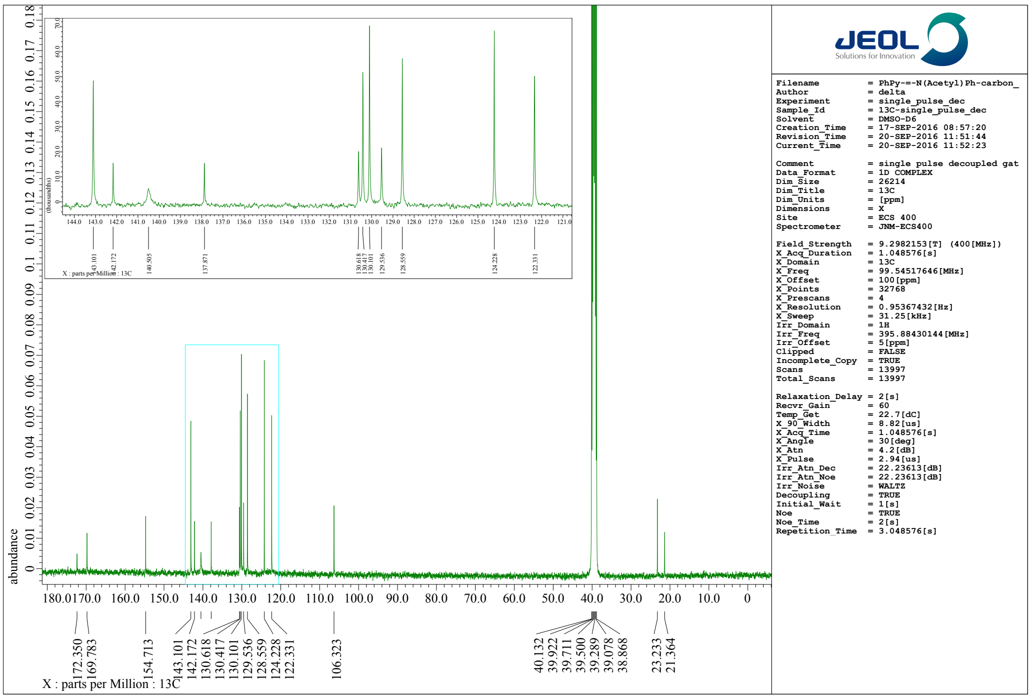


**Fig. S11-e2**. [13]C-NMR (100 MHz) spectrum of **5c** in dimethyl sulfoxide-$d_6$ ($d$-DMSO).

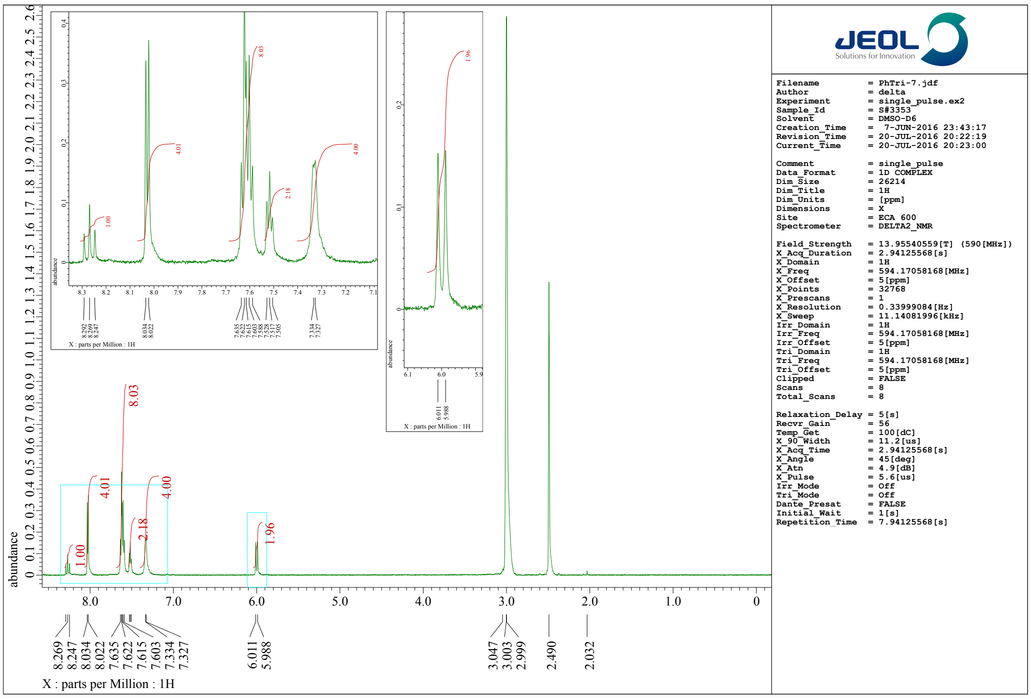


Fig. S11-e3. [1]H-NMR (400 MHz) spectrum of **PC5** in dimethyl sulfoxide-$d_6$ (*d*-DMSO).

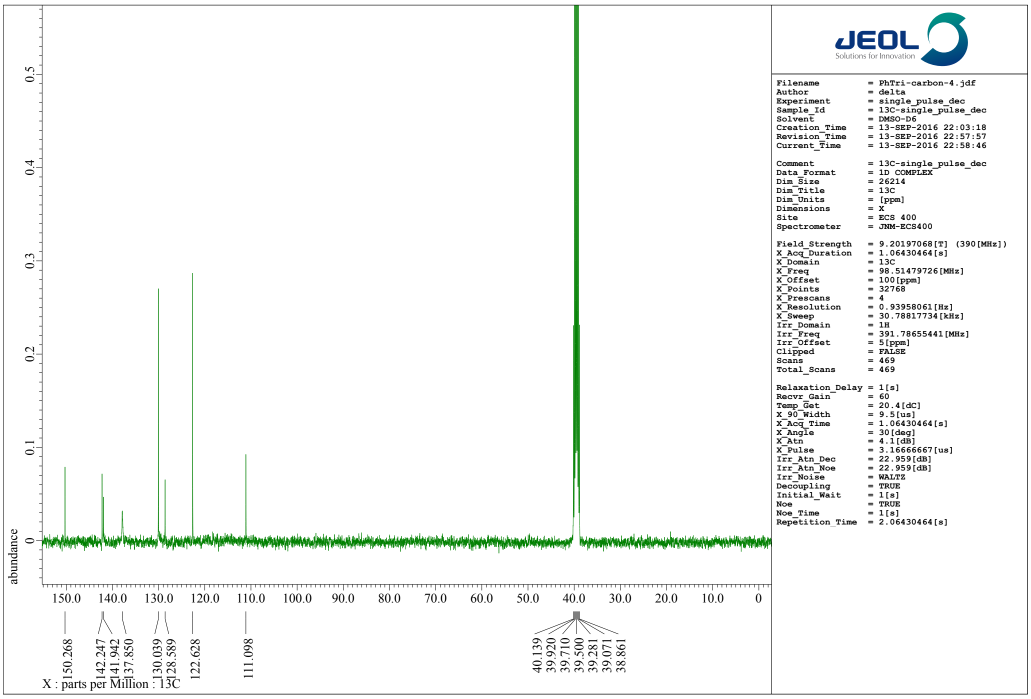


Fig. S11-e4. [13]C-NMR (100 MHz) spectrum of **PC5** in dimethyl sulfoxide-$d_6$ (*d*-DMSO).

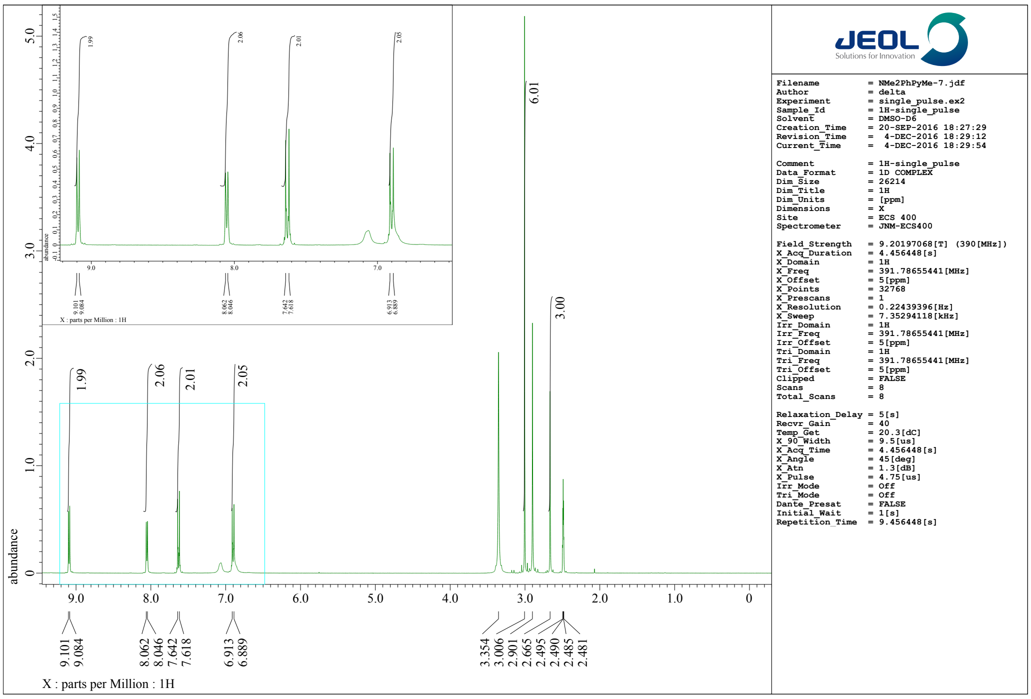


**Fig. S11-f1**. $^{1}$H-NMR (400 MHz) spectrum of **6a** in dimethyl sulfoxide-*d*$_6$ (*d*-DMSO).

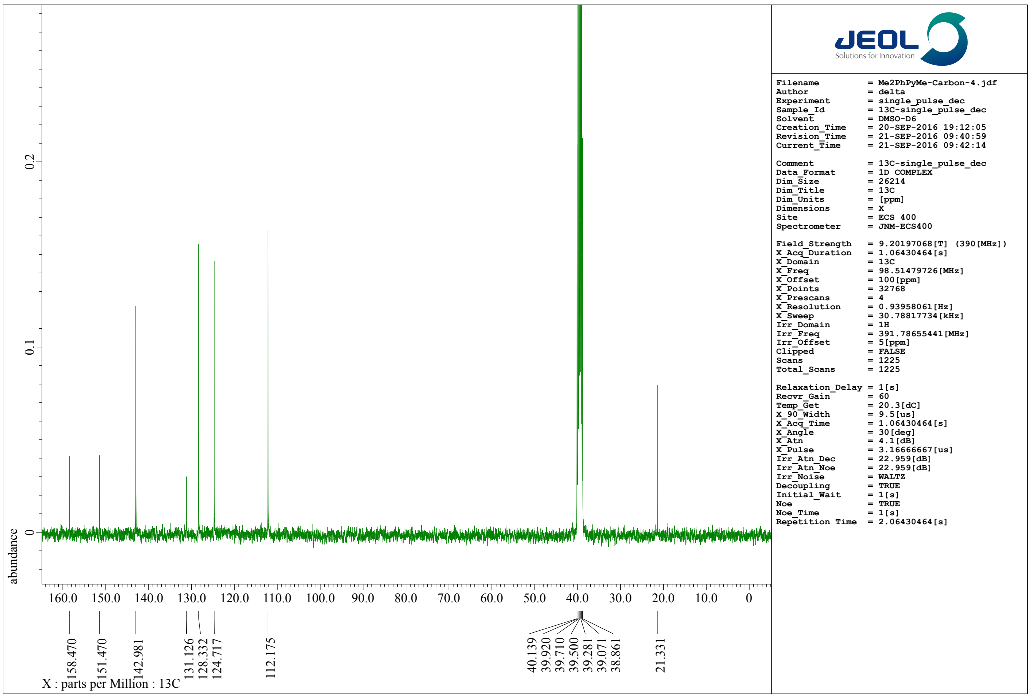


**Fig. S11-f2**. $^{13}$C-NMR (100 MHz) spectrum of **6a** in dimethyl sulfoxide-d6 (d-DMSO)

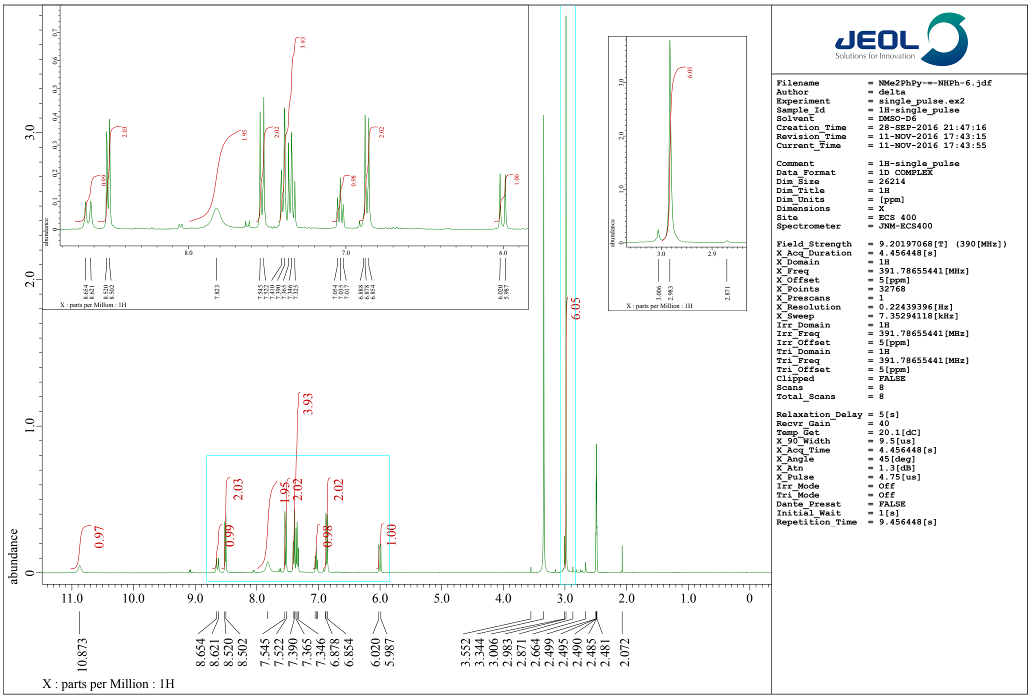


**Fig. S11-f3**. [1]H-NMR (400 MHz) spectrum of **6b** in dimethyl sulfoxide-$d_6$ ($d$-DMSO).

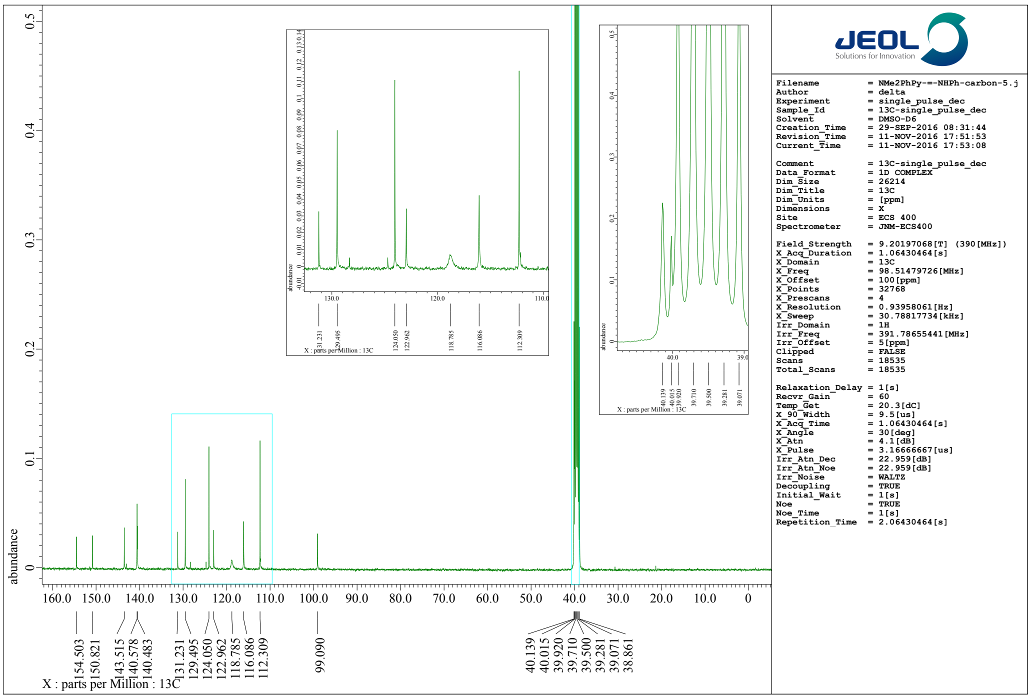


**Fig. S11-f4**. [13]C-NMR (100 MHz) spectrum of **6b** in dimethyl sulfoxide-$d_6$ ($d$-DMSO).

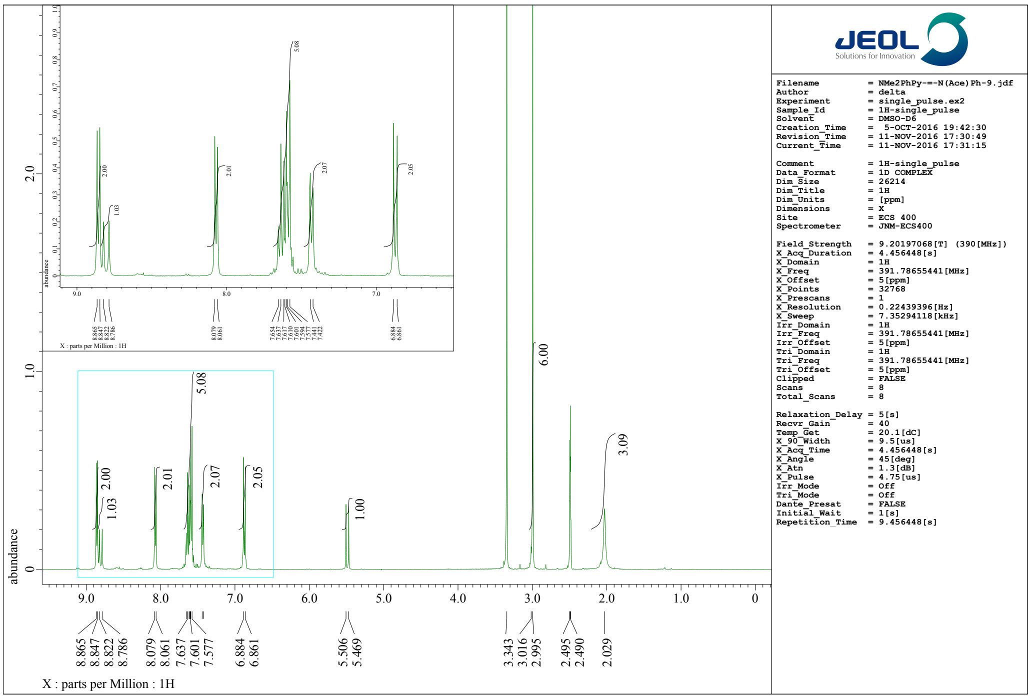


**Fig. S11-f5**. [1]H-NMR (400 MHz) spectrum of **6c** in dimethyl sulfoxide-$d_6$ ($d$-DMSO).

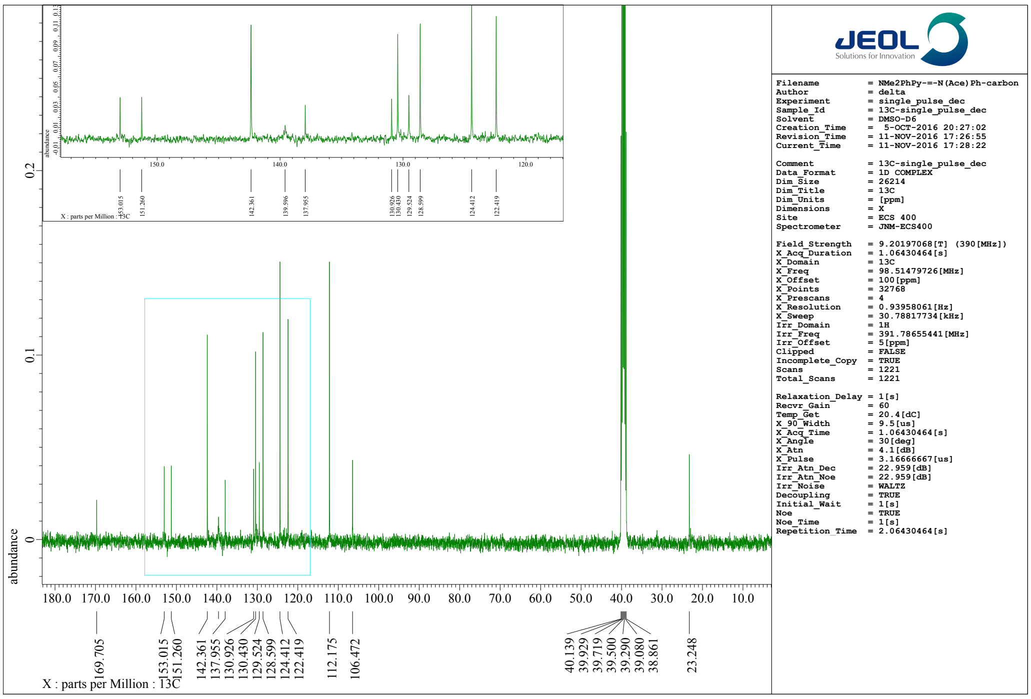


**Fig. S11-f6**. [13]C-NMR (100 MHz) spectrum of **6c** in dimethyl sulfoxide-$d_6$ ($d$-DMSO).

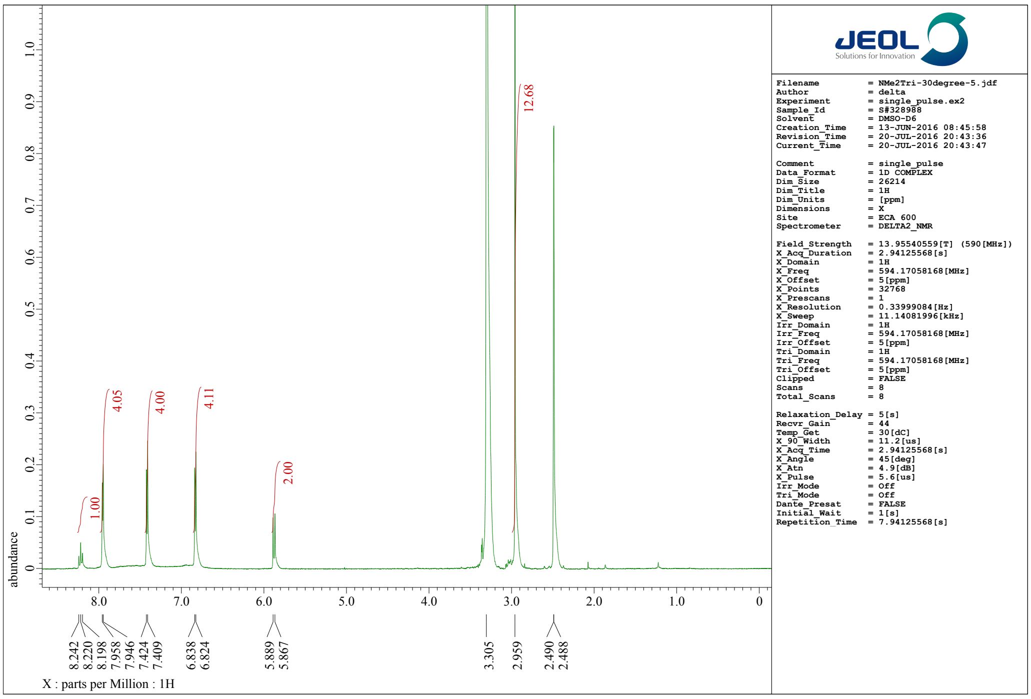


**Fig. S11-f7**. [1]H-NMR (400 MHz) spectrum of **PC6** in dimethyl sulfoxide-$d_6$ ($d$-DMSO).

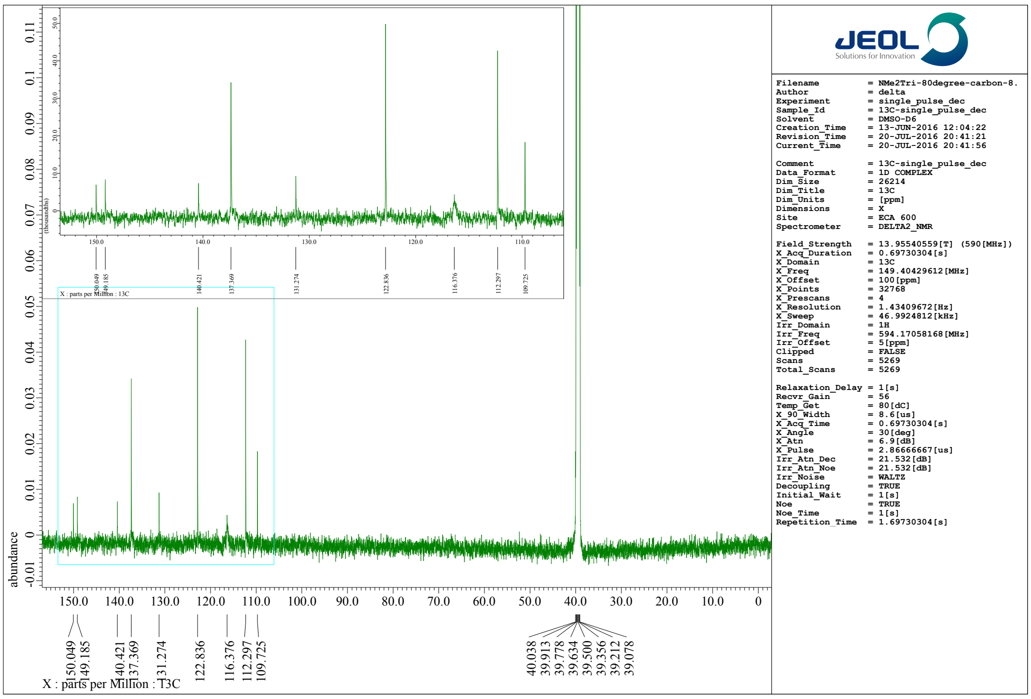


**Fig. S11-f8**. [13]C-NMR (100 MHz) spectrum of **PC6** in dimethyl sulfoxide-$d_6$ ($d$-DMSO).

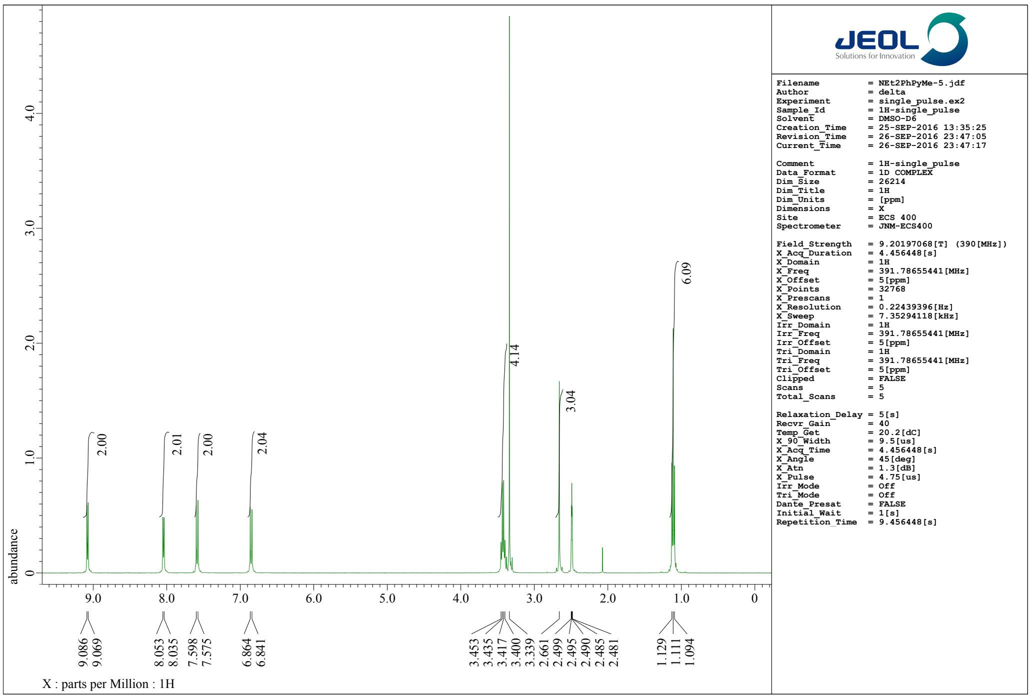


**Fig. S11-g1**. [1]H-NMR (400 MHz) spectrum of **7a** in dimethyl sulfoxide-$d_6$ ($d$-DMSO).

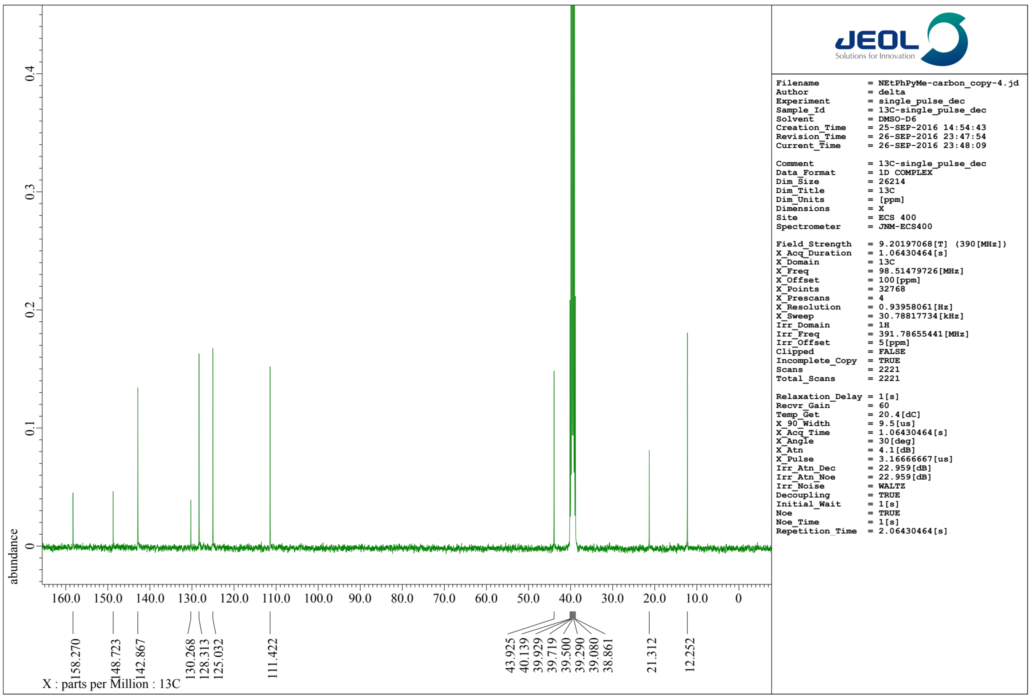


**Fig. S11-g2**. [13]C-NMR (100 MHz) spectrum of **7a** in dimethyl sulfoxide-$d_6$ ($d$-DMSO).

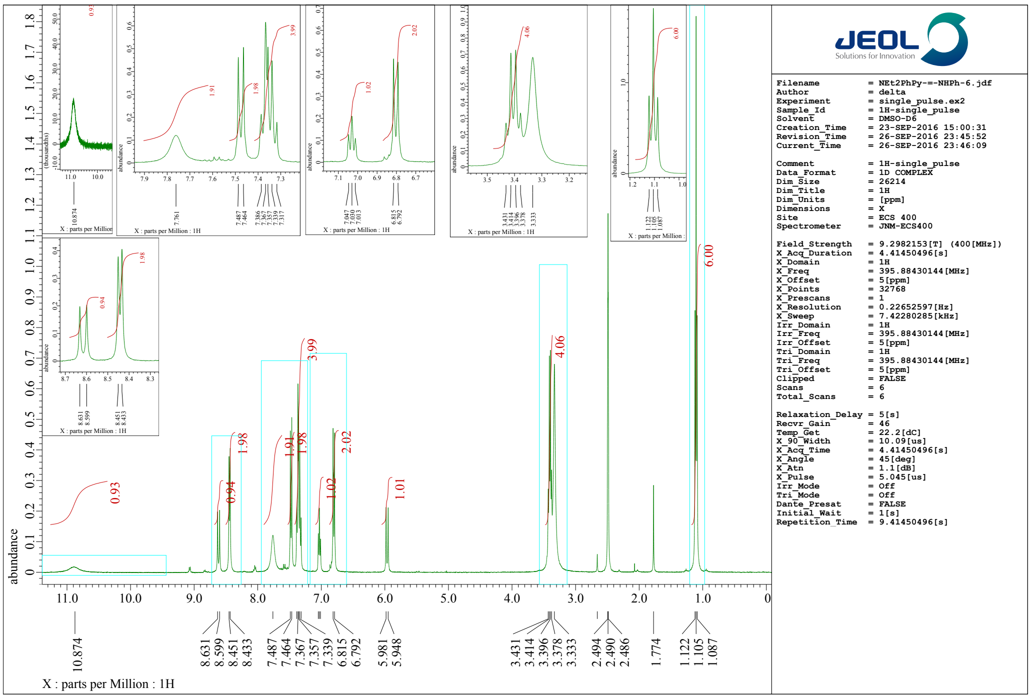


Fig. S11-g3. [1]H-NMR (400 MHz) spectrum of **7b** in dimethyl sulfoxide-d6 (d-DMSO)

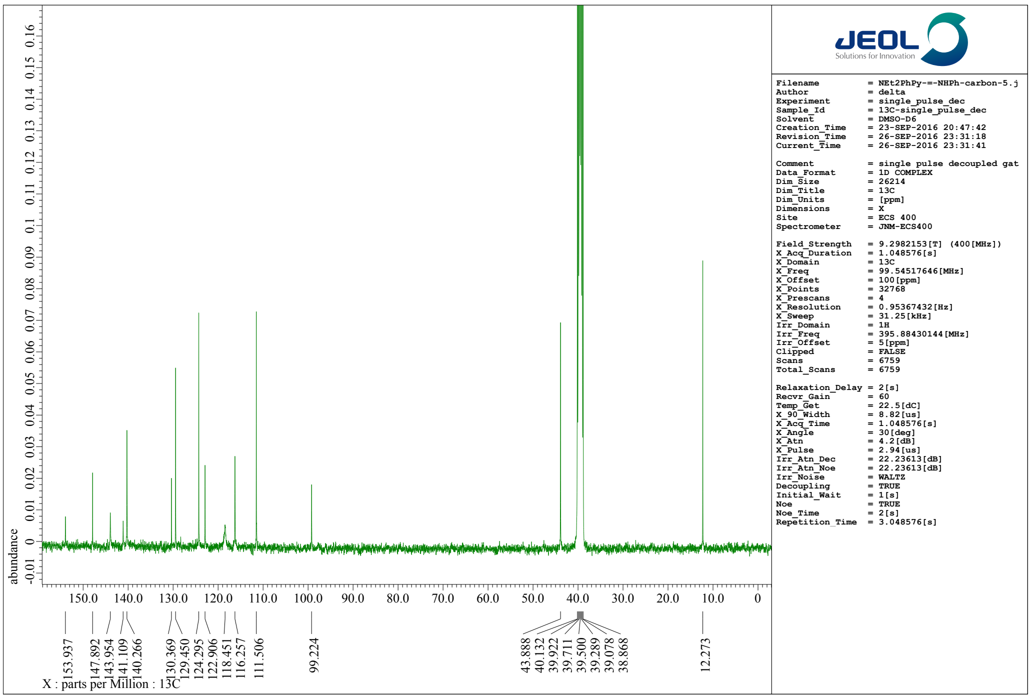


Fig. S11-g4. [13]C-NMR (100 MHz) spectrum of **7b** in dimethyl sulfoxide-$d_6$ ($d$-DMSO).

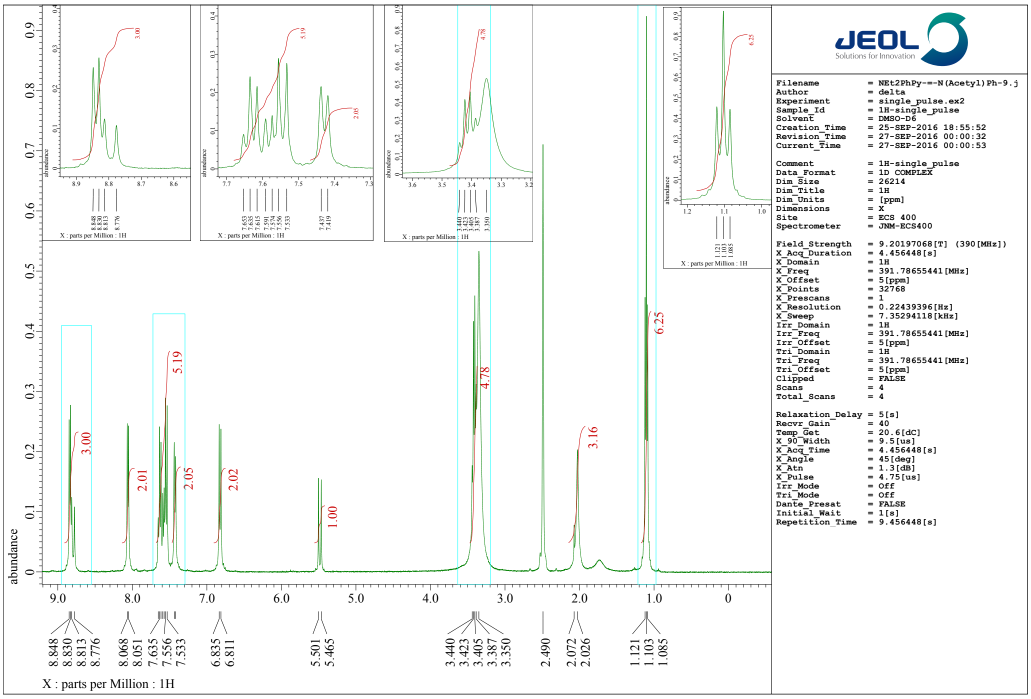


**Fig. S11-g5**. [1]H-NMR (400 MHz) spectrum of **7c** in dimethyl sulfoxide-$d_6$ ($d$-DMSO).

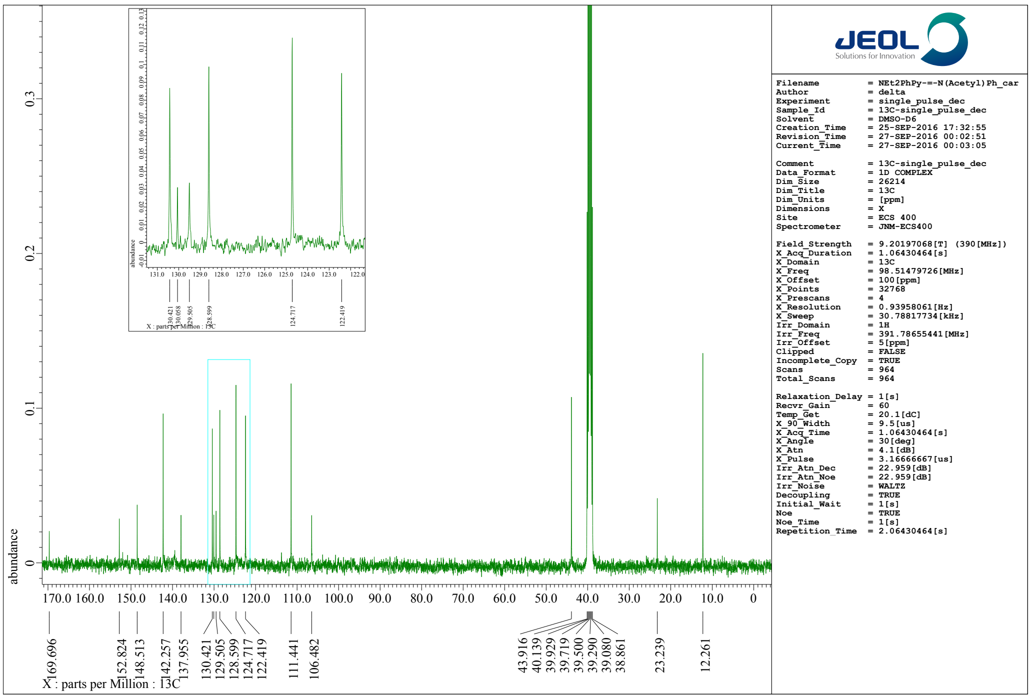


**Fig. S11-g6**. [13]C-NMR (100 MHz) spectrum of **7c** in dimethyl sulfoxide-$d_6$ ($d$-DMSO).

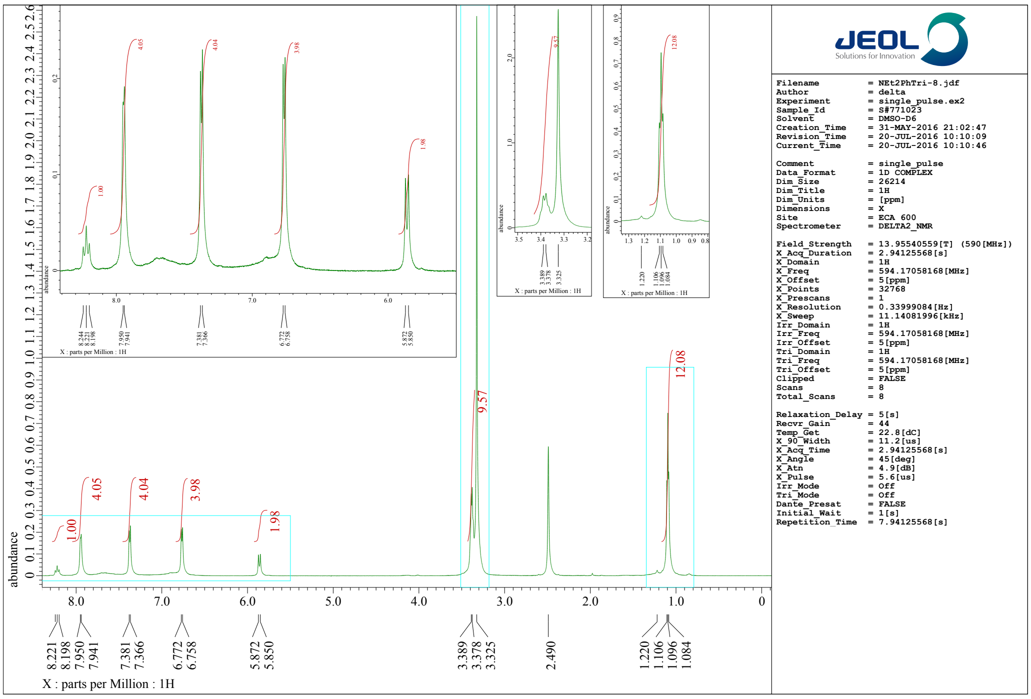


**Fig. S11-g7**. [1]H-NMR (100 MHz) spectrum of **PC7** in dimethyl sulfoxide-$d_6$ ($d$-DMSO).

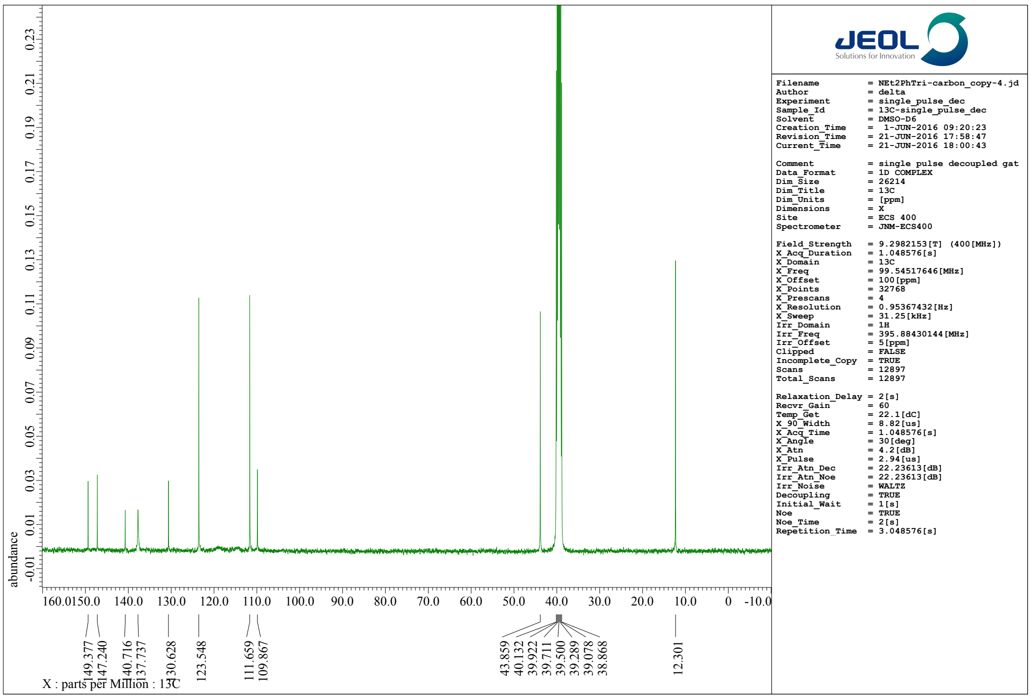


**Fig. S11-g8**. [13]C-NMR (100 MHz) spectrum of **PC7** in dimethyl sulfoxide-$d_6$ ($d$-DMSO).

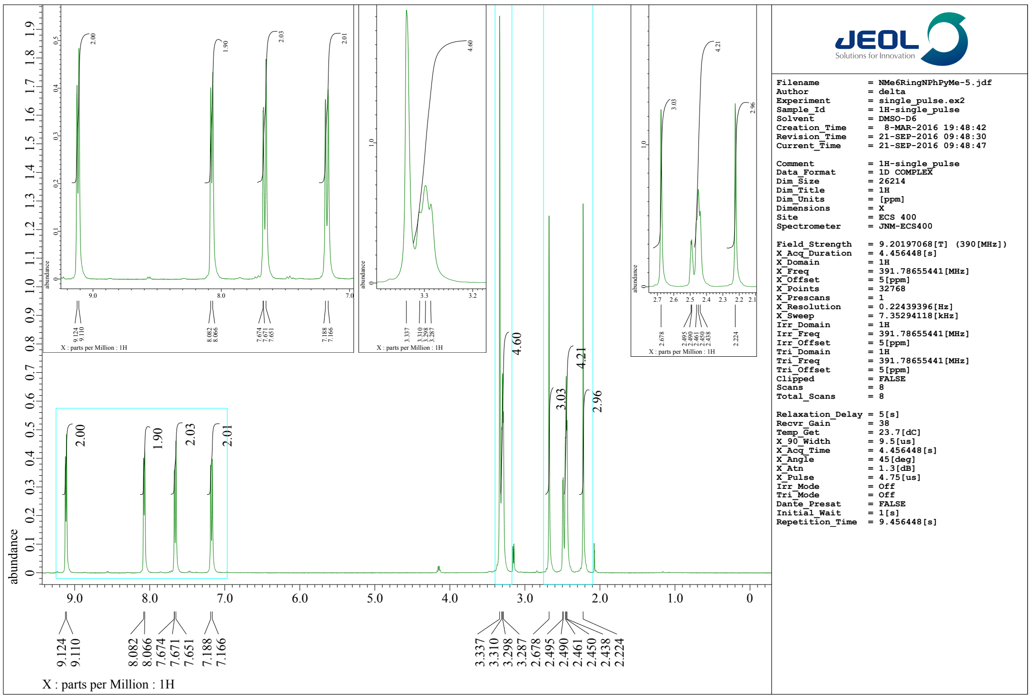


**Fig. S11-h1**. [1]H-NMR (400 MHz) spectrum of **8a** in dimethyl sulfoxide-$d_6$ (*d*-DMSO).

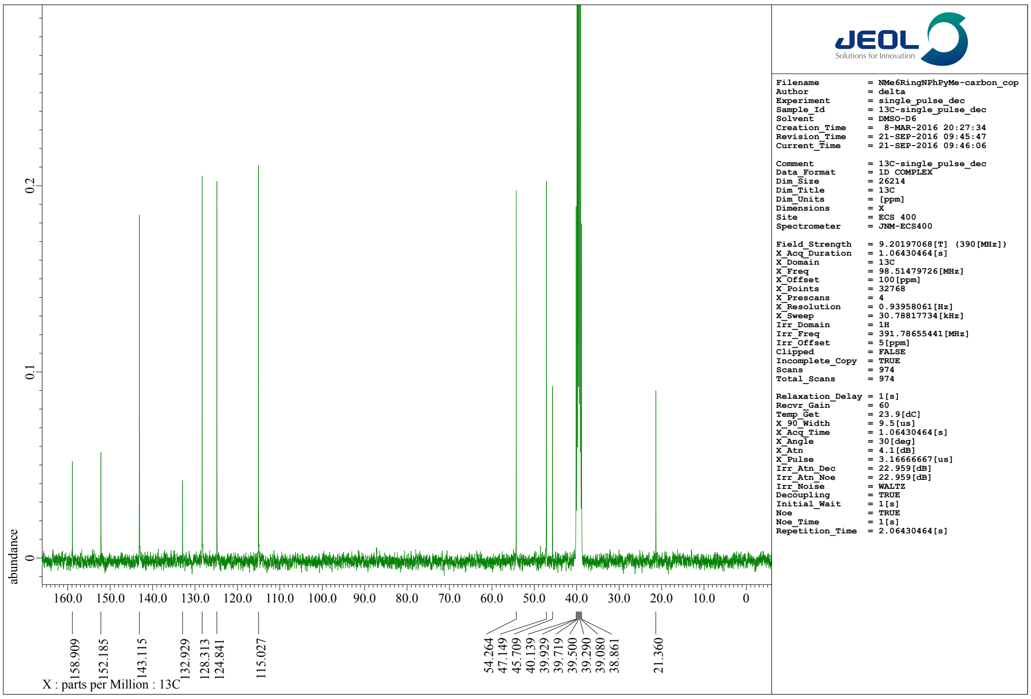


**Fig. S11-h2**. [13]C-NMR (100 MHz) spectrum of **8a** in dimethyl sulfoxide-$d_6$ (*d*-DMSO).

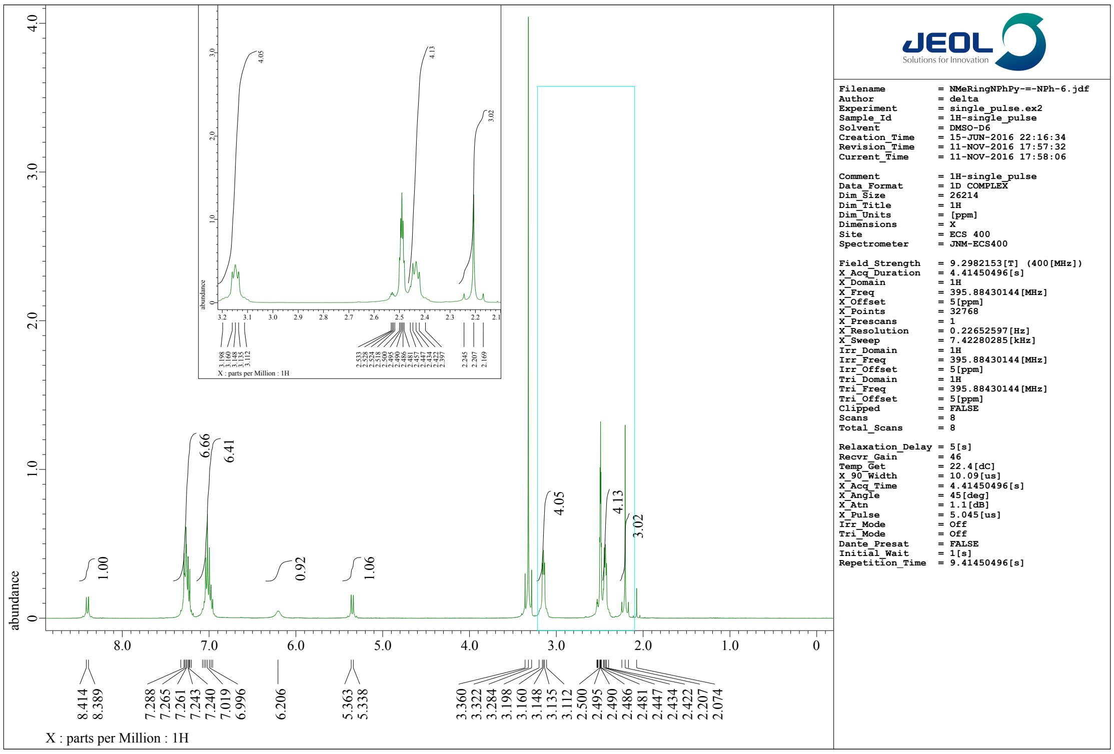


**Fig. S11-h3**. [1]H-NMR (400 MHz) spectrum of **8b** in dimethyl sulfoxide-$d_6$ (*d*-DMSO).

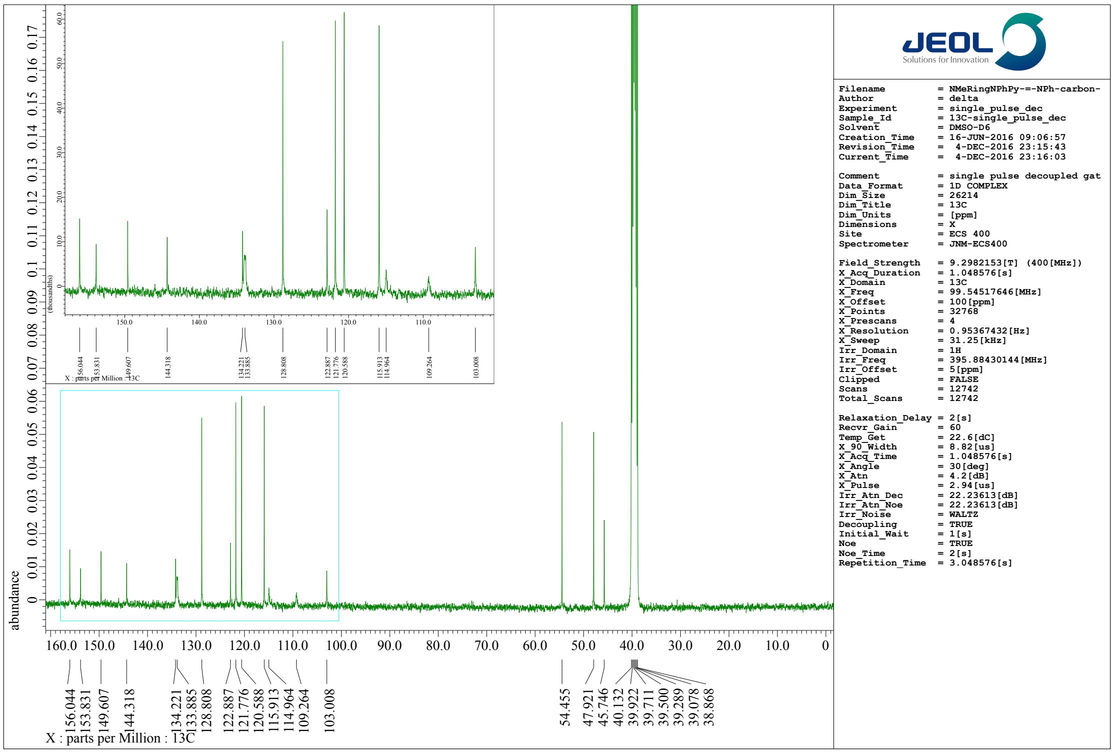


**Fig. S11-h4**. [13]C-NMR (100 MHz) spectrum of **8b** in dimethyl sulfoxide-$d_6$ (*d*-DMSO).

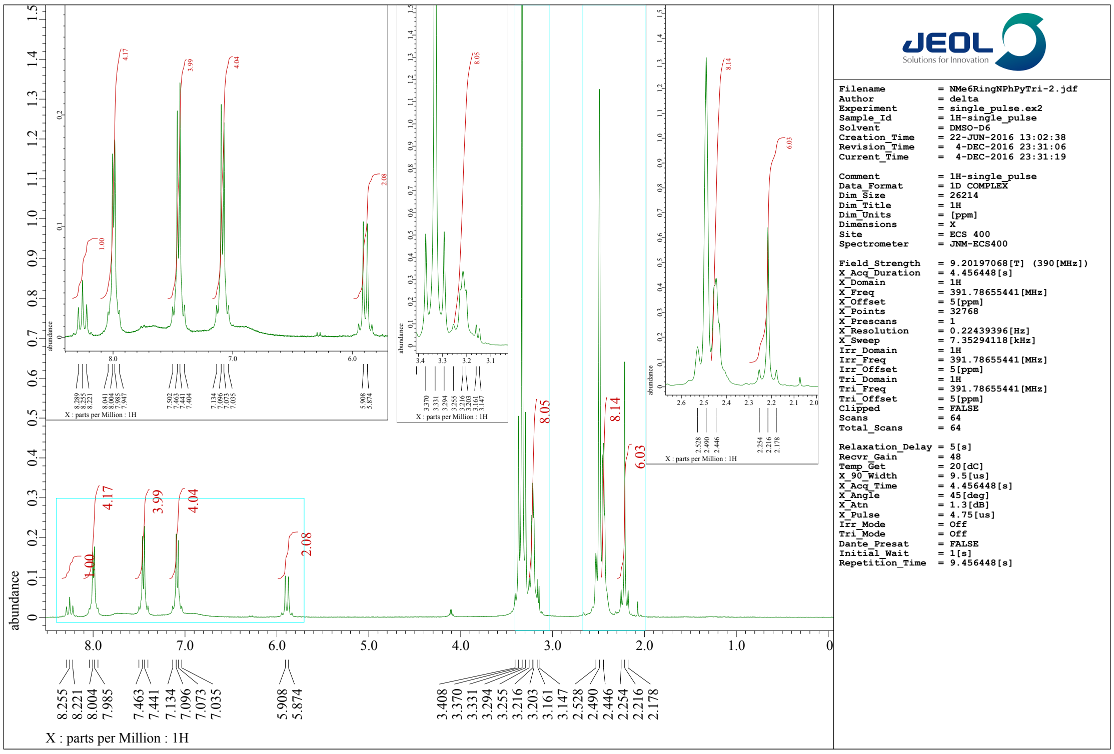


**Fig. S11-h5**. [1]H-NMR (400 MHz) spectrum of **PC8** in dimethyl sulfoxide-$d_6$ ($d$-DMSO).

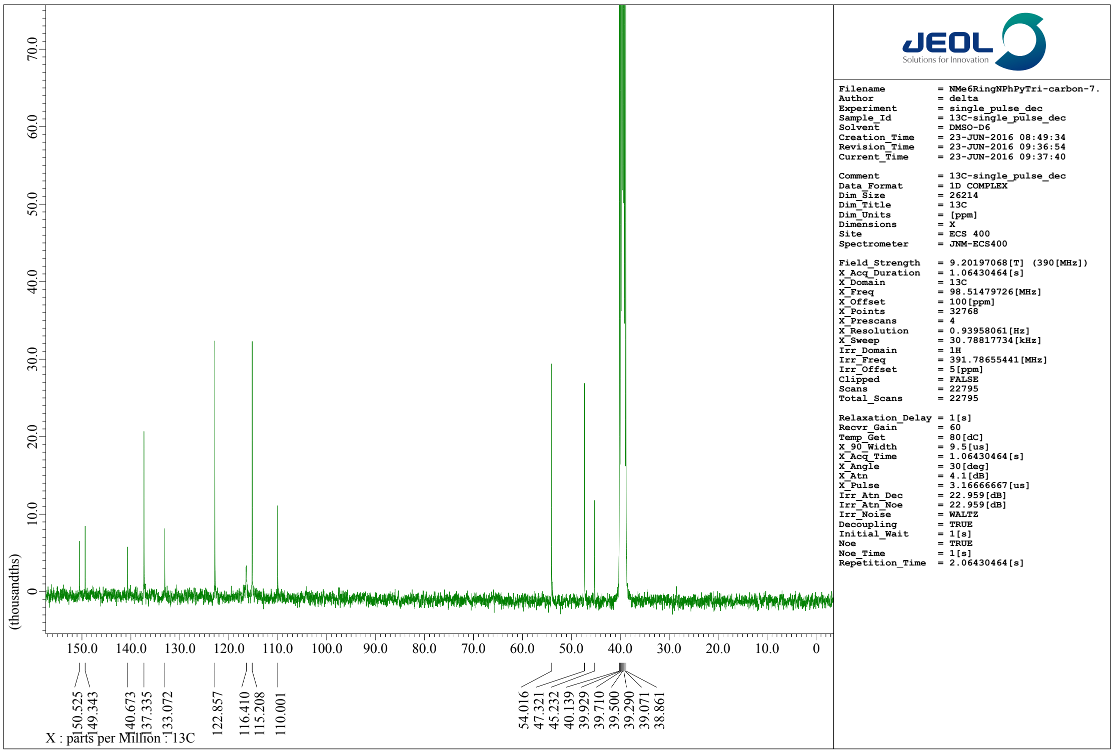


**Fig. S11-h6**. [13]C-NMR (100 MHz) spectrum of **PC8** in dimethyl sulfoxide-$d_6$ ($d$-DMSO).
